# Quantifying the impact of antibiotic use and genetic determinants of resistance on bacterial lineage dynamics

**DOI:** 10.1101/2025.02.03.636319

**Authors:** David Helekal, Tatum D. Mortimer, Aditi Mukherjee, Gabriella Gentile, Adriana Le Van, Robert A. Nicholas, Ann E. Jerse, Samantha G. Palace, Yonatan H. Grad

**Affiliations:** Department of Immunology and Infectious Diseases, Harvard T. H. Chan School of Public Health, Boston, MA 02115, USA; Department of Population Health, College of Veterinary Medicine, University of Georgia, Athens, GA 30602, USA; Department of Pharmacology, University of North Carolina at Chapel Hill, Chapel Hill, NC 27599, USA; Department of Microbiology and Immunology, Uniformed Services University of the Health Sciences, Bethesda, MD 20814, USA; Henry M. Jackson Foundation for the Advancement of Military Medicine, Inc, Bethesda, MD 20817, USA; Departments of Microbiology and Immunology, University of North Carolina at Chapel Hill, Chapel Hill, NC 27599, USA

## Abstract

The dynamics of antimicrobial resistance in bacterial populations are informed by the fitness impact of genetic determinants of resistance and antibiotic pressure. However, estimates of real-world fitness impact have been lacking. To address this gap, we developed a hierarchical Bayesian phylodynamic model to quantify contributions of resistance determinants to strain success in a 20-year collection of *Neisseria gonorrhoeae* isolates. Fitness contributions varied with antibiotic use, and genetic pathways to phenotypically identical resistance conferred distinct fitness effects. These findings were supported by *in vitro* and experimental infection competition. Quantifying these fitness contributions to lineage dynamics reveals opportunities for investigation into other genetic and environmental drivers of fitness. This work thus establishes a method for linking pathogen genomics and antibiotic use to define factors shaping ecological trends.

## INTRODUCTION

The prevalence of antimicrobial resistance (AMR) reflects competition in an ever-changing environment (*1*, *2*). When an antibiotic is introduced into clinical use, it can result in increased fitness and prevalence of resistant bacteria. Once resistance to the antibiotic becomes sufficiently widespread, its use is often reduced in favor of another antibiotic for which resistance prevalence is low. This alters the fitness landscape, such that alleles and genes that had conferred a fitness advantage in the context of the first antibiotic may become deleterious.

The complexity of the bacterial AMR fitness landscape is shaped by multiple factors. These include antibiotic pressure from drugs in current and past use as well as antibiotic resistance, which can often be achieved through multiple, at times interacting, genetic pathways (c.f., macrolide resistance in *N. gonorrhoeae* (*3*, *4*) and fluoroquinolone resistance in *E. coli* (*1*)). Population structure (*5*, *6*), linkage between AMR determinants in a changing environment (*1*, *6*), linkage with sites under balancing selection (*6*, *7*), mutation-selection balance (*8*), and non-antibiotic pressures can also inform this landscape.

While the genetic determinants of AMR play a large role in the population expansion and contraction of drug resistant lineages (*1*), the fitness impact of these determinants can vary across genetic backgrounds and environment (*9*, *10*). Efforts to quantify the fitness impact of individual genetic features across shifting patterns of antibiotic use in real world data have been fraught with many challenges, such as limited availability of data on antimicrobial use, both for targeted treatment and accounting for bystander exposure (*11*); the influence of other factors, such as pressure from host immunity, on overall pathogen fitness; and the frequent co-occurrence of multiple antibiotic resistance determinants in drug resistant strains (*5*, *6*). Additionally, for many pathogens, we lack longitudinal datasets of sufficient size, duration, and systematic collection to enable inference about fitness.

Here, we overcame these challenges to define the fitness contributions of AMR determinants in response to changes in treatment using data from *N. gonorrhoeae* in the USA. *N. gonorrhoeae* is an obligate human pathogen that causes the sexually transmitted infection gonorrhea; infection does not elicit a protective immune response (*12*, *13*). A collection of over 5000 specimens from 20 years (2000-2019) of the CDC’s Gonococcal Isolate Surveillance Project (GISP), the CDC’s sentinel surveillance program for antibiotic resistant gonorrhea, has been sequenced and has undergone resistance phenotyping, with metadata including the demographics of the infected individuals (*3*, *14–17*). Data on primary treatment in the US over this period have been reported by the CDC and reflect changes in first-line therapy, from fluoroquinolones to cephalosporins plus macrolides, and, among the cephalosporins, from the oral cefixime to intramuscular ceftriaxone (*18*). To combine sequencing data with resistance covariates, we used phylodynamic modeling. Building on prior efforts (*19–23*), we deployed a framework that can accommodate multiple lineages, multiple pathways, and multiple time-varying covariates to estimate quantitatively the resistance determinant-specific fitness costs in circulating *N. gonorrhoeae* and their interaction with antibiotic pressures.

## RESULTS

### Defining N. gonorrhoeae Drug Resistant Lineages

We first sought to define AMR-linked lineages from the GISP specimens. On epidemic timescales (*26*), *N. gonorrhoeae* maintains a lineage structure largely shaped by antimicrobials (*27*). The treatment for *N. gonorrhoeae* infections in the US over the past 20 years has been defined by three main drug classes: fluoroquinolones, third generation cephalosporins, and macrolides (figure S1). We used ancestral state reconstruction for the major AMR determinants for these antibiotics to identify clusters of specimens that had not changed state since descending from their most recent common ancestors (MRCA) <u>1.1</u>. We refined the classification by requiring that the MRCA was no earlier than 1980. As the three drug classes under study entered use after this date, this cutoff limits the analysis to a time frame over which ancestral state reconstruction is likely to remain accurate, while helping separate lineages that acquired the same resistance pattern independently. We focused on lineages that have at least 30 specimens, reasoning that a minimum cutoff helps avoid the inclusion of small outbreak clusters that could potentially produce unreliable estimates (see Materials and Methods: *Lineage Assignment & Phylogenetic Reconstruction*).

This definition led to the identification of 29 lineages across the dataset (figure 1), along with the corresponding distribution of determinants (table 1). The majority of lineages (21/29) have at least one AMR determinant, with multiple pathways to resistance for a given antibiotic present across lineages. Lineages 22 and 23, for example, carry the mosaic *penA* 34 allele, and the remaining 27 lineages all carry the Penicillin Binding Protein 2 (PBP2; encoded by *penA*) substitution *A*516*G*, each of which increases resistance to cephalosporins (*28*, *29*).

**Figure 1:**
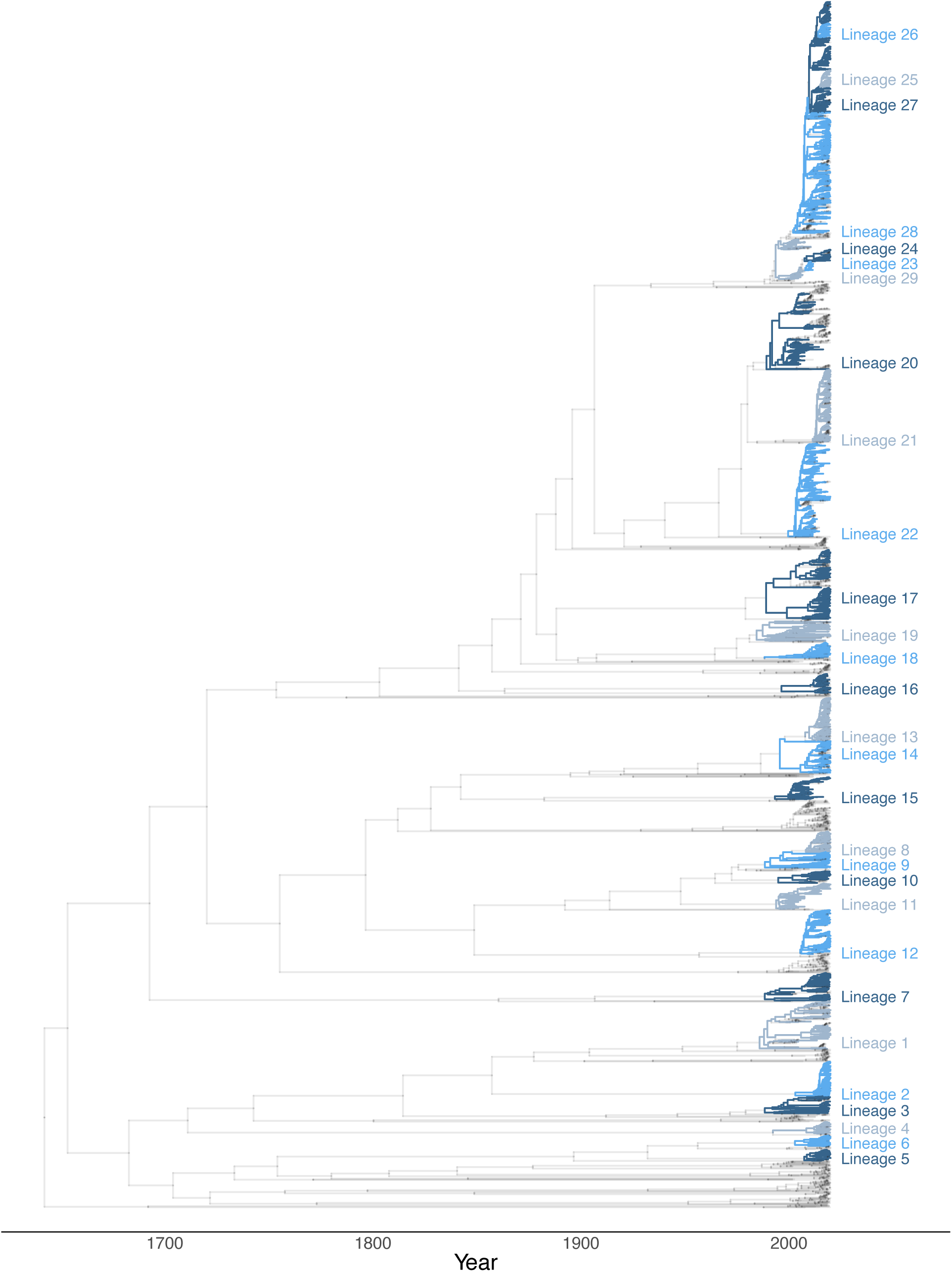
Lineage assignment based on AMR determinants. The phylogenetic tree is annotated according to the lineage assignment of the ancestral node. Gray nodes in the phylogenetic tree denote lines of descent that are not in an assigned lineage. Lineage numbering was determined by post-order traversal of the tree.

**Table 1:** Distribution of AMR determinants across lineages. Gray color corresponds to wild-type allele. For *gyrA*, *parC* and *ponA*, polymorphisms occur at several key amino acid positions. The *penA* and *mtr* loci exhibit complex patterns of polymorphism that include interspecies mosaicism as well as individual amino acid variations. non-m denotes non-mosaic alleles. Adel denotes A deletion in the *mtrR* promoter (*24*). LOF=loss of function. The only determinant in 23S rRNA that appears frequently in our dataset is the C2611T substitution, where T indicates at least one and up to four copies of 23S rRNA C2611T. Coloring highlights non-wild-type determinants in each column and changes from column to column.

| Lineage | *gyrA* |  | *parC* |  |  | *ponA* | *penA* |  |  | *mtr* |  |  |  | *23S rRNA* |
| --- | --- | --- | --- | --- | --- | --- | --- | --- | --- | --- | --- | --- | --- | --- |
|  | S91 | D95 | D86 | S87 | E91 | L421 | A501 | G542 | Type | *mtrC* | *mtrD* | *mtr* pro-moter | *mtrR* | C2611 |
| 1 | S | D | D | S | E | L | A | G | non-m | non-m | non-m | non-m | LOF | C |
| 2 | S | D | D | S | E | L | A | G | non-m | non-m | non-m | non-m | non-m | C |
| 3 | S | D | D | S | E | L | A | G | non-m | non-m | non-m | non-m | non-m | C |
| 4 | S | D | D | S | E | L | A | G | non-m | non-m | non-m | non-m | non-m | C |
| 5 | S | D | D | S | E | L | A | G | non-m | non-m | non-m | non-m | non-m | C |
| 6 | S | D | D | S | E | L | A | G | non-m | non-m | non-m | non-m | non-m | C |
| 7 | S | D | D | S | E | P | A | G | non-m | non-m | non-m | A-del | non-m | C |
| 8 | S | D | D | S | E | L | A | G | non-m | non-m | non-m | non-m | non-m | C |
| 9 | S | D | D | S | E | P | A | G | non-m | non-m | non-m | non-m | non-m | C |
| 10 | S | D | D | S | E | P | A | G | non-m | non-m | non-m | A-del | non-m | C |
| 11 | S | D | D | S | E | P | A | G | non-m | non-m | non-m | A-del | non-m | C |
| 12 | F | G | D | S | G | P | T | G | non-m | non-m | non-m | A-del | non-m | C |
| 13 | F | A | D | R | E | L | A | G | non-m | non-m | non-m | non-m | LOF | C |
| 14 | F | A | N | S | E | L | A | G | non-m | non-m | non-m | non-m | LOF | C |
| 15 | F | G | D | R | E | P | A | G | non-m | non-m | non-m | A-del | non-m | C |
| 16 | F | G | N | S | E | P | V | G | non-m | non-m | non-m | A-del | non-m | C |
| 17 | S | D | D | S | E | P | A | G | non-m | non-m | non-m | A-del | non-m | C |
| 18 | S | D | D | S | E | L | A | G | non-m | non-m | non-m | non-m | non-m | C |
| 19 | S | D | D | S | E | P | A | G | non-m | non-m | non-m | non-m | non-m | C |
| 20 | F | G | D | R | E | P | A | S | non-m | non-m | non-m | A-del | non-m | C |
| 21 | F | A | D | R | E | P | A | S | non-m | non-m | non-m | non-m | non-m | C |
| 22 | F | G | D | R | E | P | — | — | mosaic | non-m | non-m | A-del | non-m | C |
| 23 | S | D | D | S | E | L | — | — | mosaic | non-m | non-m | non-m | non-m | C |
| 24 | S | D | D | S | E | L | A | G | non-m | non-m | non-m | non-m | non-m | C |
| 25 | S | D | D | S | E | L | A | G | non-m | non-m | mosaic | mosaic | non-m | C |
| 26 | F | A | N | S | E | L | A | G | non-m | mosaic | mosaic | mosaic | non-m | C |
| 27 | S | D | D | S | E | L | A | G | non-m | mosaic | mosaic | mosaic | non-m | C |
| 28 | S | D | D | S | E | L | A | G | non-m | non-m | mosaic | non-m | non-m | C |
| 29 | S | D | D | S | E | L | A | G | non-m | non-m | non-m | non-m | non-m | T |

The related lineages 20 – 22 share many resistance determinants but have differing estimates of their effective population sizes through time, *Ne*(*t*), and illustrate the dynamics that emerge when juxtaposing *Ne*(*t*) with antibiotic use and resistance (figure 2). This cluster of lineages contains a previously described and the largest mosaic *penA* 34-carrying lineage, lineage 22 (*3*), along with its two sister-lineages. Lineage 20, the oldest lineage in this cluster, grew during the fluoroquinolone era and decreased afterwards (*22*). In this lineage, nearly all descendants sampled after the recommendation of ceftriaxone plus azithromycin dual therapy had acquired a new resistance determinant or lost an existing one. In one sublineage, this change included replacement of a resistance-conferring *gyrA* allele (encoding 91F, 95G) with the wild-type allele (91S, 95D), resulting in phenotypic susceptibility (figure 2, blue bar). Another sublineage changed *mtrR* promoter alleles (figure 2, yellow bar). Determinants at the *mtrR* promoter are associated with resistance to a wide range of antibiotics (*24*) including macrolides (*30*). Yet another sublineage acquired azithromycin resistance through C2611T substitution in 23S rRNA (figure 2, red bar). Furthermore, the descendants of Lineage 20 appeared to switch sexual networks: most recent isolates were from heterosexuals whereas past isolates were from men who have sex with men (figure S2). Lineage 21 expanded after the 2010 switch in recommended treatment to dual therapy with azithromycin plus ceftriaxone. The effective population size for the mosaic *penA* 34-carrying lineage 22 grew during the fluoroquinolone period and after, but decreased with the introduction of azithromycin and ceftriaxone dual therapy. Together, these patterns of lineage expansion and contraction indicated a relationship among the antibiotics recommended for treatment, genetic determinants of resistance, and lineage success. However, while for lineages 20 and 22 the pattern of expansion and contraction matches our expectations based on their resistance profile, it was not clear from a simple inspection what could explain the dynamics of lineage 21.

**Figure 2:**
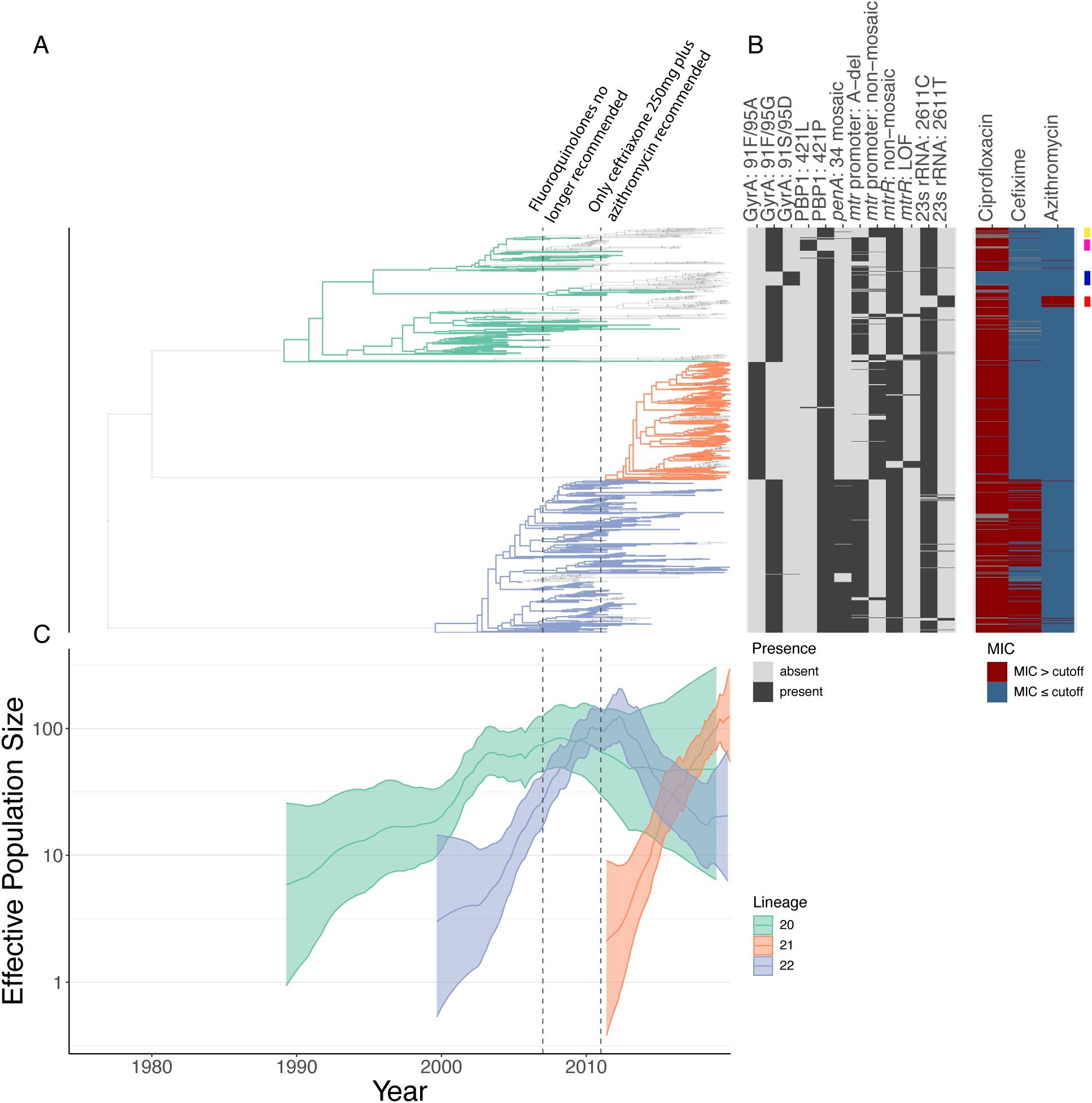
A cluster of phylogenetically related lineages with evidence of adaptation in response to changes in antibiotic use. Panel A: the phylogeny for lineages 20-22. The gray transparent tips correspond to isolates that have diverged from the ancestral determinant combination of the parental lineage. Panel B: the presence and absence of relevant determinants and antibiotic resistance phenotypes above or below each drug’s cutoff (CIP: 1*μ*g/mL; CFX: 0.25*μ*g/mL, AZI: 4*μ*g/mL). The yellow bar on the right highlights a cluster of isolates that changed *mtr* promoter alleles. The magenta bar highlights a reversion of resistance conferring PBP1 421P mutation to susceptible 421L. The blue bar on the right highlights a cluster of isolates that reverted to a fluoroquinolone-susceptible *gyrA* allele. The red bar on the right highlights a cluster of isolates that acquired azithromycin resistance. Panel C: Median effective population size trajectories for each of the lineages with 95% credible intervals as estimated by PHYLODYN (*25*).

### Hierarchical Bayesian Phylodynamic Modeling Reveals a Changing Fitness Landscape

We next sought to quantify the fitness contributions of the genetic determinants of resistance and how these varied over time. For each determinant, we estimated a set of regression coefficients, one for each of the antibiotic classes to which it conferred resistance, along with an intercept term. These modeled the effect of a given resistance determinant on the effective population size growth rate of lineages that carry that determinant as a function of the reported treatments (table S1). To account for the impact of lineage background, and over-dispersion of lineage growth rates due to effects not included in the model, we also included lineage-specific residual terms, lineage-specific background terms, and global mean growth-rate term in a hierarchical manner. We noted that as the treatment data are percentages summing to 1 and thus are not full rank, we selected ceftriaxone 250mg as the baseline for all estimated treatment use effects. Full details are available in Materials and Methods. For a high-level description and rationale behind the model, see (Materials and Methods: *Lineage-Based Hierarchical Phylodynamic Model – Overview and Rationale*). For a detailed technical description see (Materials and Methods: *Lineage-Based Hierarchical Phylodynamic Model – Construction of the Likelihood* and *Lineage-Based Hierarchical Phylodynamic Model – Detailed Description of the Statistical Model*). For implementation details see (Materials and Methods: *Lineage-Based Hierarchical Phylodynamic Model – Implementation)*.

This model formulation allowed us to answer three main questions. First, did the relative fitness of a given resistance determinant change as a function of the pattern of antibiotic use? Second, what is the fitness cost or benefit through time associated with a particular resistance allele compared to its susceptible counterpart? Third, how much of a lineage’s trajectory is explained by the fitness contributions of the resistance determinants? To answer these questions, we first calculated the predicted effect of individual determinants on the growth rate of lineages, finding that several resistance determinants had a strong impact (defined by the 95% posterior credible interval interval excluding [–0.1,0.1]) on lineage dynamics (table S2).

#### gyrA

*gyrA* is the main fluoroquinolone-resistance determining gene in *N. gonorrhoeae*, with alleles of *parC* also contributing to resistance (*30*). Our modeling revealed several phenomena among lineages encoding ParC 86D/87R/91E. First, lineages carrying GyrA 91F/95G with this *parC* allele experienced a growth rate increase during the period of recommended fluoroquinolone treatment for gonorrhea (figure 3**A**, figure S3). However, for GyrA 91F/95G, there was too much uncertainty to determine its absolute effect on the growth rate of lineages carrying it during the fluoroquinolone period compared to the baseline GyrA 91S/95D type within the wild-type ParC 86D/87S/91E context (figure 3**A**). After fluoroquinolones were no longer recommended, GyrA 91F/95G appeared weakly deleterious when combined with ParC 86D/87R/91E allele compared to wild-type (figure 3**A**, table S2). Second, lineages carrying GyrA 91F/95A had distinctly higher growth rates than the GyrA 91S/95D susceptible allele after the period in which fluoroquinolones were used for treatment (figure 3**A**, table S2). Third, lineages carrying GyrA 91F/95A had a relative growth rate advantage over GyrA 91F/95G after the end of fluoroquinolone era (figure S4). The majority of lineages carrying GyrA 91F/95A expanded only after 2007 when fluoroquinolones were no longer recommended, increasing the uncertainty in estimates of the effect that fluoroquinolone use had on these lineages.

**Figure 3:**
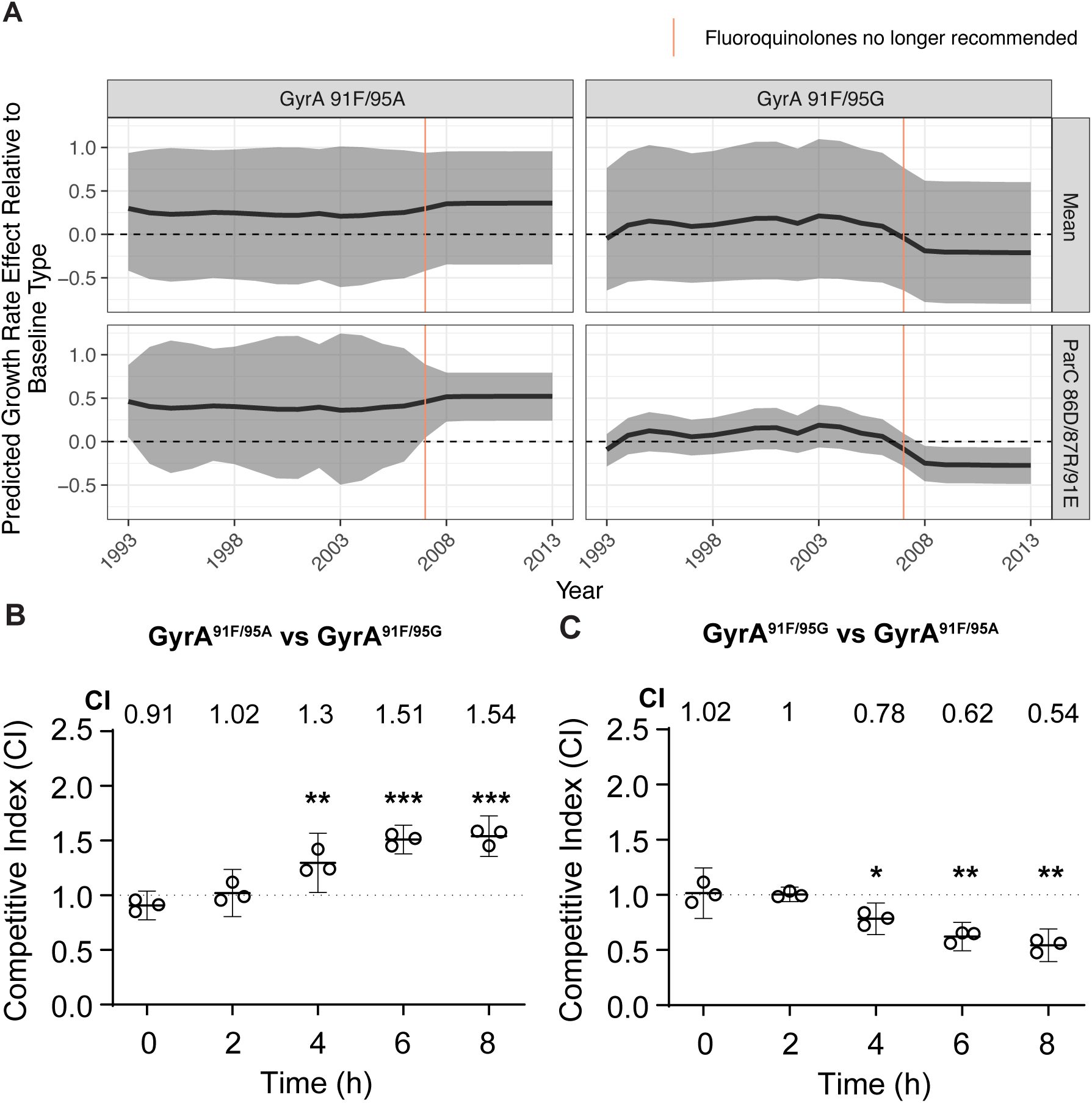
The summary of results for fitness impact of selected GyrA determinants. Panel **A**: The predicted absolute effect on lineage growth rate of selected GyrA determinants across all *parC* contexts (top, “Mean”) or within the ParC 86D/87R/91E context (bottom) based on past fluoroquinolone use patterns. The predicted effect is an absolute effect as computed compared to the baseline GyrA 91S/95D type within the wild-type ParC 86D/87S/91E context. The shaded region denotes the 95% posterior credible interval around the posterior median, depicted by the bold black line. Dashed line denotes no predicted growth rate effect relative to baseline allele. Panel **B** and **C**: *In vitro* competition in liquid culture between (**B**) unlabeled GCGS0481 GyrA 91F/95G and kanamycin-labeled GyrA 91F/95A; and (**C**) unlabeled GCGS0481 GyrA 91F/95A and kanamycin-labeled GyrA 91F/95G. The competitive index (CI) is shown for the kanamycin-labeled strain relative to the unlabeled strain. Statistical significance (at 2, 4, 6 and 8 hours: (**B**) p = 0.13, 0.005, 0.0002, 0.0003; (**C**) p = 0.84, 0.02, 0.003, 0.0017). N = 3/time point, representative of three independent experiments. Error bars represent mean with 95% CI. Statistical significance of CI measurements was assessed using an unpaired two-sided Student’s *t-test* (\**p* < 0.05, \*\**p* < 0.005 and \*\*\**p* < 0.0005).

To investigate whether the degree of resistance provided by GyrA 91F/95G differed from GyrA 91F/95A in lineages containing the ParC 86D/87R/91E allele, we fitted a linear model to log_2_-transformed ciprofloxacin MIC, while accounting for determinants at the *mtrCDE* operon. There was no significant difference in log_2_-transformed ciprofloxacin MICs (GyrA 91F/95G coefficient = 0.183, two-sided T-test P-value > 0.242; see for details).

Given these results, we tested whether GyrA 91F/95A contributes to a growth rate advantage over GyrA 91F/95G *in vitro*. Both sets of mutations increased the ciprofloxacin MIC ≥128-fold over the susceptible *gyrA* allele (table S3). In an *in vitro* competition assay between GyrA 91F/95A and GyrA 91F/95G isogenic strains in the GCGS0481 strain background with ParC 86D/87R/91E, GyrA 91F/95A conferred a fitness benefit (figure 3**B**, figure 3**C**): the competitive index (CI) after 8 hours of competition for GCGS0481 kanamycin-labeled GyrA 91F/95A versus GyrA 91F/95G was 1.54 (p = 0.0003), consistent with the reciprocal competition, in which the CI of kanamycin-labeled GyrA 95F/95G versus GyrA 91F/95A was 0.54 (p = 0.0017). Both GyrA 91F/95G and GyrA 91F/95A strains were less fit than the susceptible parental strain (figure S5). After 8 hours of competition, the CI of kanamycin-labeled GyrA 91F/95A versus GyrA 91S/95D was 0.67 (p = 0.0015), and for kanamycin-labeled GyrA 91F/95G versus GyrA 91S/95D, it was 0.58 (p = 0.0009) (figure S5). Consistent with this, the CI after 8 hours of competition of GCGS0481 kanamycin-labeled GyrA 91S/95D versus GyrA 91F/95A was 1.45 (p = 0.0023) and kanamycin-labeled GyrA 91S/95D versus GyrA 91F/95G was 1.72 (p = 0.0001) (figure S5). See Materials and Methods for details of the *gyrA* competition experiments.

GyrA 91F/95A within the ParC 86N/87S/91E context conferred a growth rate advantage after 2007, when fluoroquinolones were no longer in use, compared to the baseline type that does not carry any of the resistance determinants studied (table S2, figure S6).

#### penA

Variants in the *penA* gene, which encodes PBP2, contribute to resistance to cephalosporins as well as other beta lactams, with mosaic *penA* alleles the major determinants of resistance to cephalosporins (*14*, *31*, *32*). Most of the cephalosporin resistance determinants in our dataset appeared in 1-3 lineages each (table 1), which limited our ability to estimate their impact on growth rates (table S2, figure S7). However, we estimated a major beneficial effect of mosaic *penA* 34 on growth rates when cefixime and ceftriaxone 125mg were widely used, as well as the subsequent loss of this beneficial effect when treatment with cephalosporins other than ceftriaxone 250mg declined. (figure 4, table S2). Similarly, carriage of PBP2 501T was associated with a large relative decrease in fitness when ceftriaxone 250mg became the sole recommended treatment (table S2, figure S7). However, the absolute effect for PBP2 501T compared to wild-type cannot be identified (table S2, figure S7), as PBP2 501T appeared only in a single lineage that carries a unique GyrA/ParC combination. The decrease in the predicted growth rate effect for both mosaic *penA* 34 and PBP2 501T started in 2008 and aligns with a shift in primary treatment with cephalosporins to ceftriaxone 250mg, even before the guidelines changed in 2012 (figure S1).

**Figure 4:**
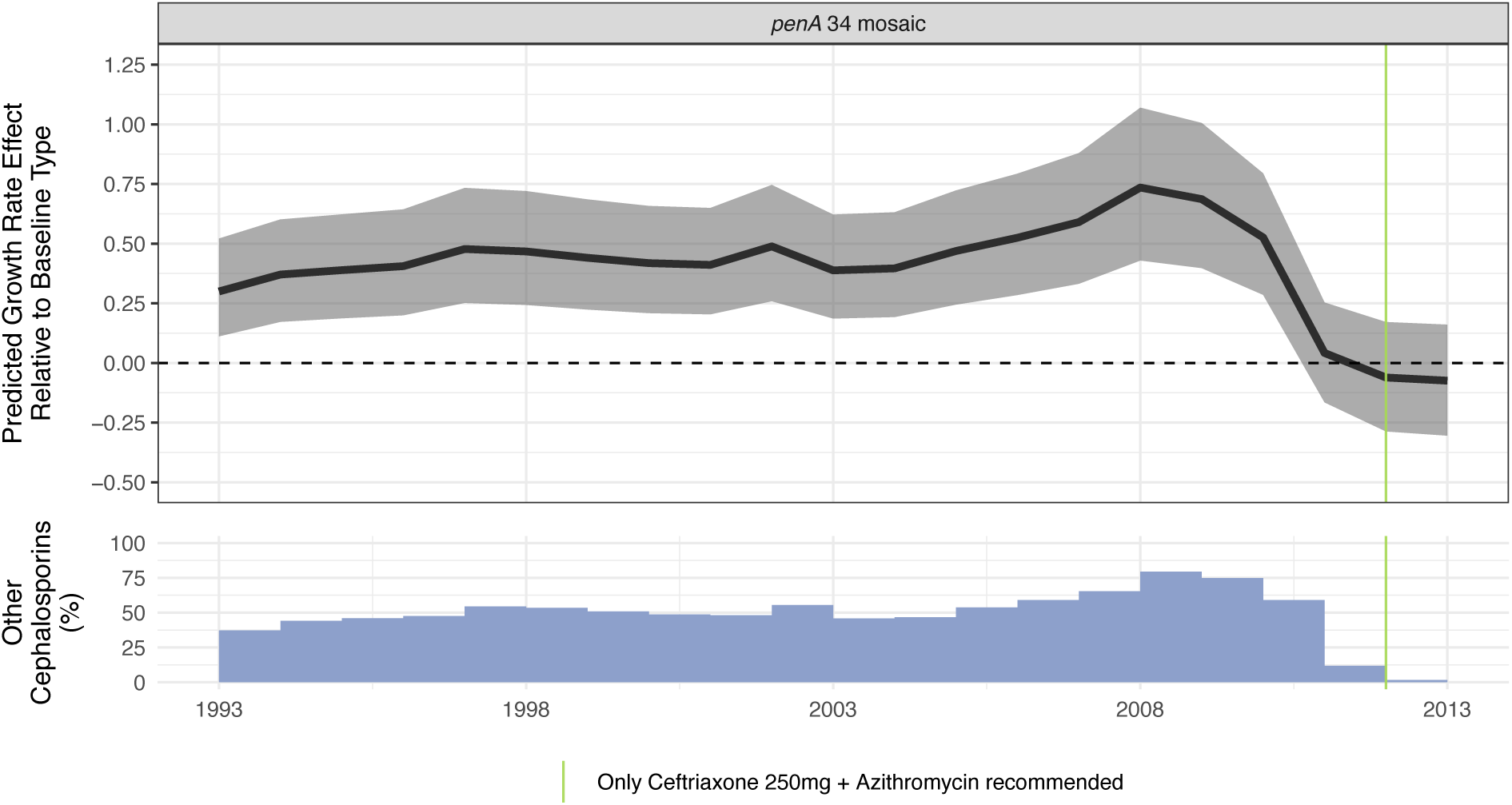
Summary of results for mosaic *penA* 34. Top Panel: The predicted absolute growth rate effect for mosaic *penA* 34 compared to baseline. The predicted effect was computed based on reported treatments. The shaded region denotes the 95% credible interval around the posterior median, depicted by the bold black line. Dashed line denotes no predicted growth rate effect relative to the baseline type that does not carry any of the resistance determinants studied. Bottom Panel: The use of cephalosporins other than ceftriaxone 250mg as a percentage of primary treatment. Other cephalosporins consist mainly of ceftriaxone 125mg, cefixime and other unclassified cephalosporins.

PBP2 501V was associated with a weak increase in fitness after the switch in treatment to ceftriaxone 250mg (table S2, figure S8). Both the PBP2 501V and 501T have wide credible intervals for their absolute effects compared to the baseline type that does not carry any of the resistance determinants studied (table S2, figure S7). These alleles occur in a single lineage each, with both lineages carrying unique *gyrA*/*parC* combinations (table 1), making the intercept term for these determinants unidentifiable.

#### ponA

The *ponA* gene encodes Penicillin Binding Protein 1 (PBP1). Variants in *ponA* contribute to resistance to penicillin (*33*), while also being associated with an increase in cephalosporin MICs (*28*).

The PBP1 421P variant was associated with a weak disadvantage compared to the baseline type that does not carry any of the resistance determinants throughout the study period (figure 5**A**, table S2). However, there was no discernible change to its fitness effect as the antibiotic treatment composition changed, (table S2, figure S11), consistent with PBP1 421P being primarily associated with an increase in penicillin MICs and only a mild increase in cephalosporin MICs (*28*). This is consistent with the observation of reversions from the resistance-associated PBP1 421P to the wild-type PBP1 421L. To test if the PBP1 421P confers a fitness cost in the absence of antibiotic treatment, we competed the wild-type PBP1 421L against PBP1 421P in two genetic backgrounds: the penicillin susceptible FA19 strain that normally carries the wild-type PBP1 421L, and the penicillin resistant FA6140 strain (*34*) that normally carries PBP1 421P. While the PBP1 421L and PBP1 421P derivatives of each strain had similar fitness during *in vitro* growth (Figure S), the PBP1 421P allele incurred a fitness cost to both FA19 and FA6140 when in competition with the isogenic PBP 421L strain in a female murine infection model: 5 days post infection, the competitive index for the FA19 *rpsL* PBP1^421P^ versus FA19 *rpsL* PBP1^421L^ mutant was 0.02 (p=0.0006) (figure 5**B**), and the competitive index for FA6140 *rpsL* PBP1^421L^ vs FA6140 *rpsL* PBP1^421P^ was 3.46 (p=0.03) (figure S9). See Materials and Methods for details of the *ponA* competition experiments.

**Figure 5:**
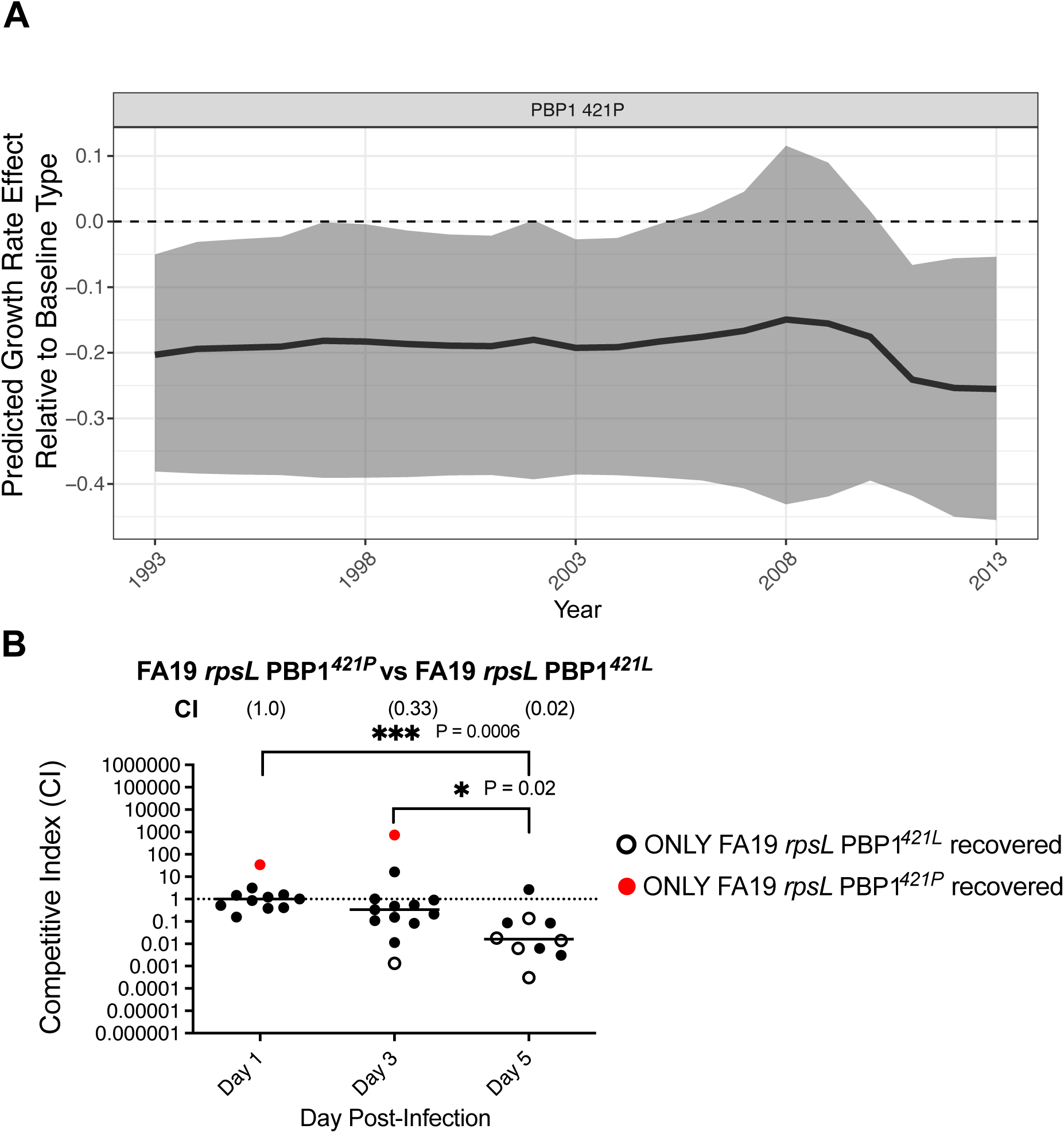
The summary of results for fitness impact of PBP1 421P. Panel **A**: The predicted absolute growth rate effect for PBP1 421P on lineage growth rate. The predicted effect was computed based on reported treatments. Shaded region denotes the 95% posterior credible interval around the median, depicted by the bold black line. Dashed line denotes no predicted growth rate effect relative to the baseline type that does not carry any of the resistance determinants studied. Panel **B**: *in vivo* murine competition assay between FA19 *rpsL* PBP1^421P^ vs FA19 *rpsL* PBP1^421L^. Competitive index (CI) was measured 1, 3, and 5 days post-infection. Where only one strain was detected, the CI is imputed at the limit of detection (see Materials and Methods). N = 2×7/time point, equating to two independent experiments. Each point represents the CI measurement for an individual mouse. The horizontal lines indicate the geometric mean. Statistical significance of CI measurements was assessed using the Mann-Whitney test (\**p* < 0.05, \*\**p* < 0.005 and \*\*\**p* < 0.0005).

#### *mtr* locus

*mtrCDE* encodes an efflux pump that modulates resistance to a wide range of antibiotics in *N. gonorrhoeae* (*24*), including to macrolides (*30*), and it is regulated by its transcriptional repressor, MtrR. Of particular relevance are mosaic *mtrC*, *mtrD*, and *mtrR* promoter as these are associated with azithromycin resistance (*3*, *35*). While our modeling recovered a growth rate increase associated with the carriage of mosaic *mtrR* promoter and the mosaic *mtrD* compared to wild-type baseline during the azithromycin co-treatment era (table S2, figure S12), the fact that mosaic *mtrR* promoter only occurred on mosaic *mtrD* backgrounds in our dataset (table 1) raises the concern that the estimated effects of the *mtrR* promoter and the mosaic *mtrD* maybe be only weakly identified. As such, we focused on the combined effect of mosaic *mtrR* promoter, mosaic *mtrC*, and mosaic *mtrD*. The predicted combined effect had large uncertainty, limiting interpretation, with clear support for a growth rate benefit only in 2010-2011 (figure 6).

**Figure 6:**
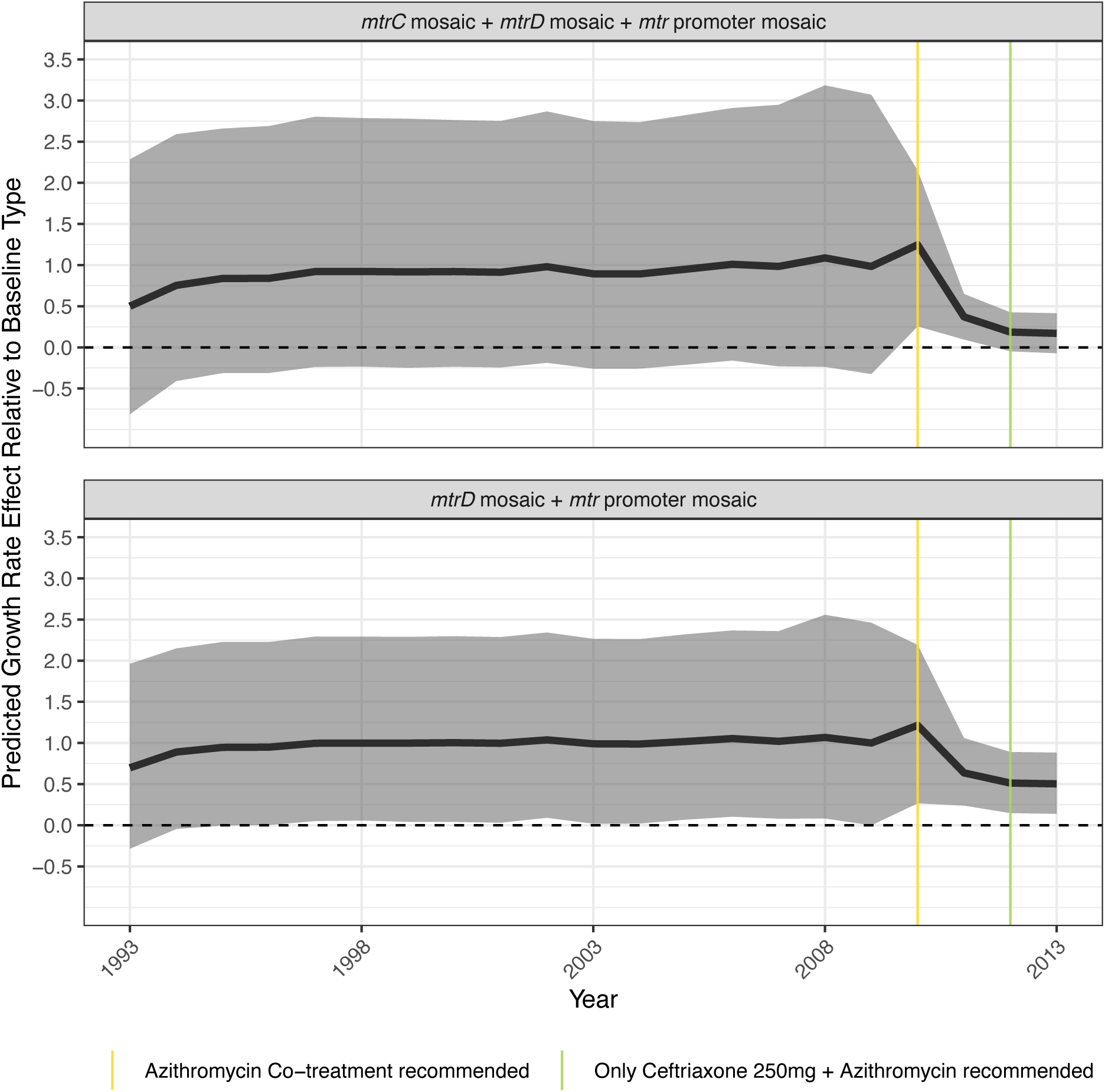
Predicted absolute effect for the total impact of the two most common combinations of determinants at the *mtrCDE* locus. The predicted effect was computed based on reported treatments. The shaded region denotes the 95% credible interval around the posterior median, depicted by the bold black line. Dashed line denotes no predicted growth rate effect relative to the baseline type that does not carry any of the resistance determinants studied.

The *mtrR* promoter A deletion in the 13-bp inverted repeat – a determinant implicated in an increase in resistance to a wide range of antibiotics including macrolides (*24*) – was associated with a weak increase in growth rate after the switch to ceftriaxone 250mg and azithromycin. This increase led to a weak advantage in growth rate after 2012 compared to the baseline type that does not carry any of the resistance determinants studied (table S2, figure S12, figure S13).

### Extent of lineage growth trajectory explained by resistance determinants

To quantify the extent to which the set of resistance determinants and lineage background explains each lineage’s growth rate over time, for each lineage we visualized the average growth rate effect of individual resistance determinants, along with lineage residual effect and lineage background effect, and summarized the total effect in the four treatment recommendation periods in the study period (figure 7, figures S14–S41).

**Figure 7:**
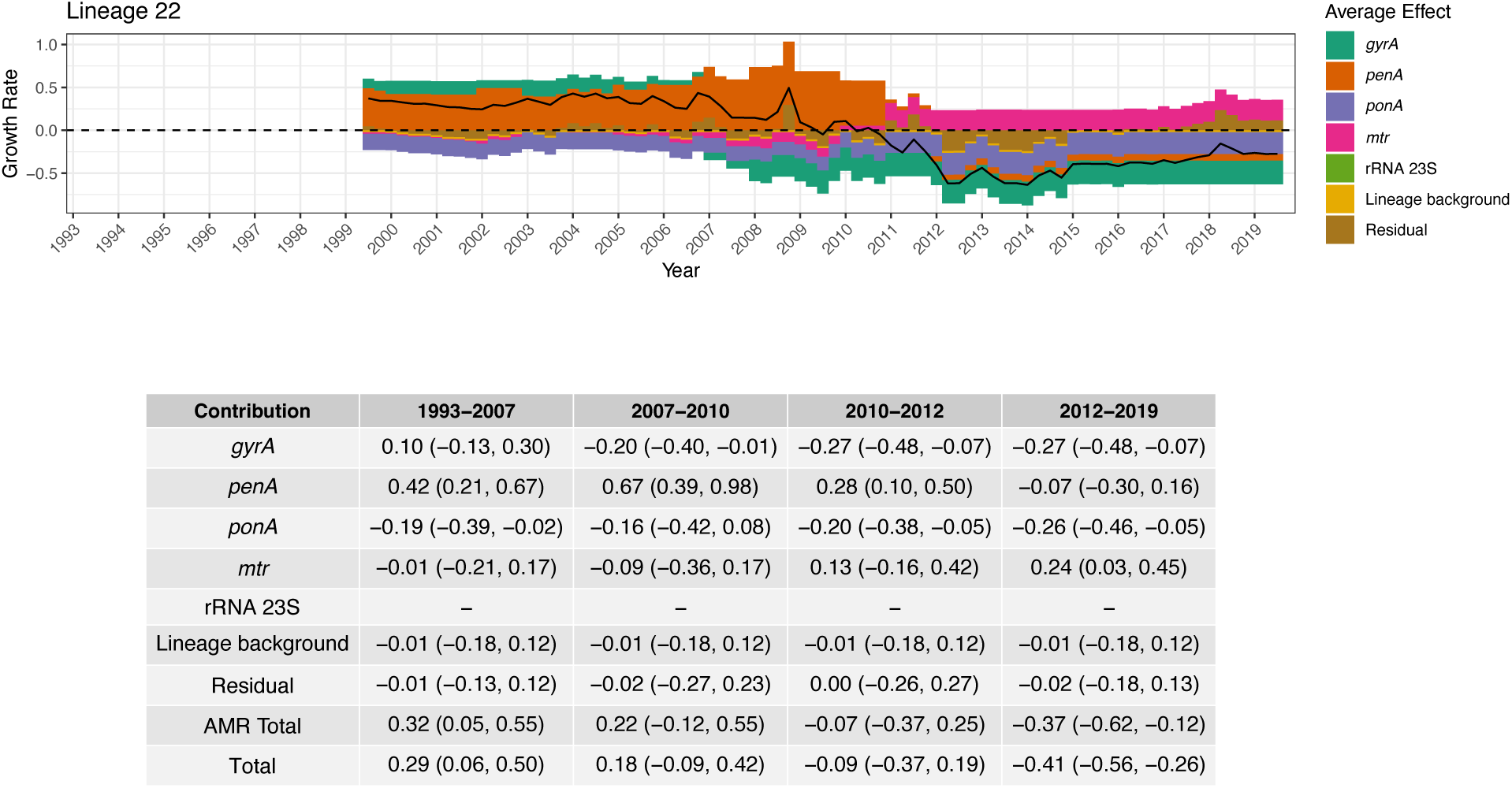
Growth rate effect summary for Lineage 22. The top panel shows the combined average growth rate effect of resistance determinants along with the lineage background term and the residual. The black solid line represents the total average effect. The dashed horizontal line indicates zero. The bottom panel depicts a table summarizing the median total growth rate effect across 4 treatment periods, as well as the 95% credible interval around the median in brackets. The period 1993-2007 corresponds to when fluoroquinolones were recommended as primary treatment; 2007-2010 to when multiple cephalosporins were recommended; 2010-2012 to when multiple cephalosporins were recommended along with azithromycin co-treatment; and 2012-2019 to when only ceftriaxone 250mg along with azithromycin co-treatment was recommended.

This per-lineage analysis revealed the shifting contributions of resistance determinants to individual lineage growth dynamics, provided examples in which fitness costs of one determinant are counterbalanced by the fitness benefits of another, and identified lineages with dynamics unexplained by these determinants. Lineage 22 carried a combination of GyrA 91F/95G along with mosaic *penA* 34. Despite carrying GyrA 91F/95G, which was associated with a fitness cost after fluoroquinolones were no longer recommended in 2007 (figure 3**A**, table S2), the growth of this lineage peaked in 2010, reflecting the fitness benefit of the mosaic *penA* 34 (figure 7). The shift in use to cephalosporins plus azithromycin in 2010 was accompanied by a fitness benefit from *mtr* variants, though cumulatively the fitness costs from other resistance determinants

The per-lineage analysis also addressed the question of what drove the growth of lineage 21, indicating that its expansion post-2010 was driven primarily by the presence of GyrA 91F/95A (figure S34).

While the fitness contributions of the set of resistance determinants in our model accounted for much of the lineage dynamics, some dynamics remained unexplained. For each lineage, we computed the number of years in which the absolute value of the average of the sum of the residual and lineage background terms exceeded the threshold of 0.1, representing approximately 10% growth or decline in a given year. In 11/29 lineages (lineages 2-4, 11, 13, 15, 16, 23, 26-28), there was at least one such year, and in six of those (lineages 2-4, 11, 15, 27), there were at least 3 such years.

To investigate this pattern, we examined lineages 2, 3, and 4. Lineage 2 carried none of the determinants we included in our model and underwent substantial growth starting in 2012 and peaking in 2018. We revisited the resistance phenotypes and genotypes for this lineage and noted that the isolates in Lineage 2 carry *tetM*, which confers high-level resistance to tetracycline-class antibiotics (*36*). We did not include *tetM* in our model because we lacked data on the extent of tetracycline-class antibiotics use for *N. gonorrhoeae* treatment and for syndromic treatment of known or presumed chlamydial co-infection and because none of the other lineages carry *tetM*. Lineages 3 and 4 had high residual effects for at least 3 years and carried none of the resistance determinants in our model. The large and consistently positive residual effects (figure S16 and figure S17) thus point to factors other than the antibiotic pressures examined here in shaping these two lineages’ success.

## DISCUSSION

Antibiotic exposure selects for resistant strains over their susceptible counterparts, whereas in the absence of antibiotics the resistant strains may suffer a fitness cost. While this relationship plays a central role in shaping microbial population dynamics, we have lacked quantitative estimates of the environmentally varying fitness effects of genetic elements in natural populations. Here, we used *N. gonorrhoeae* population genomics from large-scale surveillance data, detailed understanding of the genetics underlying antibiotic resistance, and data on antibiotic treatment to quantify the contribution of and the interactions among antibiotic resistance determinants and how these shaped *N. gonorrhoeae* AMR dynamics in the US over the study period (1993-2019).

Models of antibiotic use-resistance relationships typically treat all phenotypically resistant strains as if they have the same fitness costs and benefits (*37*). However, our findings suggest that a single amino acid difference in a resistance determinant may result in markedly different dynamics. The fluoroquinolone resistance-conferring alleles of GyrA 91F/95G and 91F/95A have phenotypically similar levels of resistance in the context of ParC 86D/87R/91E; however, after fluoroquinolones were no longer recommended, GyrA 91F/95G was associated with a fitness cost whereas GyrA 91F/95A was associated with a benefit. In line with this, the rising prevalence of fluoroquinolone resistance in *N. gonorrhoeae* (figure S1) masked the replacement of lineages carrying GyrA 91F/95G with those carrying GyrA 91F/95A. To help distinguish whether this advantage was due to GyrA 91F/95A itself or a tightly linked variant, *in vitro* competition assays demonstrated that the GyrA 91F/95A-containing strain was more fit than an isogenic GyrA 91F/95G strain, supporting the hypothesis that single amino acid differences in resistance determinants can drive distinct evolutionary trajectories. Several potential explanations exist for this phenomenon. One possibility is that reduced fitness cost of GyrA 91F/95A facilitates an overall fitness benefit in the presence of bystander exposure to fluoroquinolones, whereas GyrA 91F/95G is simply too costly to provide a net benefit from bystander exposure to fluoroquinolone alone, in the absence of direct use. Another possibility is that there may be a fitness benefit irrespective of treatment exposure *in vivo*. Studies have characterized an *in vivo* advantage of *N. gonorrhoeae* fluoroquinolone-resistant *gyrA* mutants compared to those carrying wild-type GyrA 91S/95D in a mouse model (*38*) in the absence of fluoroquinolone administration. The marked difference in fitness between GyrA mutants is also consistent with *in vitro* estimates of fitness differences between resistant *gyrA* mutants in *E. coli*, as resistant *E. coli* GyrA 87G mutants were fitter than resistant 87Y mutants (*9*). These results underscore the importance of accounting for pathways to resistance when analyzing and modeling antimicrobial resistance dynamics.

Lineages carrying GyrA 91F/95G lost fitness after fluoroquinolones were no longer recommended, and persistence of sublineages point to *N. gonorrhoeae*’s strategies for responding to this fitness change. In lineage 20, one sublineage reverted to the susceptibility-conferring GyrA 91S/95D allele. Others changed the *mtr* locus, and one acquired azithromycin resistance through the C2611T 23S rRNA mutation (figure 2). At the same time, the sexual network in which the sublineages circulated appeared to change from men who have sex with men to heterosexuals (figure S2). While the number of isolates and sublineages limited quantification of these phenomena, they suggest responses to pressures from both antibiotics and host environments.

Similarly, for mosaic *penA* 34, we saw clear evidence of a large growth rate advantage compared to the baseline type that does not carry any of the resistance determinants studied when the cephalosporin cefixime was recommended. This advantage was rapidly lost after the switch to ceftriaxone 250mg, consistent with the observed decline of cefixime resistance (*39*).

For PBP1 421P, we estimated a consistent small fitness defect across the study period (1993-2019). PBP1 421P mainly provides penicillin resistance with only relatively modest increase in cephalosporin MICs (*28*), but the absence of a fitness benefit suggests this resistance phenotype was insufficient even in the context of cephalosporin use to confer an advantage. Moreover, the carriage of PBP1 421P in 12 of 29 lineages is consistent with at most a mild fitness defect. Consistent with this result, murine competition assays demonstrated that in isogenic strains, the strain carrying PBP1 421P was less fit than the one carrying PBP1 421L, regardless of which PBP1 allele was originally native to the strain background. We conclude that the PBP1 421P allele is likely a relic of the decades when penicillin was the backbone of *N. gonorrhoeae* infection treatment (figure S1).

We estimated a large growth rate benefit of mosaic *mtrR* promoters once azithromycin co-treatment was introduced in 2010. This is consistent with the findings of continued rapid expansion of lineages carrying mosaic *mtrR* promoters noted in Europe (*40*), but we note that the interaction with other *mtrCDE* mosaics makes the overall picture challenging to interpret. In the dataset we used, mosaic *mtrR* promoters always co-occur with at least one of mosaic *mtrC* or *mtrD*, both of which have their own distinct fitness impacts (table S2, figure 6). The trajectory of the largest lineage carrying mosaic *mtrR* promoter, lineage 29, plateaus around 2015 (figure S42). A similar behavior can be seen in the overall prevalence of azithromycin resistance (figure S1), whereby the growth rate seems to decline post 2015. Possible explanations include a drop in azithromycin use (*41*) and sexual network-dependent fitness of the *mtrCDE* mosaics, as suggested by the over-representation of *mtrC* loss-of-function alleles in cervical specimens (*42*).

Investigating the fitness contributions at the level of individual lineages (figure 7, figures S14–S41) allowed us to interrogate how much of each lineage’s growth trajectory can be explained by the combination of AMR and changes in treatment policy. This revealed an example of hitchhiking, where in Lineage 22 (figure 7) the fitness cost incurred by one resistance determinant, GyrA 91F/95G, conferring resistance to fluoroquinolones, was outweighed by the fitness benefit from another resistance determinant, mosaic *penA* 34, conferring resistance to cephalosporins. Several lineages did not carry any of the determinants and displayed a consistent trend in their residual terms. In the case of Lineage 2 (figure S15), we identified the presence of *tetM*, which confers resistance to tetracycline class antibiotics. This suggests that for some lineages, there may be a substantial impact of bystander exposure on their fitness trajectories (*41*, *43*). A similar phenomenon may explain the dynamics of Lineages 3 and 4. Incorporating population-wide antimicrobial use may enable quantification of the impact of bystander exposure. Lineages that displayed large residual effects despite carrying resistance determinants included in our model may reflect the impact of lineage background, environment pressures, or bystander exposure, suggesting avenues for further investigation.

Our approach has limitations. First, we were only able to uncover sufficient signal in the data to quantify the impact of determinants that have a large effect or that appear on multiple lineage backgrounds. Even as we captured the dominant effects of resistance determinants, there was too much uncertainty to define the impact of many resistance determinants on fitness landscape of *N. gonorrhoeae*. Reducing the uncertainty requires either a larger number of sequences, more representative sampling, or both. This may also enable use of birth-death-sampling processes (*44*), especially variations of the multi-type birth-death process (*45*). Larger sample sizes and higher data quality would enable more robust estimates under more complex models that could, for example, accommodate time-varying relationships between prescribing data and the growth rate effect of resistance determinants. Second, in our study, deviations from the model are captured by the residual terms. To include these phenomena, non-parametric methods such as splines or Gaussian processes could be used to model the relationship between treatment composition, time, and growth rate effect of determinants. Third, the need to explicitly define fixed lineages *a priori* is an approximation and may result in fragmentation of otherwise linked lineages. Fourth, the approach presented is only applicable for determinants that give rise to lineage-like dynamics. This effectively means that the estimates for the determinants are valid for sufficiently compatible genetic backgrounds where any putative fitness cost is not too large. If the fitness cost was large, the observed dynamics would likely resemble mutation-selection balance in the case of strong mutation and strong negative selection (*8*). Fifth, our approach can only estimate the association between the presence of resistance determinants if carried by sufficiently successful or ‘major’ lineages. In effect, this conditions on determinants being present on compatible genetic backgrounds, as it is unlikely that a clone would give rise to a major lineage in the absence of compatibility between the genetic background and the resistance determinant. Sixth, we have ignored any spatial heterogeneity in transmission and treatment. As the data collection is spatially very sparse, heterogeneity within the US in transmission and treatment is unlikely to impact the results. Lastly, importations from outside of the US may distort the results. Due to the focus on only major lineages, and the size of the *N. gonorrhoeae* epidemic in the US, we do not expect this to play a major role, and any remaining effects of importation should be compensated by the residual over-dispersion terms in the statistical model.

Our results demonstrate how the expanding collection of microbial genomic data together with antibiotic prescribing data and phylodynamic modeling can be used to explain microbial ecological dynamics and quantify the fitness contributions of genetic elements in their changing *in vivo* environments. The power of this approach will be augmented with continued surveillance, sequencing, and systematic data collection, and the growing datasets will enable model refinement and development. While we focused on *N. gonorrhoeae* and AMR, these methods could be more broadly applied to other microbes and pressures, aiding in efforts to understand how combinations of genetic elements inform strain fitness across antimicrobial exposure, host niche, and other environmental pressures.

## Acknowledgments

We thank Daniel H. F. Rubin for the pDR53 plasmid.

## Funding

This work was supported by NIH R01 AI132606 to Y.H.G. and R01 AI153521 to Y.H.G., A.E.J., and R.A.N.

## Author contributions

DH and YHG conceptualized the study. DH and TDM designed and performed the computational analysis. AM, GG, and ALV performed the competition assays. DH and YHG wrote the original draft. YHG acquired funding. DH, YHG, TDM, AM, SGP, RAN, and AEJ discussed the results and contributed to writing, reviewing, and editing of the manuscript.

## Competing interests

There are no competing interests to declare.

## Data and materials availability

The code and data necessary to reproduce the statistical analysis, along with the metadata and accession numbers for the isolates analyzed, are available at: https://github.com/gradlab/GC_AMR_Lineages.

## Supplementary materials

Materials and Methods

Figs. S1 to S44

Tables S1 to S9

References (*46–85*)

## Supplementary Materials

### Materials and Methods

#### Genomic Analysis

We collected publicly available genomic data, minimum inhibitory concentrations, and demographic data from GISP isolates (n = 5367) sequenced between years 2000 and 2019 (*3*, *14–17*). *De novo* assembly was performed using SPAdes v 3.12.0 (*46*) with the –careful flag, and reference-based mapping to NCCP11945 (NC_011035.1) was done using BWA-MEM v 0.7.17 (*47*). We used Pilon v 1.23 to call variants (minimum mapping quality: 20, minimum coverage: 10X) (*48*) after marking duplicate reads with Picard v 2.20.1 (https://broadinstitute.github.io/picard/) and sorting reads with samtools v 1.17 (*49*). We generated pseudogenomes by incorporating variants supported by at least 90 % of reads and sites with ambiguous alleles into the reference genome sequence. We mapped reads to a single copy of the locus encoding the 23S rRNA and called variants using the same procedures (*50*). We identified resistance-associated alleles from *de novo* assemblies and pseudogenomes. Likewise, we identified the presence of single nucleotide variants (e.g., mutations in *gyrA*, *parC*, *ponA*, and *penA*) and the copy number of resistance-associated variants in 23S rRNA from variant calls. To determine the presence or absence of genes, mosaic alleles, promoter variants, and small insertions or deletions we used the results of blastn v 2.9.0 (*51*) searches of assemblies for resistance-associated genes. We typed mosaic *penA* alleles according to the nomenclature in the NG-STAR database (*52*). We defined mosaic *mtr* alleles as those with <95% identity to the *mtr* operon encoded by FA1090 (NC_002946.2). Alleles were defined as loss-of-function (LOF) if frameshifts or nonsense mutations led to the translated peptide being less than 80% of the length of the translated reference allele.

Prior to phylogenetic reconstruction, we filtered assembled genomes based on the following criteria: 1; The total assembly length was longer than 1900000 bp and less than 2300000 bp. 2; Reference coverage was more than 30%. 3; Percentage of reads mapped to reference was at least 70%. 4; Less than 12% of positions were missing in pseudogenomes. This resulted in (n = 4573) retained samples.

#### Lineage Assignment & Phylogenetic Reconstruction

We used GUBBINS (*53*) to estimate recombining regions and IQTREE (*54*) for phylogenetic reconstruction. The molecular clock model was *GTR+G+ASC* as selected by MODELFINDER (*55*).

We estimated the dates of ancestral nodes using the resulting tree with BACTDATING (*56*) under the additive relaxed clock model (*57*). We estimated ancestral states for all determinants under study apart from the 23S rRNA as the joint maximum likelihood estimate under the F81 model (*58*) using PASTML (*59*). In the case of *penA*, the ancestral state reconstruction was performed using allele types and the resulting reconstruction was then mapped to *penA* determinants for subsequent analysis and lineage calling. For the 23S rRNA, we used maximum-parsimony ancestral reconstruction based on the DELTRAN algorithm (*60*) as implemented in PASTML (*59*). We chose this approach because the C2611T substitution is usually present in 4 copies, making reverse mutation unlikely, and DELTRAN prioritizes parallel mutation (*60*). Furthermore, the 23S rRNA C2611T variant does not display a clonal pattern of inheritance apart from one exception (figure S43).

We excluded samples with missing values in any of the determinants from the analysis prior to ancestral state reconstruction, leaving (n = 5215) samples. We defined a subset of tips as a lineage if it was the maximal subset such that there was no change of ancestral state in any of the loci across the unique path from each tip to the most recent common ancestor of the subset and the timing of the most recent common ancestor was estimated to no earlier than 1980 with at least 99% posterior probability. We then defined included lineages in the analysis if they contained at least 30 tips.

#### Lineage-Based Hierarchical Phylodynamic Model – Overview and Rationale

Our aim was to study how interactions among six resistance-associated genes and operons– *gyrA*, *parC*, *ponA*, *penA*, the *mtr* operon, and the 23S rRNA–and the major antimicrobial classes used as primary treatment of gonorrhea between 1993-2019 (table S1; (*28*, *61*)) affected the success and failure of resistant *N. gonorrhoeae* lineages in the US. For *gyrA*, we considered alleles given by codons 91 and 95. For *ponA*, we considered the L and P variants encoded at codon 421. For *parC*, we considered alleles given by combinations of codons at positions 86, 87, and 91. For each of the loci that make up the *mtr* operon (*mtrC*, *mtrD*, *mtrR*, and the *mtr* promoter), we considered whether the locus was non-mosaic, mosaic, affected by a loss-of-function mutation, and whether there was an A-deletion in the *mtr* promoter. For *penA*, we considered variants at each site listed by NG-STAR *penA* allele types (*52*), along with whether the *penA* allele was mosaic. Within the lineages derived from our dataset, we only observed variation at codon sites 501 and 542 and the presence of the mosaic *penA* 34 allele. The mosaic *penA* 34 allele carries variants at other sites; however, since these variants do not occur on other backgrounds in the dataset, we could not estimate their contributions. The determinants at *parC* act as mutations modulating the impact of *gyrA* resistance mutations (*30*). Consequently, we used partial pooling to estimate the effects of *gyrA* determinants across different *parC* contexts. For 23S rRNA we considered the presence of at least one copy carrying the C2611T substitution. While the 23S substitution A2059G is associated with a more dramatic increase in azithromycin resistance, we did not include it in any analysis as it appeared in fewer than 20 isolates. We did not include determinants at the *porB* locus despite this locus contributing toresistance (*28*), as these determinants are rapidly lost and re-acquired with a limited amount of vertical inheritance. This was evident from previous phylogenetic analysis of *N. gonorrhoeae* resistance (*17*), and our own analysis <u>5.44</u>.

We used phylodynamic modeling (*62*) to mitigate the impact of inconsistent sampling on reconstructing lineage ecology. Because coalescent phylodynamic modeling can be less sensitive than traditional incidence-based modeling to violations of sampling assumptions (*63*), it can accommodate the overrepresentation of antibiotic resistant specimens in collections of sequenced isolates (*3*, *14*, *15*).

The data used in the statistical model consisted of 1; *L* genealogies *G* = {**g***_i_*}_1≤*i*≤*L*_, each corresponding to a particular AMR-linked lineage (figure 1); 2; resistance determinant presence by lineage (table 1); and 3; treatment data from GISP clinics (figure S1). As our aim was to quantify the impact of individual AMR determinants on lineage success and failure, we estimated the growth rate of the lineage-specific effective population size *r*(*t*) = *Ne*(*t*)/*Ne*(*t*) (*22*, *64*). We extended prior work (*22*, *64*) to a multiple lineage, multiple treatment, multiple AMR determinant scenario by constructing a hierarchical Bayesian regression model that accounted for intrinsic variation among lineages. We formulated the growth rate of the effective population size as a hierarchical linear model to estimate how much of lineage growth and decline could be explained as a function of the interaction between AMR determinants and the pattern of antimicrobial use. Disentangling the contributions of individual AMR determinants from external factors required accounting for the overall epidemic dynamics, for which we included a global trend term shared by all lineages; the effect of lineage background on baseline fitness, for which we included lineage-specific terms; and the over-dispersion in the growth and decline of individual lineages that occurs due to factors unaccounted for.

The growth rate of the effective population size serves as a proxy for lineage success and can be used to solve for the effective population size (equation 5.1). While the effective population size is not necessarily directly proportional to incidence (it is a non-linear function of incidence and prevalence (*63*)), if fitness benefits are small in comparison to the per capita transmission rate *β*(*t*) or if *β*(*t*) is approximately constant, then the growth rate of the effective population size will approximately match the growth of the epidemic (*64*). The effective population size can then be linked to individual genealogies via the coalescent likelihood (equation 5.2). The key quantity of interest was the marginal impact of individual determinants on lineage growth rates. This is formulated in (equation 5.4) and (equation 5.5).

Coalescent based phylodynamic approaches condition on sampling and are therefore more robust to sampling bias across individual lineages (*63*). The sampling bias present in the data from (*3*, *14*, *15*) is a form of ascertainment bias for macrolide and cephalosporin resistant phenotypes. The lineages are defined based on resistant determinants;, and thus the resistance phenotype is approximately identical within a lineages. This means that the ascertainment bias translates to different sampling intensity between lineages, but not within a lineage. We reasoned that the majority of the impact of the ascertainment bias should be mitigated as the coalescent likelihood of a given lineage (see Materials and Methods: *Lineage-Based Hierarchical Phylodynamic Model – Construction of the Likelihood*) is conditional on sampling. We therefore worked within the coalescent framework as opposed to a birth-death-sampling process framework to mitigate the impact of sampling being artificially enriched for particular resistant phenotypes. While the effective population size relates to census size only through a complex and potentially non-linear relationship, in the case of infectious disease this relationship can be made explicit under mild assumptions (*63*, *65*, *66*). In the context of AMR resistance, phylodynamic approaches have been used in e.g. (*19*), (*20*), or (*22*). These applications been limited to scenarios that either considered an interaction between a single population and covariate – in the AMR case the covariate being treatment usage data as in (*19*), or to a simplistic scenario with multiple populations and only a single covariate as in (*22*). With the exception of (*21*, *23*), all approaches have been restricted to comparisons between resistant and susceptible types. In this work we studied multiple sites as in (*21*, *23*), but in contrast we introduced several covariates in the form of antimicrobial use data in order to include the impact of a changing environment.

#### Lineage-Based Hierarchical Phylodynamic Model – Construction of the Likelihood

Our aim was to study to what extent the growth and decline of individual lineages can be explained through the interaction of the motifs a given lineage carries at each of the loci in *S* and the changes in direct antimicrobial usage over time. Since we could not observe the population size directly, we instead modeled the effective population sizes from the genealogies in *G*, and used these estimates to inform past population dynamics using what is frequently referred to as phylodynamic modeling (*62*, *66*, *67*). In particular, for a given lineage we aimed to estimate the growth rates of the corresponding effective population size 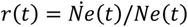 as has been done before in (*22*, *64*). From *r*(*t*), *Ne*(*t*) can be recovered as

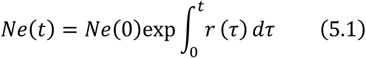

and related to the respective genealogy **g** using the coalescent likelihood (*68*).

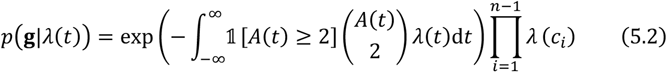

where *λ*(*t*) = 1/*Ne*(*t*), *A*(*t*) is the number of active lineages present in the genealogy at time *t*, also referred to as the block counting process (*69*), and *c_i_* are the times of internal nodes corresponding to coalescent events.

Since neither the integral in 5.1 nor the one in 5.2 are tractable, we discretized time as a regular grid of size *N*, denoting the end points of this discretization as *t*_1_ < … < *t_i_* < … < *t*_0_. We approximated *r*(*t*) as a piecewise constant function on this regular grid, and denoted this approximate quantity as 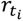.

We defined *Δ_t_* = *t_i_* – *t_i_*_-1_. We then approximated *r*(*t*), as well as *Ne*(*t*) as a piecewise constant function. Note that we measured time in reverse with *t*_1_ corresponding to present and *t*_0_ to the beginning of the first year for which treatment data was available. This approximation was originally introduced as a part of the skygrid approach (*70*).

For a lineage *l*, The discretized version of *Ne*^(*l*)^(*t*), denoted 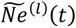 is then

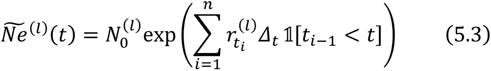

As 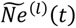 is a simple function, the integral in the coalescent likelihood was then trivial to evaluate.

#### Lineage-Based Hierarchical Phylodynamic Model – Detailed Description of the Statistical Model

In this section we will describe the statistical model that characterizes the lineage specific growth rates 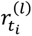 as function of what determinants a particular lineage carries and the the past treatment policies. As in SKYGROWTH (*19*), our aim was to model the growth and decline in the effective population size 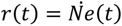 as a function of covariates, while accounting for over-dispersion. Unlike in (*19*) and akin to (*22*), we assumed multiple lineages. Expanding on both approaches, we modeled multiple covariates interacting with lineages through multiple resistance-linked loci, along with random effects to account for over-dispersion, and the impact of lineage backgrounds. First, some notation must be introduced.

First, we review the notation used for the data. By *G* = {**g***_i_*}_1≤*i*≤*L*_, we denote each of the *L* genealogies corresponding to the *L* lineages under study. We further consider 4 genomic regions associated with resistance, and denoted these by *S*. *S* consisted of an indicator for the *gyrA* locus, the *penA* locus, the *ponA* locus, 23S rRNA, as well as an indicator for the *mtrCDE* operon. For each lineage we kept track of AMR determinants present in each of the regions in *S*. Each region in *S* then interacts with the proportion at which a subset of the 4 treatment classes was used. We denoted the set of these treatment classes as *D*. *D* consisted of indicators for fluoroquinolones, cephalosporins other than ceftriaxone 250mg, penicillin, and azithromycin co-treatment. For each treatment class the data consisted of the proportion of cases treated with that class as primary treatment.

Let *m*(*s*): *s* ∈ *S* denote the set of determinant indices associated with the region *s* that can be found in that dataset we studied. For example for the *gyrA* locus this consisted of indices for 91F/95G, 91F/95A, and 91F/95N.

Let *I*(*s*) ⊆ *D*: *s* ∈ *S* denote the subset of treatment class indices that the region *s* interacts with (Table S). For example for *s* corresponding to *penA* locus, *I*(*s*) consists of the indices for cephalosporins other than ceftriaxone 250mg, and penicillin, respectively.

Let *u_t_*(*d*) ∈ [0,1]: *d* ∈ *D* denote the proportion of cases treated treatment *d* at time *t*.

Let *X_S_*(*j*, *l*) ∈ {0,1} be an indicator function equal to 1 if the determinant *j* at region *s* is present in lineage *l*, and 0 otherwise.

Finally denote the set of regions that depend on (known) compensatory mutations as *S^C^* ⊆ *S*. For each *s* ∈ *S^C^*, define the *G*(*s*) to be set of indices corresponding to the types of compensatory mutations. Let *g_S_*(*l*) ∈ *G*(*s*) be the type of compensatory mutation associated with region *s* that is present in lineage *l*. In our case *S^C^* consists of just one index for *gyrA*. For *gyrA*, *G*(*s*) consisted of indices corresponding to four different *parC* types: 86D/87S/91G, 86D/87R/91E, 86N/87S/91E, and 86D/87I/91E.

Using this notation we introduced functions that characterise the mean effect of AMR determinants on lineage *Ne*(*t*) growth rates. For determinant a *j* ∈ *m*(*s*) belonging to region *s* we denote the effect at time *t_i_* as 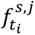:

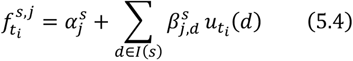

This can be understood as a linear model characterizing the impact of a given determinant on the *Ne*(*t*) growth rate of a lineage as a function of past treatment composition, compared to the impact of changes in treatment composition on the baseline. The coefficients 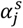 represent the change in intercept for determinant *j* in region *s* and the coefficients 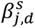 represent the change in the growth rate associated to the presence of determinant *j* in region *s* when the proportion of primary treatment corresponding to antimicrobial class *d* changes. Note that for regions that depend on compensatory mutations these coefficients represent the average coefficient over all compensatory mutation backgrounds.

For those determinants that depend on compensatory mutations, we further defined the effect of the compensatory mutation background as

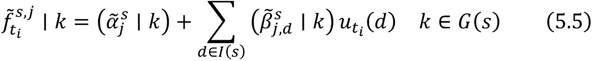

The terms 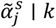 represent the change in the intercept term for determinant *j* on background indexed by *k* compared to the the determinant’s mean intercept 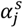. Analogously 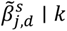 represent the impact of the compensatory background indexed by *k* on the antimicrobial-determinant interaction coefficients.

We modeled the growth rate of lineage *l* at time *t_i_*, denoted by 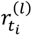 as:

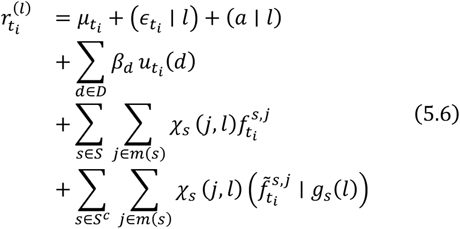

The first line corresponds to the global trend 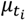, the lineage background intercept *a* | *l*, and the lineage residual term 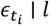 that accounts for over-dispersion in the lineage trajectory.

The second line corresponds to the impact of changing treatment policy on the growth rate of lineages with only the baseline motif. This impact is represented by coefficients *β_d_*.

The third line contains the mean effects of the determinants carried by a particular lineage on its *Ne*(*t*) growth rate.

The final line accounts for the impact of compensatory mutation background on the effect of determinants present in the corresponding regions.

The last step was to characterise the prior model that was used for the parameters included in 5.6. Note that the choice of priors required a choice of a time scale. We therefore set the unit of time to one year.

We modeled the global trend 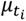 as a Gaussian random walk with independent, stationary increments, and an unknown marginal standard deviation *τ_μ_* itself equipped with a half-normal prior:

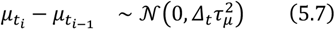

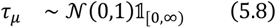

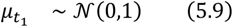

We modeled the lineage specific residual terms 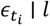 as:

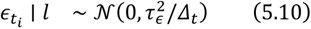

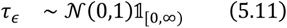

This can be understood as a multi-level model component accounting for temporal trends in lineage trajectories not explained by either the motifs that we focus on or the global trend.

The hierarchical prior on the lineage intercept terms was:

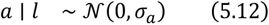

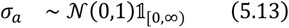

For the baseline coefficients treatment impact coefficients *β*_8_ we used standard normal priors, which corresponds to a weakly informative choice:

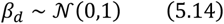

We also used a similar strategy for the site and motif specific interaction coefficients 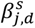. Motifs at loci associated with resistance to several classes of antimicrobials (Table S) will typically increase or decrease resistance to all of these classes at once. To include this, we modeled 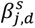 for a fixed *s*, *j* as correlated:

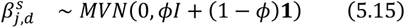

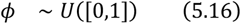

Where *MVN* stands for a multi-variate normal distribution, **1** represents the matrix with unit entries, *I* is the identity matrix and *U*([0,1]) is the uniform distribution on the unit interval. *ϕ* is a pooling factor that determines the level of correlation between the individual antimicrobial specific coefficients. This parametrization preserves a marginal variance of 1.

Finally, we modeled the effect of compensatory mutation background with the following prior

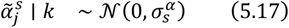

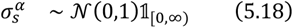

And

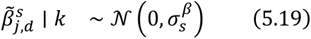

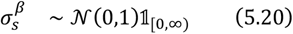

respectively. This hierarchical prior enables us to estimate the overall impact of compensatory mutations.

#### Lineage-Based Hierarchical Phylodynamic Model – Implementation

The model was implemented in the STAN probabilistic programming language (*71*) and R language version 4.4.0 (*72*). Sampling was performed using Hamiltonian Monte Carlo as implemented in STAN (*71*). Four chains were run in parallel for 1000 sampling iterations each. For all model parameters, the bulk effective sample size (bulk-ESS) was always at least 500, the R^ statistic always lower than 1.05 (*73*).

#### Comparison of the Impact of GyrA 91F/95G versus GyrA 91F/95A on ciprofloxacin MICs

To investigate possible differences between the levels of fluoroquinolone resistance we selected all isolates that 1; carried ParC 86D/87R/91E 2; carried GyrA 91F/95G or GyrA 91F/95A. For each of these isolates, we then log_2_-transformed the corresponding ciprofloxacin MIC, removing any isolates that were either missing a ciprofloxacin MIC value or had a reported MIC of 0. To assess whether the carriage of GyrA 91F/95G as opposed to GyrA 91F/95A on ParC 86D/87R/91E background, we then fitted a linear model using the lm function in R version 4.4.0 (*72*), while controlling for determinants at the *mtrCDE* operon. The model used was standard linear regression with categorical predictors as specified by the following regression formula:

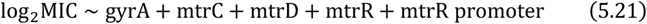

#### *gyrA* Mutants Competition Assay – *N. gonorrhoeae* culture conditions

*N. gonorrhoeae* was cultured on GCB agar (Difco) supplemented with Kellogg’s supplement (GCB-K) at 37^°^C with 5% CO_2_ (*74*). We performed pairwise competition experiments in liquid GCP medium containing 15g/L proteose peptone 3 (Thermo Fisher), 1g/L soluble starch, 1g/L KH_2_PO_4_, 4g/L K_2_HPO_4_, and 5g/L NaCl (Sigma-Aldrich) with Kellogg’s supplement (*75*).

#### *gyrA* Mutants Competition Assay – Generation of isogenic *N. gonorrhoeae* strains and antibiotic susceptibility testing

Antibiotic susceptibility testing for ciprofloxacin was performed on GCB-K agar via Etest (BioMerieux). All minimum inhibitory concentration (MIC) results represent the mean of three independent experiments. Strains, plasmids and primers used for *gyrA* mutant experiments are listed in (tables S3 – S6). All the isogenic *N. gonorrhoeae* strains were generated in a ciprofloxacin-resistant clinical isolate, GCGS0481, which carries GyrA 91F/95G and ParC 87R. To clone a GyrA 91S/95D fragment with a chloramphenicol resistant cassette (CMR), pAM_3 plasmid was constructed using Gibson assembly in a pUC19 (*76*) backbone. The GyrA 91S/95D fragment was amplified from pDRE77 (*77*) using the primer pair AM_7 and AM_8 and the chloramphenicol cassette from pKH37 (*78*) using the primer pair AM_9 and AM_10. Fragments were amplified using Phusion high-fidelity DNA polymerase (NEB), checked for appropriate size by gel electrophoresis, column purified (Qiagen PCR purification kit), assembled with Gibson Master Mix (NEB), and transformed into chemically competent DH5*α E. coli* (Invitrogen). Individual colonies were selected on LB agar supplemented with 20*μ*g/mL chloramphenicol and grown overnight at 37^°^C. Plasmids were isolated using Miniprep Kit (Qiagen) according to the manufacturer’s instructions and sequences were confirmed by Sanger sequencing. For the insertion of GyrA 91S/95D allele into *N. gonorrhoeae* GCGS0481, the isolate was grown overnight on a GCB-K plate at 37^°^C with 5% CO_2_. After 16-20 hours, the strain was scraped and suspended in 0.3M sucrose (Sigma-Aldrich), electroporated with 200 ng of pAM_3 plasmid, and rescued with GCP medium supplemented with Kellogg’s for 30 minutes. The transformants were then plated on non-selective GCB-K agar plates for 4-6 hours followed by selection on GCB-K plates supplemented with 4.5*μ*g/mL chloramphenicol. Finally, individual colonies were re-streaked on non-selective GCB-K agar plates and the *gyrA* allele checked by Sanger sequencing. For cloning of GyrA 91F/95G and GyrA 91F/95A, fragments of *gyrA* were amplified using primers AM_5 (F) and AM_6 (R) from the genomic DNA of clinical *N. gonorrhoeae* isolates GCGS0481 and NY0842 respectively. Electroporation was done as described above, and individual colonies were selected on GCB-K plates supplemented with 2*μ*g/mL ciprofloxacin. For all the transformations performed, transformations without DNA were used as negative controls.

#### *gyrA* Mutants Competition Assay – Competitive fitness measurement of GyrA variants

GCGS0481 GyrA 91S/95D, GyrA 91F/95G and GyrA 91F/95A containing the CMR cassette were transformed with pDR53, a kanamycin cassette (KanR) derivative of pDR1 (*79*) (constructed using the primer pair DR_395 and DR_396). The resulting transformants were selected on GCB-K agar supplemented with 70*μ*g/ml kanamycin. Colony PCR was performed to screen the kanamycin positive clones using the primer pair (DR_62 and DR_63) (table S6). During the pairwise competition experiments, the competitive paired strains from overnight cultured plates (one kanamycin-sensitive and one kanamycin-resistant strain) were mixed and co-cultured (at a ratio of 1:1 by optical density) in antibiotic-free GCP media with Kellogg’s supplement for 8 hours. At each timepoint, cultures were serially diluted, and same volume was plated on both GCB-K agar and GCB-K agar supplemented with 70*μ*g/ml kanamycin. Finally, dilutions on both plates were quantified and the competitive index (CI) was calculated at each timepoint by counting CFUs. The CI value at any timepoint was calculated as (*R_t_*/*S_t_*)/(*R*_0_/*S*_0_) where *R_t_* and *S_t_* are the proportions of kanamycin-resistant and kanamycin-sensitive strains, respectively at any time point and *R*_0_ and *S*_0_ are the proportions of kanamycin-resistant and kanamycin-sensitive strains at time 0. Statistical analysis of CI measurements was performed using an unpaired two-sided Student’s t-test.

#### *ponA* Mutants Competition Assay – Generation of Strains and Plasmids

Strains, plasmids and primers used for *ponA* mutant experiments are listed in (tables S7 – S9). Strains used in these *in vivo* and *in vitro* competitive fitness experiments possess were made streptomycin resistant to facilitate the mouse infection studies (*80*). To achieve this, the *rpsL1* allele from FA1090, which encodes a mutant form of ribosomal protein S12 that confers streptomycin resistance (*81*), was transformed into the penicillin-resistant clinical isolate FA6140 (*34*) and the antibiotic-susceptible laboratory strain FA19 (*82*) by allelic exchange (*83*). Clones that acquired *rpsL1* were selected on GCB agar plates containing 100 *μ*g/mL streptomycin and verified by Sanger sequencing. FA19 rpsL and FA6140 rpsL were transformed with a pUC plasmid containing a portion of the *ponA* gene starting at bp 831 and harboring either 421L or 421P, the *aad1* resistance cassette (*Ω*) conferring spectinomycin/streptomycin resistance, and 531 bp of sequence downstream of *ponA* to facilitate recombination. Transformants were selected on GCB agar plates containing 25 *μ*g/mL spectinomycin and verified by Sanger sequencing. All gonococcal strains were propagated on solid GCB agar containing Kellogg’s supplements I and II (*74*) for 18 to 20 h at 37^°^C in a 5% CO_2_-enriched atmosphere.

#### *ponA* Mutants Competition Assay – *in vitro* Growth Curve Experiments

Nonpiliated bacterial colonies were harvested and inoculated into GCB supplemented with Kellogg’s supplements I and II (*74*). Cultures were shaken at 180 rpm at 37^°^C in a 5% CO_2_-enriched atmosphere, and bacterial growth was assessed by measuring the OD_600_ (Optical Density) at hourly intervals for a total of 8 hours. Experiments were repeated in biological triplicate, and statistical significance between individual strains was determined by using a repeated-measures 2-way analysis of variance (ANOVA) with Tukey’s multiple comparisons.

#### *ponA* Mutants – Competitive Murine Infections Experiments

Animal experiments were conducted at the Uniformed Services University of the Health Sciences according to the guidelines of the Association for the Assessment and Accreditation of Laboratory Animal Care under a protocol that was approved by the University’s Institutional Animal Care and Use Committee. Female BALB/c mice (6 to 8 weeks old; National Cancer Institute) were treated with water-soluble 17*β*-estradiol and antibiotics to increase susceptibility to N. gonorrhoeae (*84*). Groups of Female BALB/c mice were inoculated vaginally with 20 *μ*L of a PBS suspension containing similar numbers wild-type FA19 *rpsL* PBP1^421L^ and isogenic FA19 *rpsL* PBP1^421P^; or wild-type FA6140 *rpsL* PBP1^421P^ and isogenic FA6140 *rpsL* PBP1^421L^ CFU (total dose, 10^6^ CFU; 7 mice/group). Vaginal swabs were collected on days 1, 3, and 5 post-inoculation and suspended in 1.0 mL GCB. Vaginal swab suspensions and inoculum were cultured quantitatively on GCB agar with streptomycin (100 *μ*g/mL) for total CFUs and GCB agar with streptomycin (100 *μ*g/mL) and spectinomycin (25 *μ*g/mL) for FA19 *rpsL* PBP1^421P^ or FA6140 *rpsL* PBP1^421L^ CFUs. Results were expressed as the competitive index (CI) by counting CFUs. The CI value at any timepoint was calculated as (*R_t_*/*S_t_*)/(*R*_0_/*S*_0_) where *R_t_* and *S_t_* are the proportions of spectinomycin-resistant and spectinomycin-sensitive strains, respectively at any time point and *R*_0_ and *S*_0_ are the proportions of spectinomycin-resistant and spectinomycin-sensitive strains at time 0.

If only one strain was recovered from an infected mouse in a swab, the limit of detection of 1 CFU was assigned to the strain that was not recovered, and the CI was calculated and plotted at this limit of detection. When swabs yielded no culturable *N. gonorrhoeae*, the corresponding data point is omitted from the CI plot. FA19 competition experiments were performed in biological duplicate (total 7 mice/group). FA6140 competition experiments were performed once (total 7 mice/group). Statistical analysis of CI values was performed using Mann-Whitney test.

**Figure S1:**
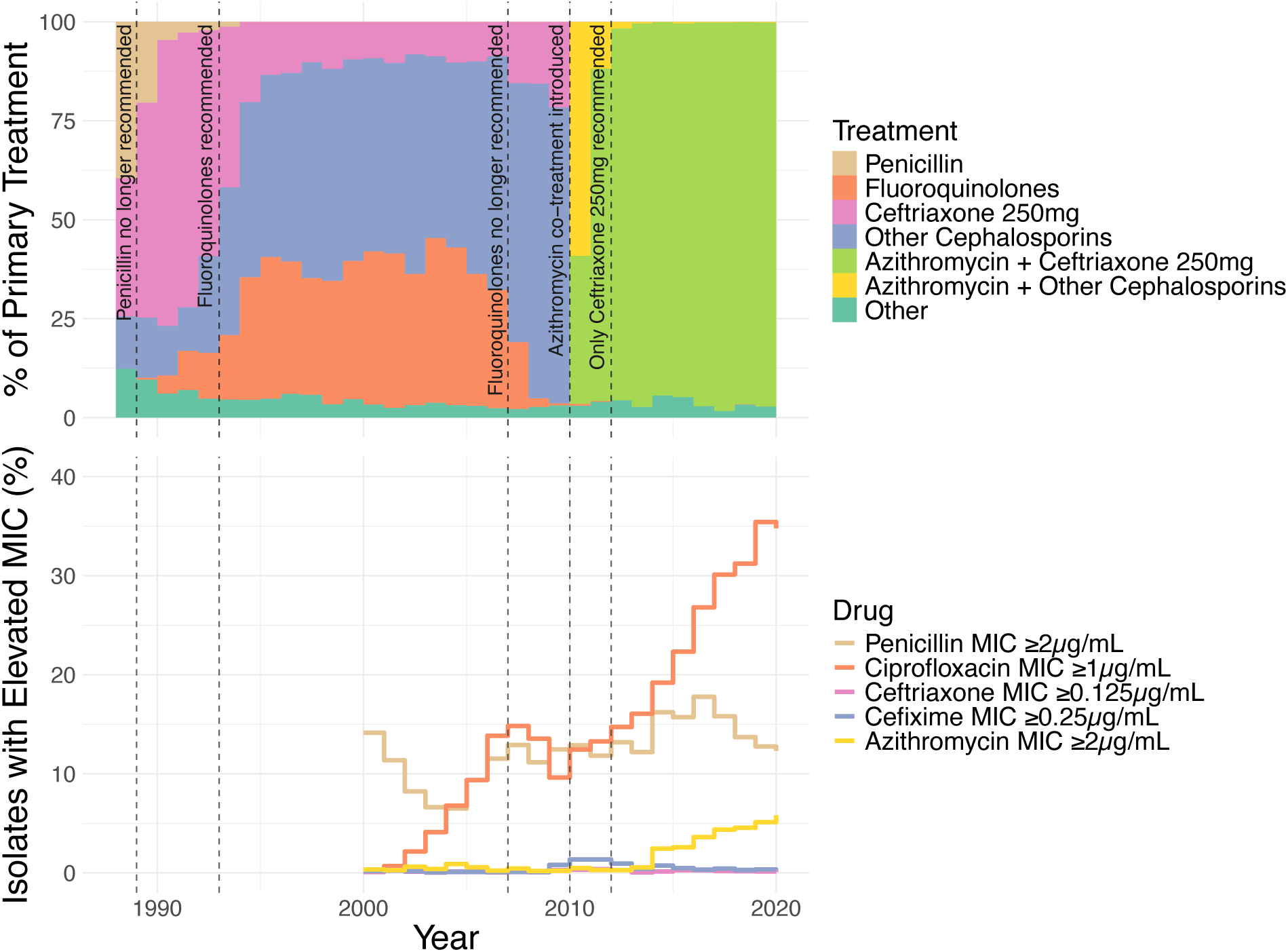
Gonococcal Isolate Surveillance Project treatment and resistance trends. Data from *CDC’s 2022 STI Surveillance Report*(*85*). Top panel: Past treatment composition. The fluoroquinolone category consists of ciprofloxacin and ofloxacin. Other cephalosporins consists of cefixime, ceftriaxone 125mg, and otherwise unspecified cephalosporins. Bottom panel: Prevalence of resistance over time. Note that resistance data is only available up from year 2000 onwards, and treatment data from 1988 onwards. CDC data does not report usage rates for azithromycin, however azithromycin co-treatment has been recommended with all cephalosporins since 2010 until 2020 (*85*). We therefore impute it to be equal to the total proportion for all cephalosporins past 2010.

**Figure S2:**
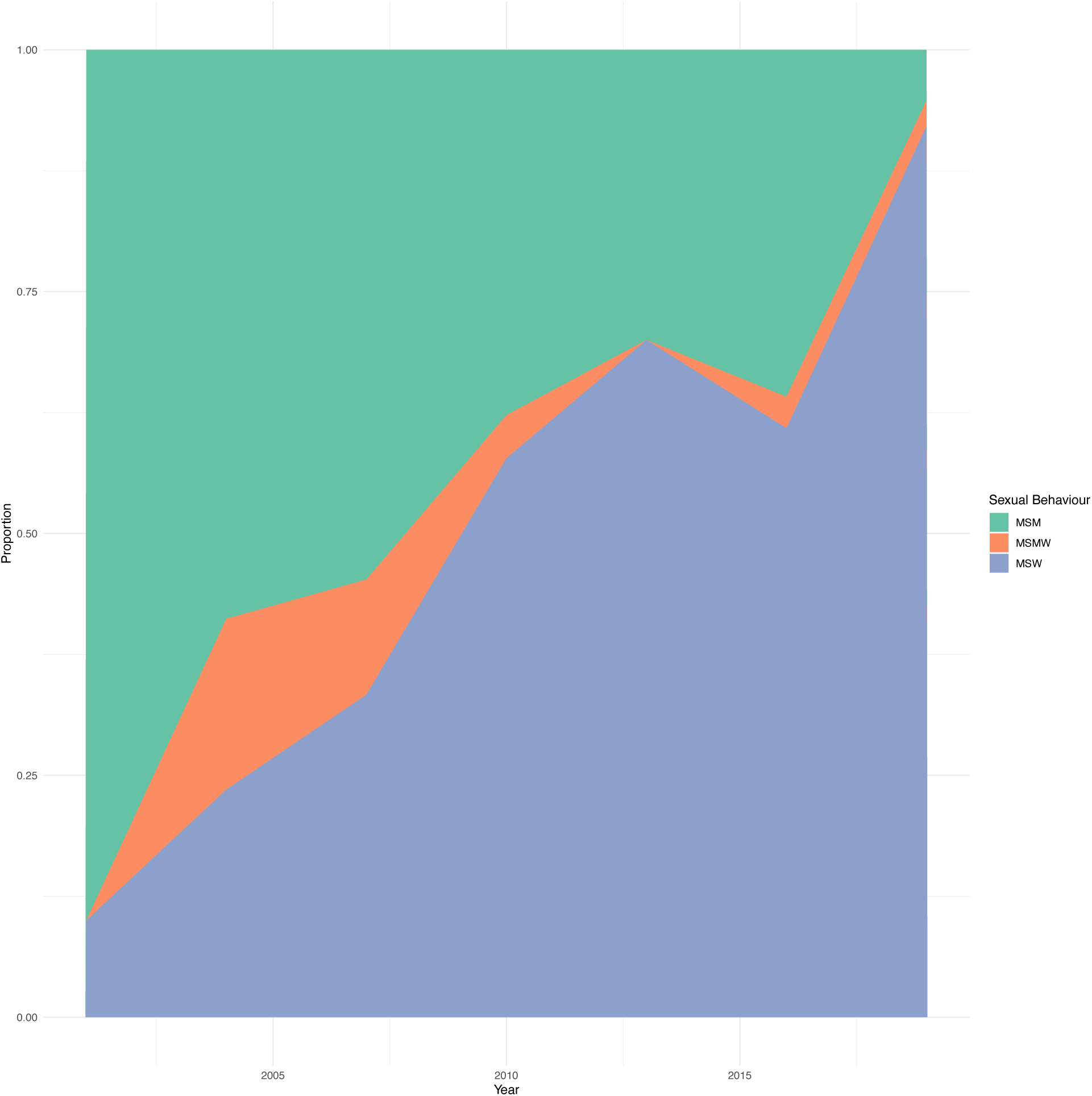
Distribution of reported sexual behaviour corresponding to all descendants of lineage 20. Proportions are calculated over 3 year intervals.

**Figure S3:**
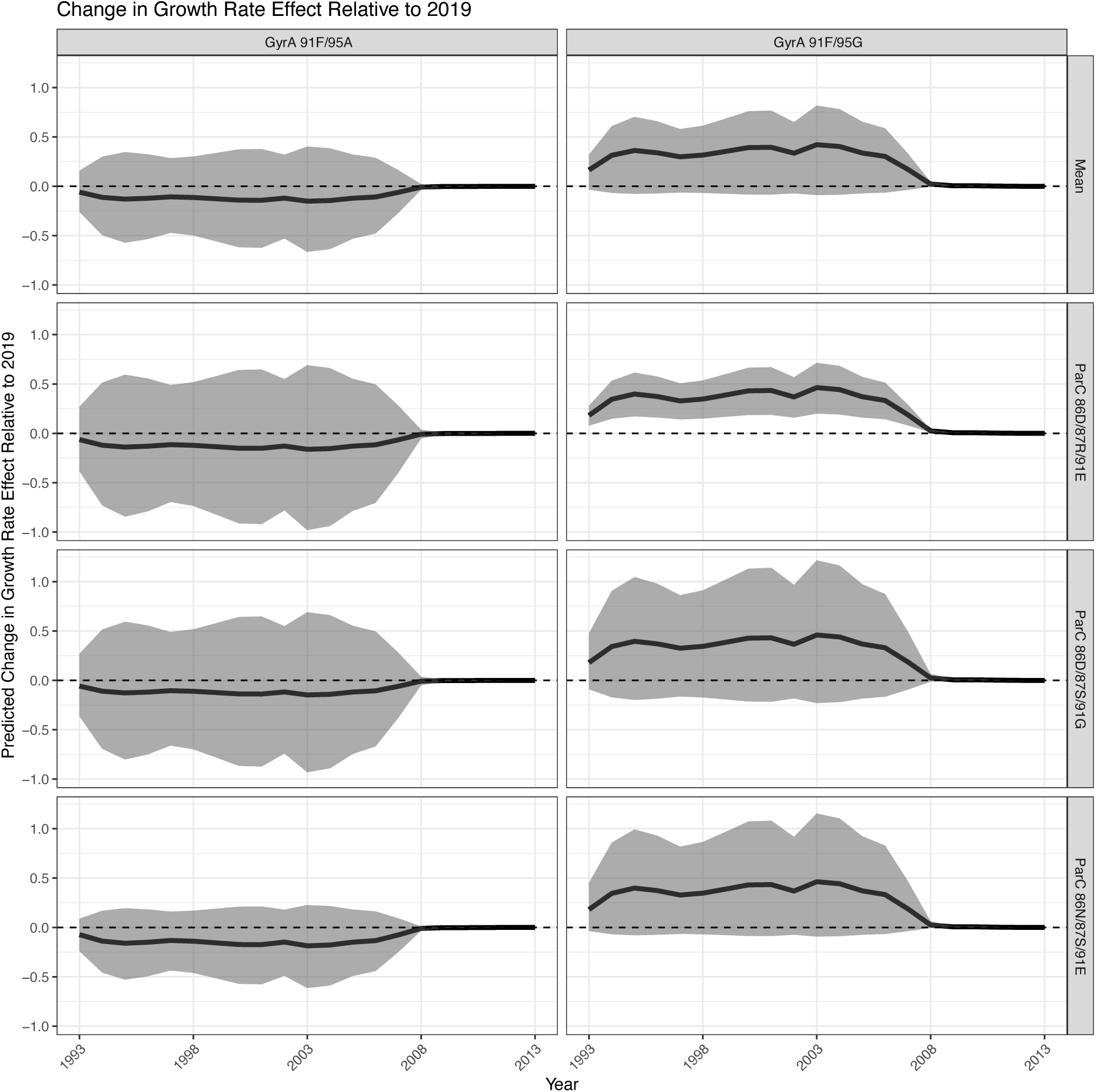
The predicted relative effect on growth rate for all *gyrA* determinants occurring in the dataset across all possible *parC* allele contexts. The predicted effect shows how the growth rate effect of a given determinant has changed based on the reported treatments compared to that determinant’s estimated growth rate effect in 2019. The average effect across all *parC* contexts for each of the *gyrA* alleles is denoted by Mean. The shaded region denotes the 95% posterior credible interval around the posterior median, depicted by the bold black line. Dashed line denotes no change in predicted effect of a given determinant compared to its’ predicted effect in 2019.

**Figure S4:**
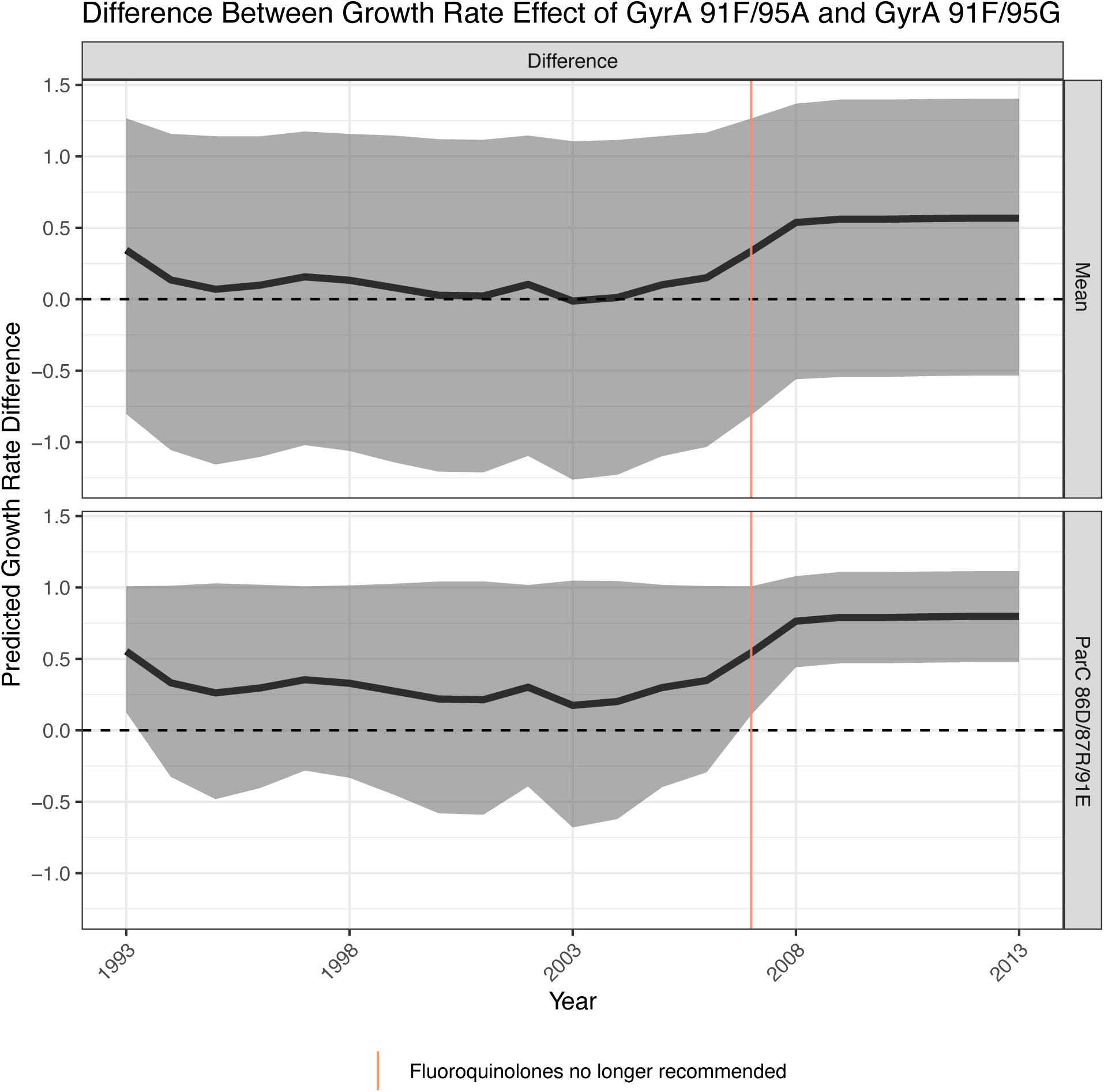
The difference between the predicted absolute growth rate effect as a function of fluoroquinolone usage of GyrA 91F/95G and GyrA 91F/95A averaged across all *parC* backgrounds and on ParC 86D/87R/91E background. The shaded region denotes the 95% posterior credible interval around the posterior median, depicted by the bold black line. Dashed line denotes 0.

**Figure S5:**
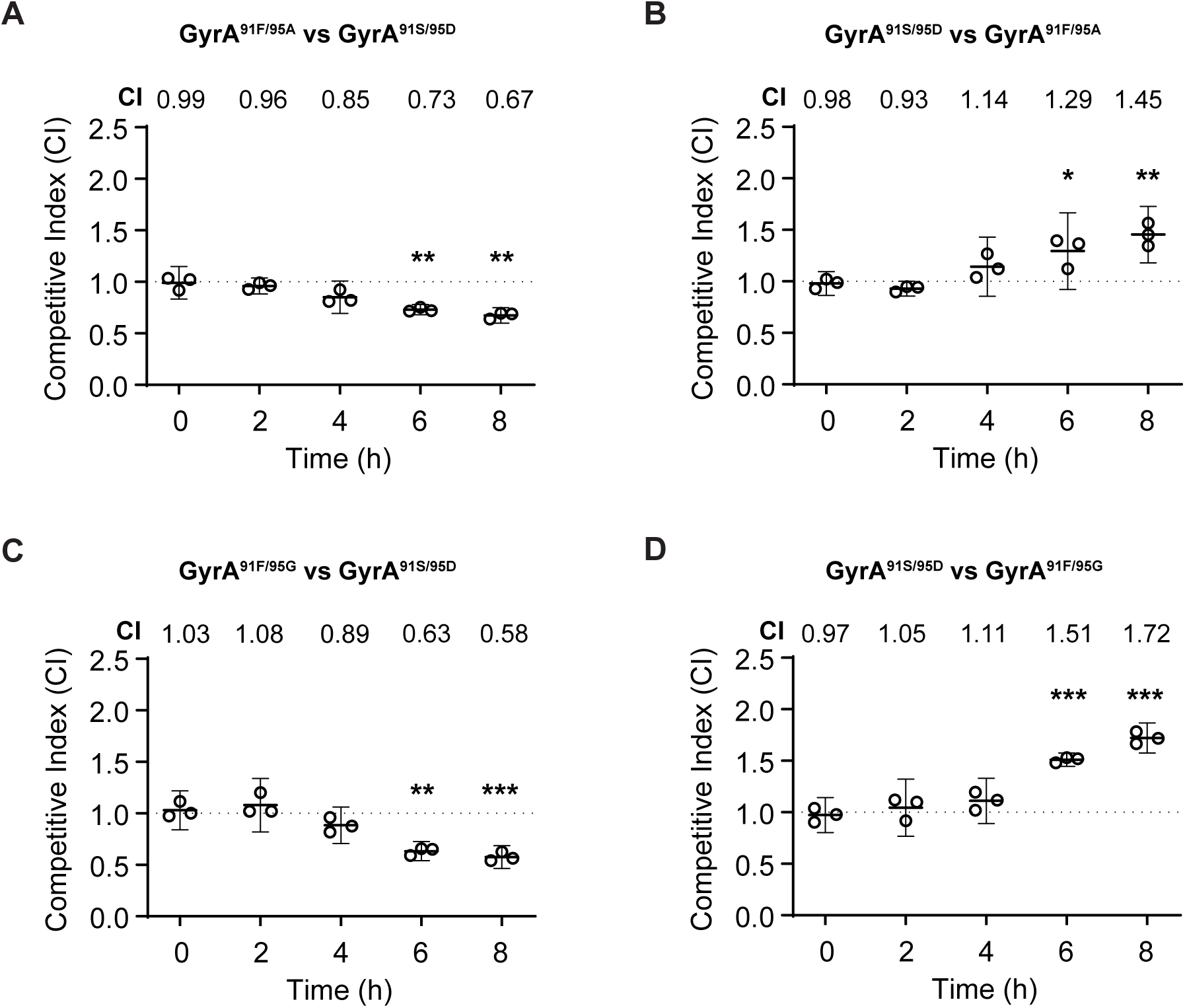
*In vitro* competition assays between GCGS0481 GyrA 91S/95D, GyrA 91F/95A and GyrA 91F/95G in the ParC 86D/87R/91E context. Panel A: Competition between unlabeled GCGS0481 GyrA 91S/95D and kanamycin-labeled GyrA 91F/95A. Statistical significance (for 2, 4, 6 and 8 hours, p = 0.49, 0.05, 0.0025, 0.0015 respectively). Panel B: Competition between unlabeled GCGS0481 GyrA 91F/95A and kanamycin-labeled GyrA 91S/95D. Statistical significance (for 2, 4, 6 and 8 hours, p = 0.19, 0.09, 0.03, 0.0023 respectively). Panel C: Competition between unlabeled GCGS0481 GyrA 91S/95D and kanamycin-labeled GyrA 91F/95G. Statistical significance (for 2, 4, 6 and 8 hours, p = 0.54, 0.07, 0.0013, 0.0009 respectively). Panel D: Competition between unlabeled GCGS0481 GyrA 91F/95G and kanamycin-labeled GyrA 91S/95D. Statistical significance (for 2, 4, 6 and 8 hours, p = 0.4, 0.1, 0.0002, 0.0001 respectively). N = 3/timepoint, representative of three independent experiments performed in absence of any antibiotic pressure. Error bars represent mean with 95% CI. Statistically significant differences in CI values were analyzed using an unpaired two-sided Student’s *t-test* and are indicated (\**p* < 0.05, \*\**p* < 0.005 and \*\*\**p* < 0.0005).

**Figure S6:**
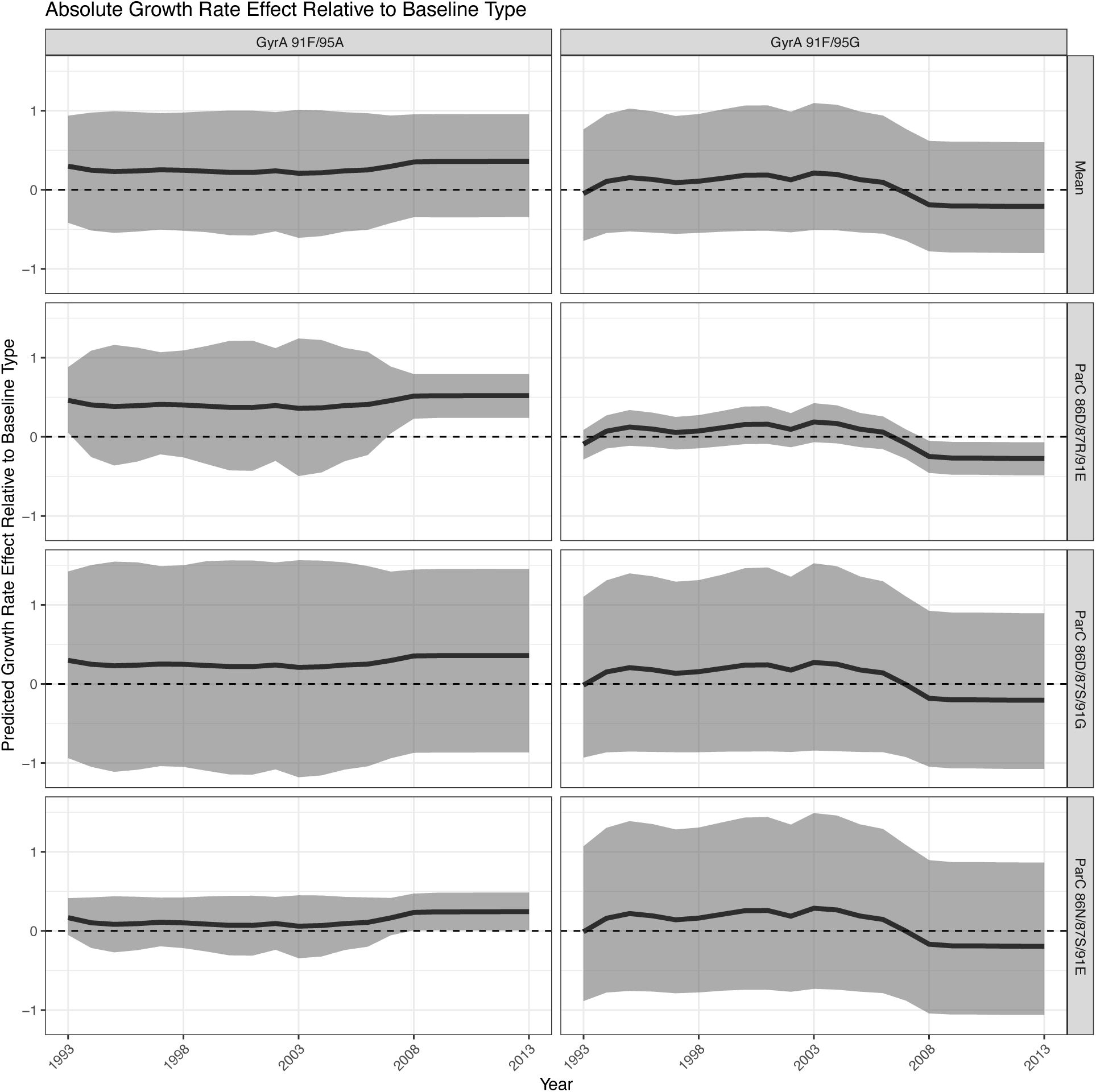
The predicted the absolute growth rate effect for all *gyrA* determinants occurring in the dataset across all possible *parC* allele contexts. The predicted effect was computed based on reported treatments. The average effect across all *parC* contexts for each of the *gyrA* alleles is denoted by Mean. The shaded region denotes the 95% posterior credible interval around the posterior median, depicted by the bold black line. Dashed line denotes no predicted growth rate effect relative to the baseline type that does not carry any of the resistance determinants studied.

**Figure S7:**
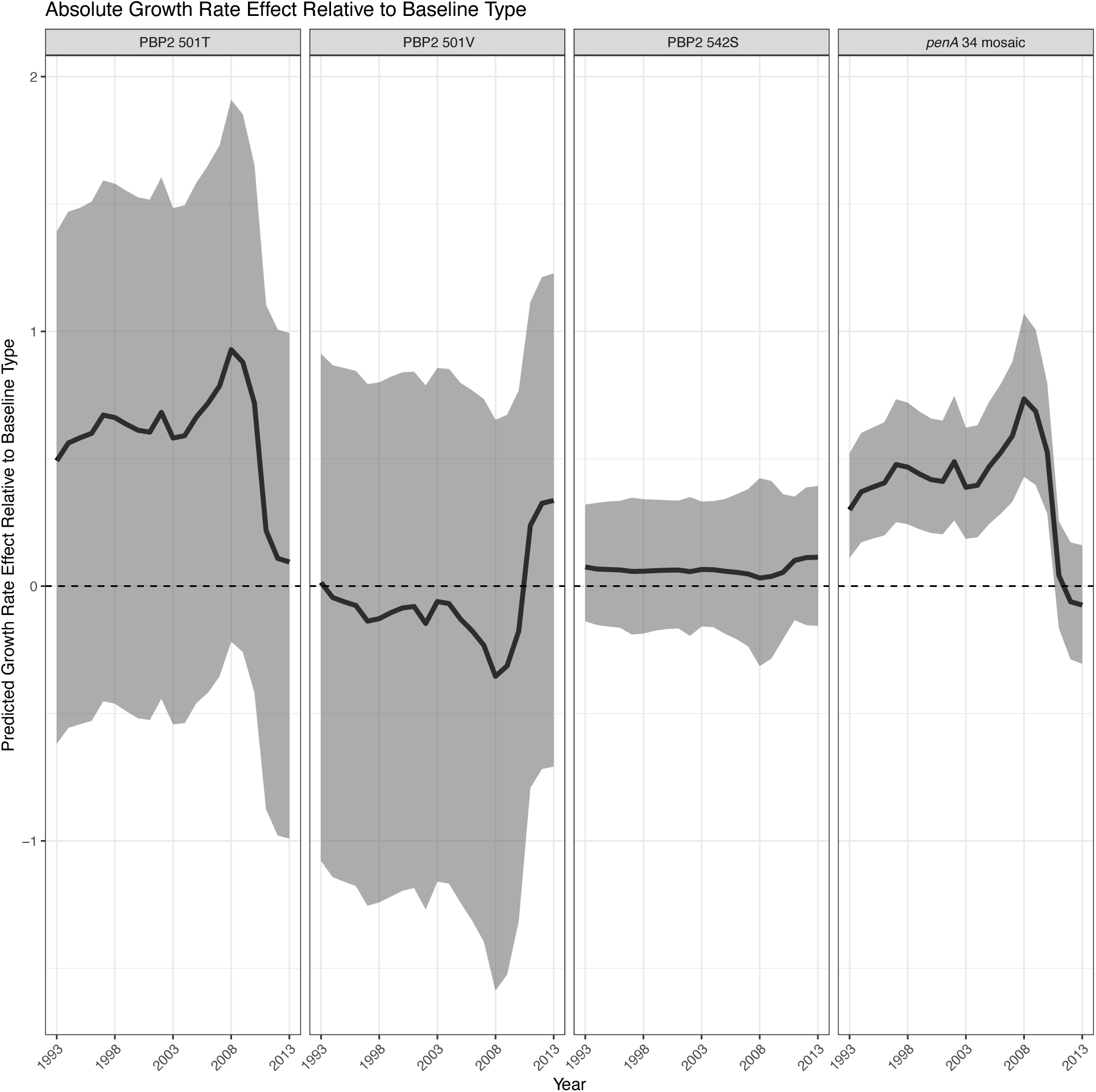
The predicted the absolute growth rate effect for all determinants at the *penA* locus occurring in the dataset on lineage growth rate. The predicted effect was computed based on reported treatments. Shaded region denotes the 95% posterior credible interval around the median, depicted by the bold black line. Dashed line denotes no predicted growth rate effect relative to the baseline type that does not carry any of the resistance determinants studied.

**Figure S8:**
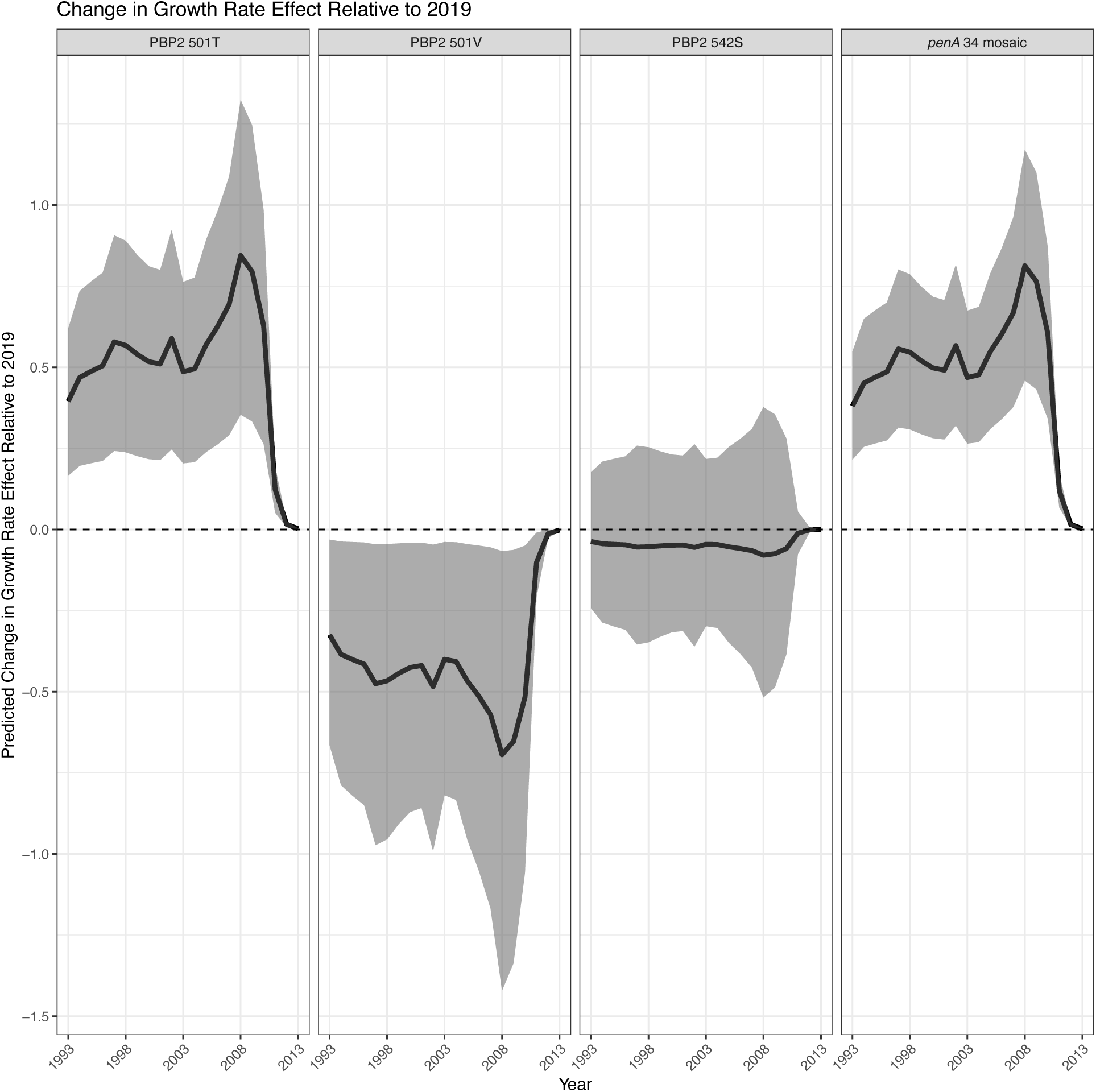
The predicted relative effect on growth rate for all determinants at the *penA* locus occurring in the dataset. The predicted effect shows how the growth rate effect of a given determinant has changed based on the reported treatments compared to that determinant’s estimated growth rate effect in 2019. Shaded region denotes the 95% posterior credible interval around the median, depicted by the bold black line. Dashed line denotes no change in predicted effect of a given determinant compared to its’ predicted effect in 2019.

**Figure S9:**
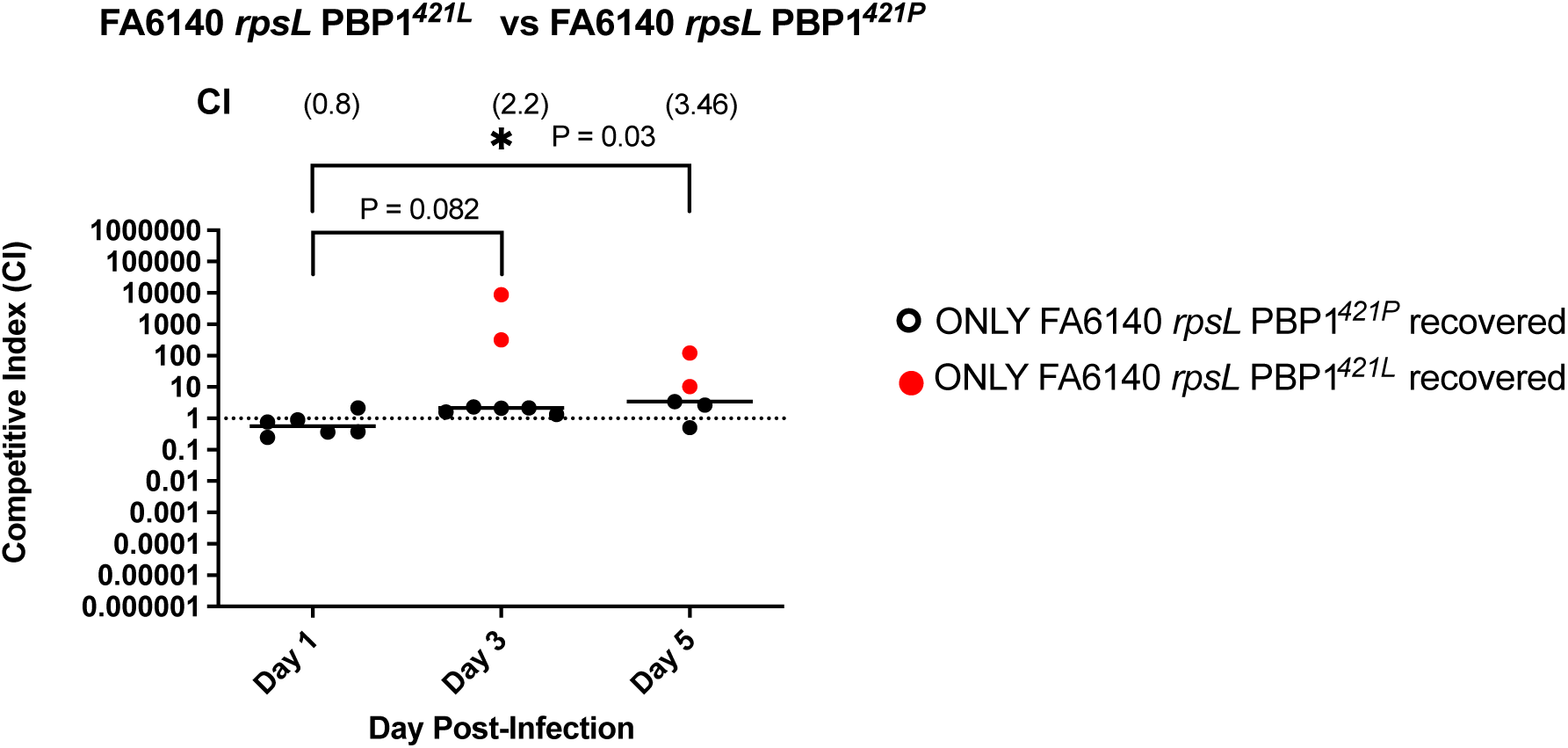
*In vivo* murine competition assay between FA6140 *rpsL* PBP1^421L^ vs FA6140 *rpsL* PBP1^421P^. Measurements were taken at three time points: 1, 3, and 5 days post-infection. N = 7 mice/time point from a single infection experiment. Each point represents the CI measurement for an individual mouse. The horizontal lines indicate the geometric mean of the CI, also shown in parantheses. Statistical significance of CI measurements was assessed using the Mann-Whitney test (\**p* < 0.05, \*\**p* < 0.005 and \*\*\**p* < 0.0005).

**Figure S10:**
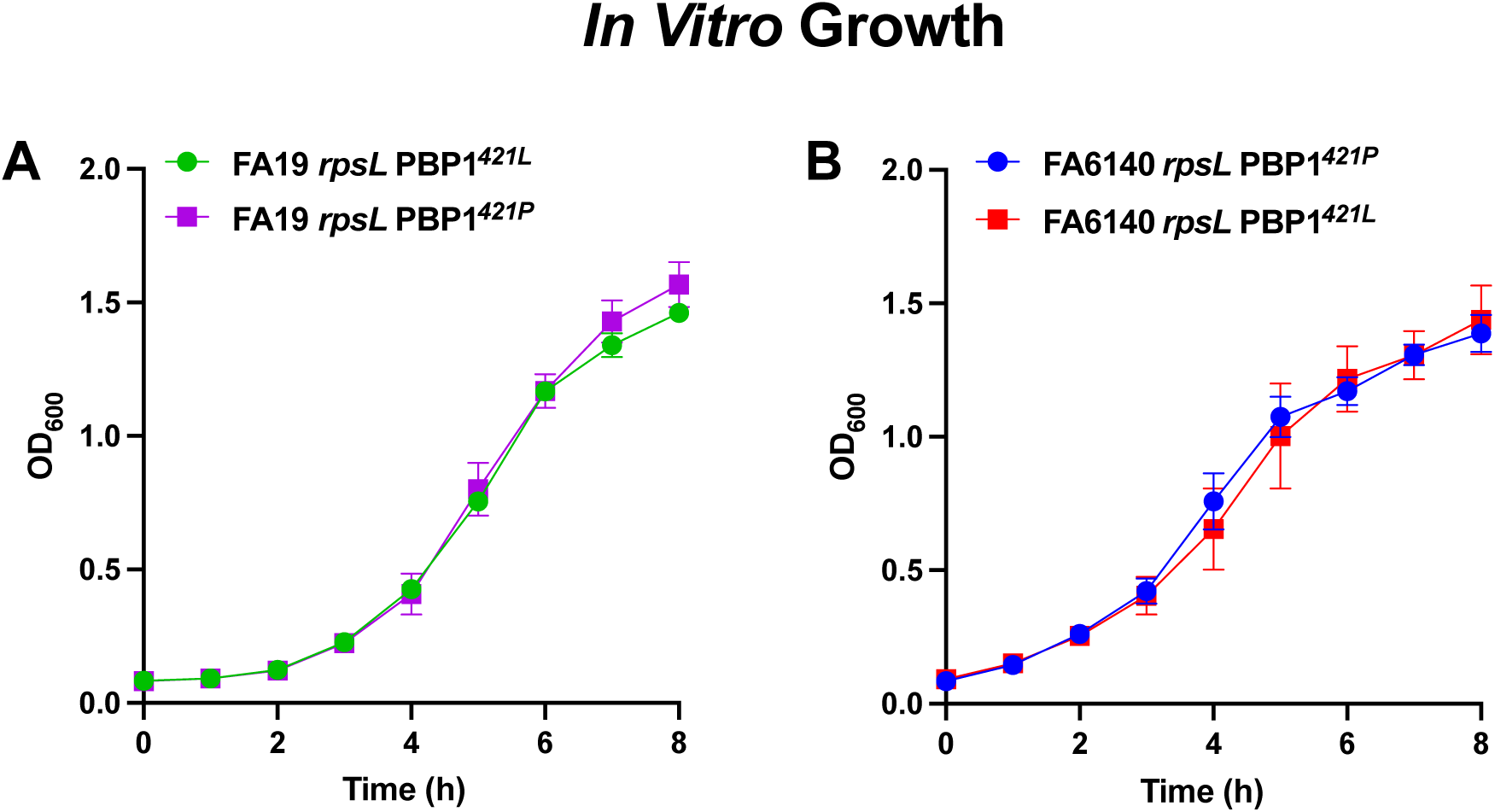
*In vitro* growth phenotypes of isogenic PBP1 421L and PBP1 421P strain pairs grown in monoculture. (A) Growth curves for strains FA19 *rpsL* and FA19 *rpsL* PBP1^421P^. (B) Growth curves for strains FA6140 *rpsL* and FA6140 *rpsL* PBP1^421L^. Data are plotted as the mean OD_600_ at each time point from three independent experiments. Statistical significance was assessed using a repeated-measures 2-way ANOVA with Tukey’s multiple comparisons. No significant differences were found.

**Figure S11:**
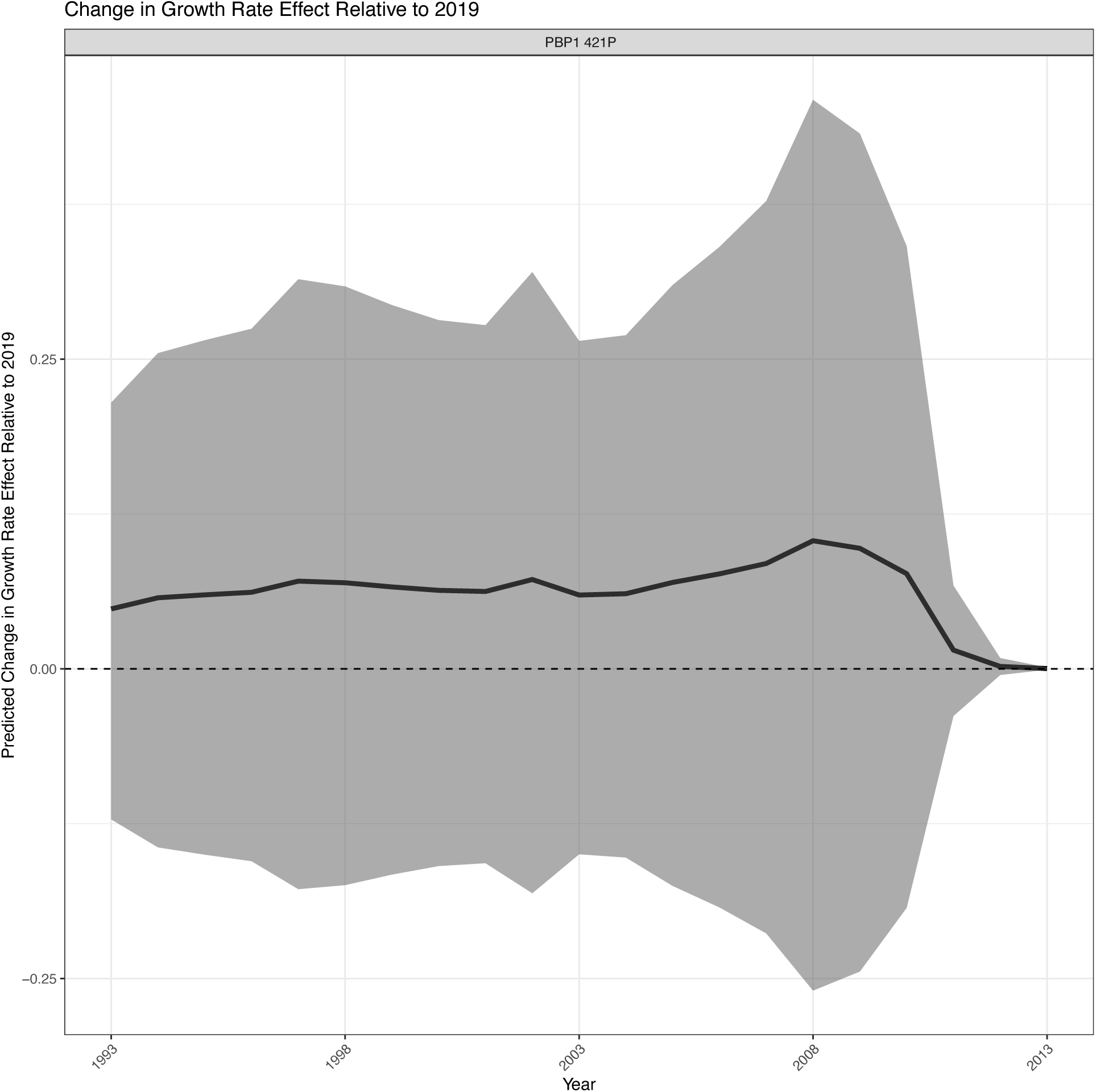
The predicted relative effect on growth rate for all determinants at the *ponA* locus occurring in the dataset. The predicted effect shows how the growth rate effect of a given determinant has changed based on the reported treatments compared to that determinant’s estimated growth rate effect in 2019. Shaded region denotes the 95% posterior credible interval around the median, depicted by the bold black line. Dashed line denotes no change in predicted effect of a given determinant compared to its’ predicted effect in 2019.

**Figure S12:**
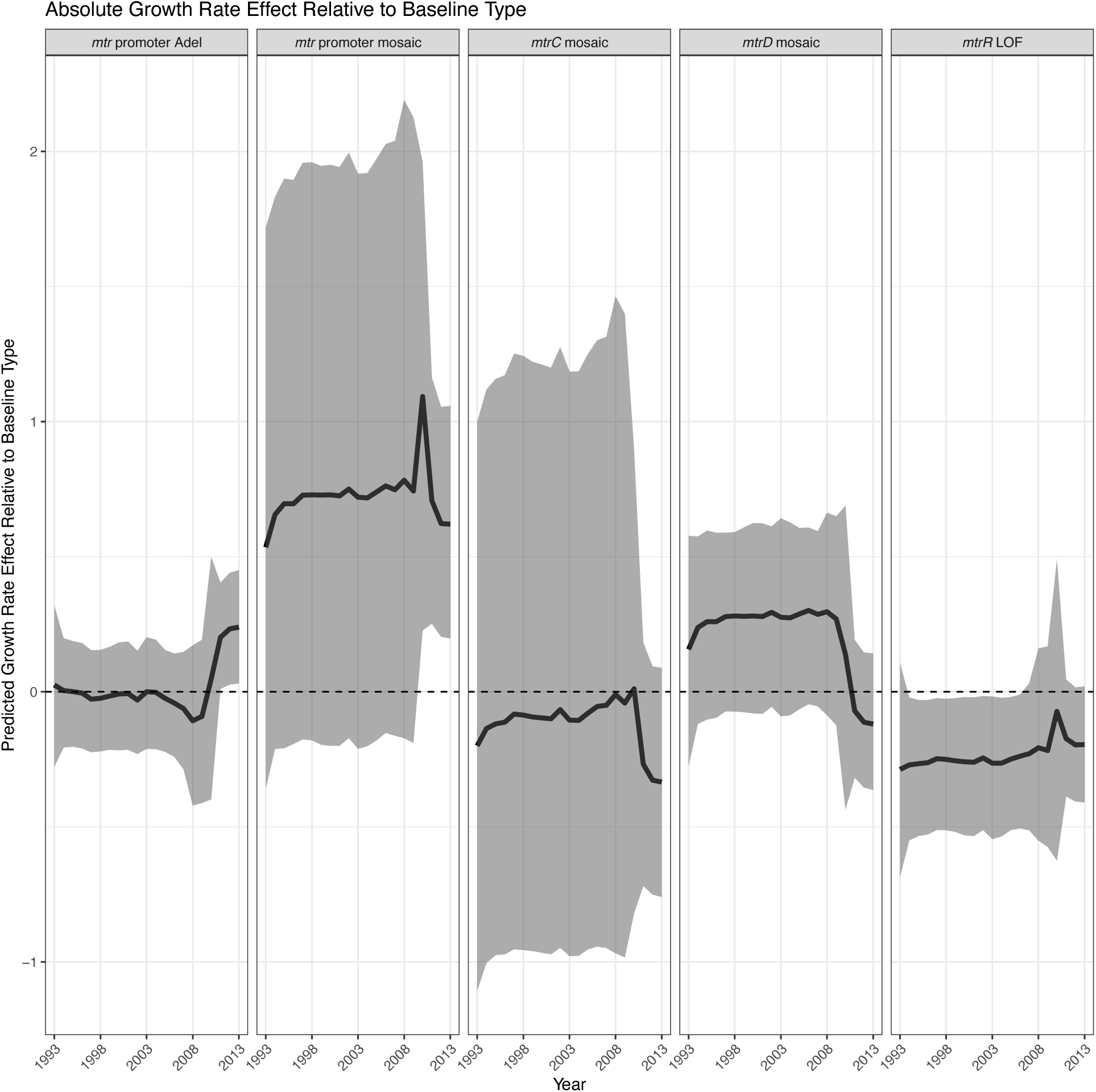
The predicted the absolute growth rate effect for all determinants at the *mtr* operon occurring in the dataset on lineage growth rate. The predicted effect was computed based on reported treatments. Shaded region denotes the 95% posterior credible interval around the median, depicted by the bold black line. Dashed line denotes no predicted growth rate effect relative to the baseline type that does not carry any of the resistance determinants studied.

**Figure S13:**
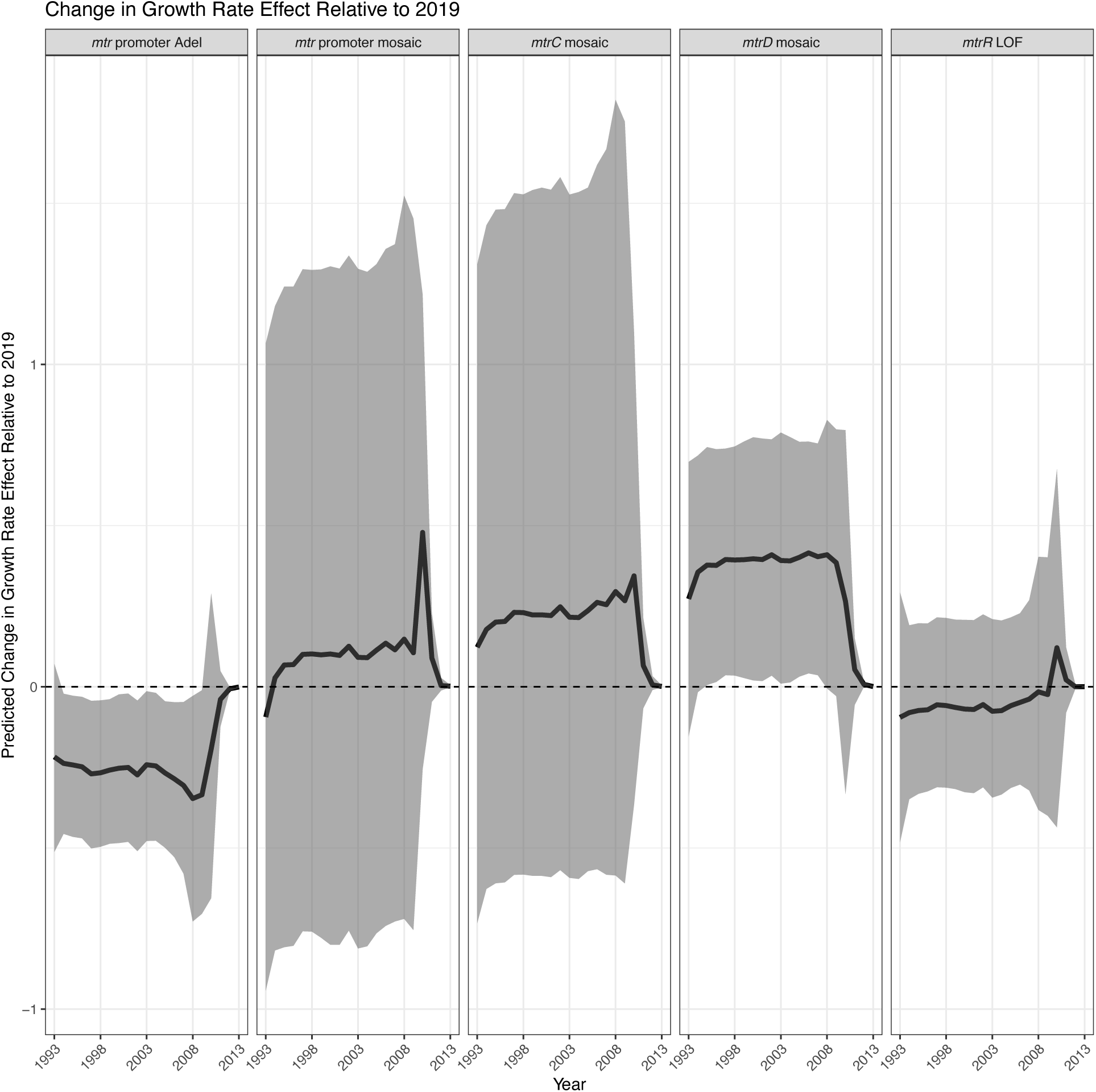
The predicted relative effect on growth rate for all determinants at the *mtr* operon occurring in the dataset. The predicted effect shows how the growth rate effect of a given determinant has changed based on the reported treatments compared to that determinant’s estimated growth rate effect in 2019. Shaded region denotes the 95% posterior credible interval around the median, depicted by the bold black line. Dashed line denotes no change in predicted effect of a given determinant compared to its’ predicted effect in 2019.

**Figure S14:**
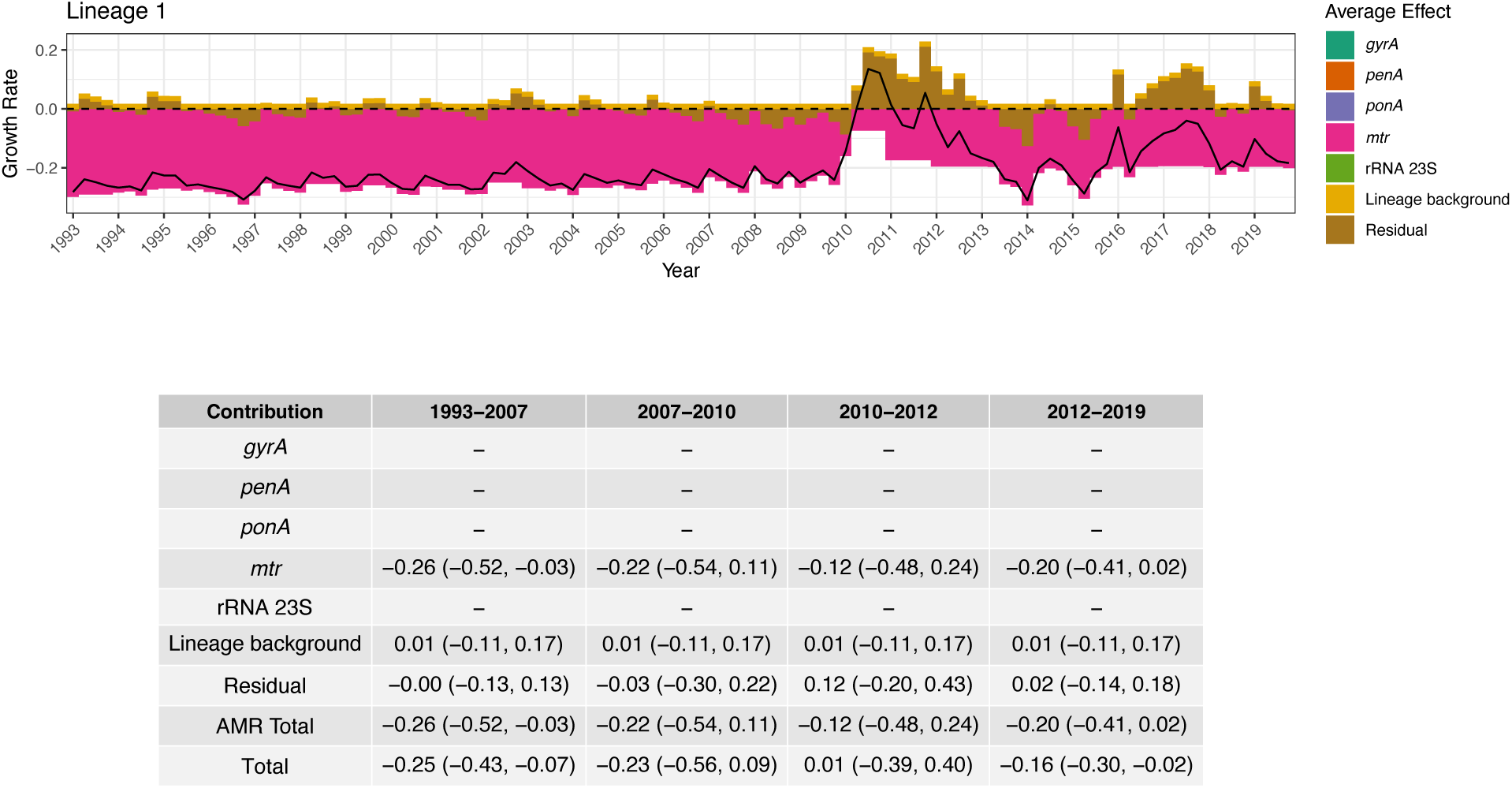
Growth rate effect summary for Lineage 1. The top panel shows the combined average growth rate effect of resistance determinants along with the lineage background term and the residual. The black solid line represents the total average effect. The dashed horizontal line indicates zero. The bottom panel depicts a table summarizing the median total growth rate effect across 4 treatment periods, as well as the 95% credible interval around the median in brackets. The period 1993-2007 corresponds to when fluoroquinolones were recommended as primary treatment; 2007-2010 to when multiple cephalosporins were recommended; 2010-2012 to when multiple cephalosporins were recommended along with azithromycin co-treatment; and 2012-2019 to when only ceftriaxone 250mg along with azithromycin co-treatment was recommended.

**Figure S15:**
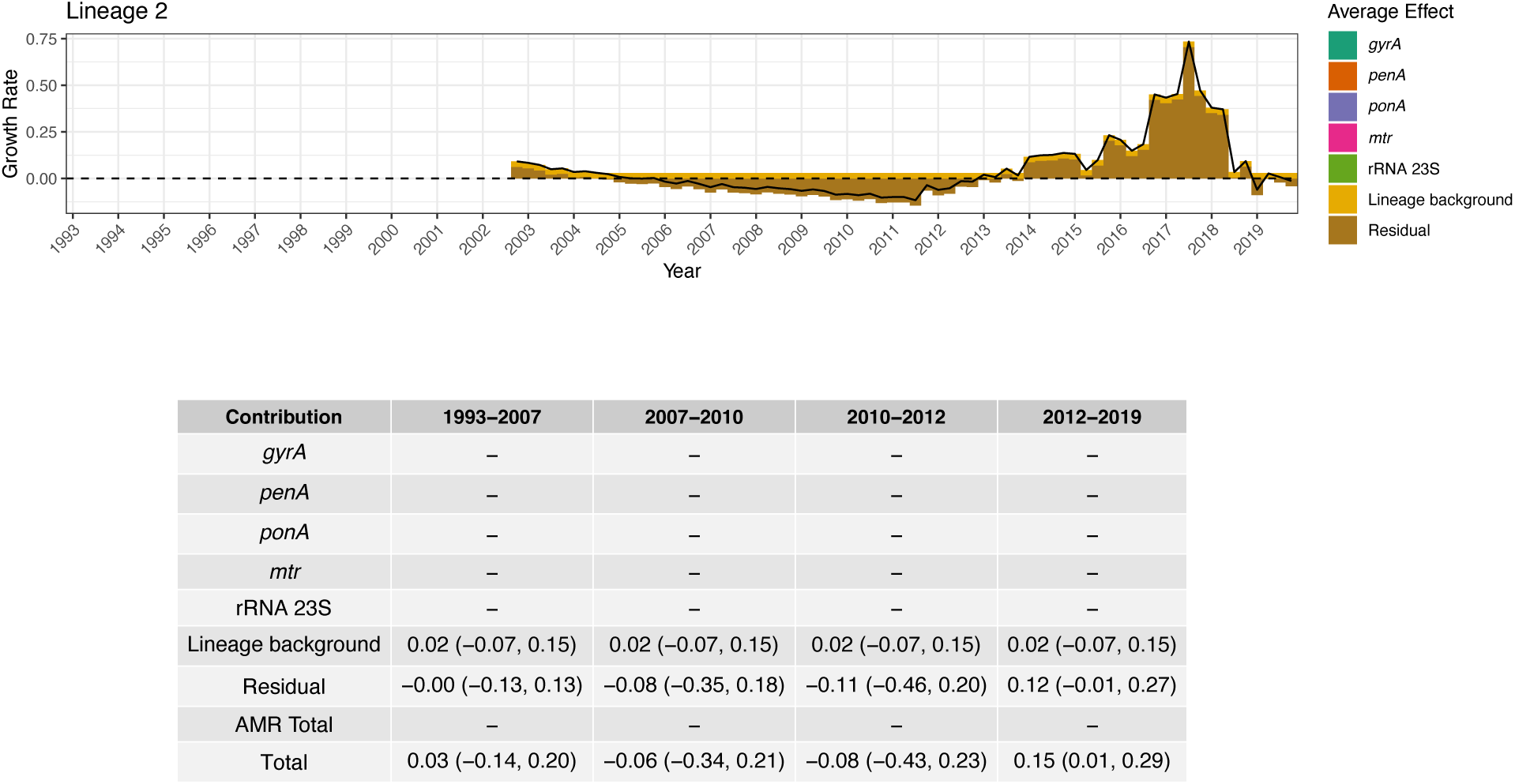
Growth rate effect summary for Lineage 2. The top panel shows the combined average growth rate effect of resistance determinants along with the lineage background term and the residual. The black solid line represents the total average effect. The dashed horizontal line indicates zero. The bottom panel depicts a table summarizing the median total growth rate effect across 4 treatment periods, as well as the 95% credible interval around the median in brackets. The period 1993-2007 corresponds to when fluoroquinolones were recommended as primary treatment; 2007-2010 to when multiple cephalosporins were recommended; 2010-2012 to when multiple cephalosporins were recommended along with azithromycin co-treatment; and 2012-2019 to when only ceftriaxone 250mg along with azithromycin co-treatment was recommended.

**Figure S16:**
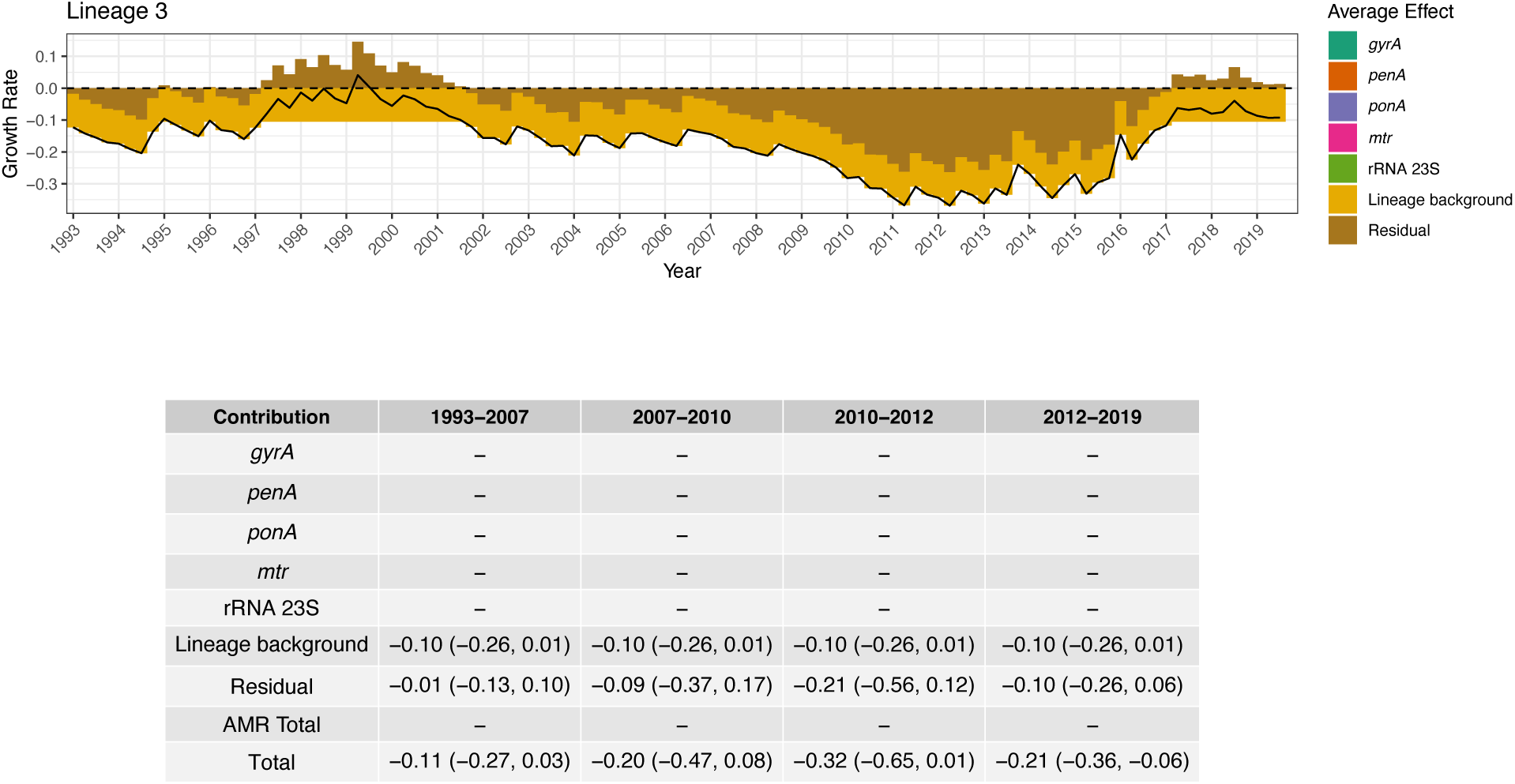
Growth rate effect summary for Lineage 3. The top panel shows the combined average growth rate effect of resistance determinants along with the lineage background term and the residual. The black solid line represents the total average effect. The dashed horizontal line indicates zero. The bottom panel depicts a table summarizing the median total growth rate effect across 4 treatment periods, as well as the 95% credible interval around the median in brackets. The period 1993-2007 corresponds to when fluoroquinolones were recommended as primary treatment; 2007-2010 to when multiple cephalosporins were recommended; 2010-2012 to when multiple cephalosporins were recommended along with azithromycin co-treatment; and 2012-2019 to when only ceftriaxone 250mg along with azithromycin co-treatment was recommended.

**Figure S17:**
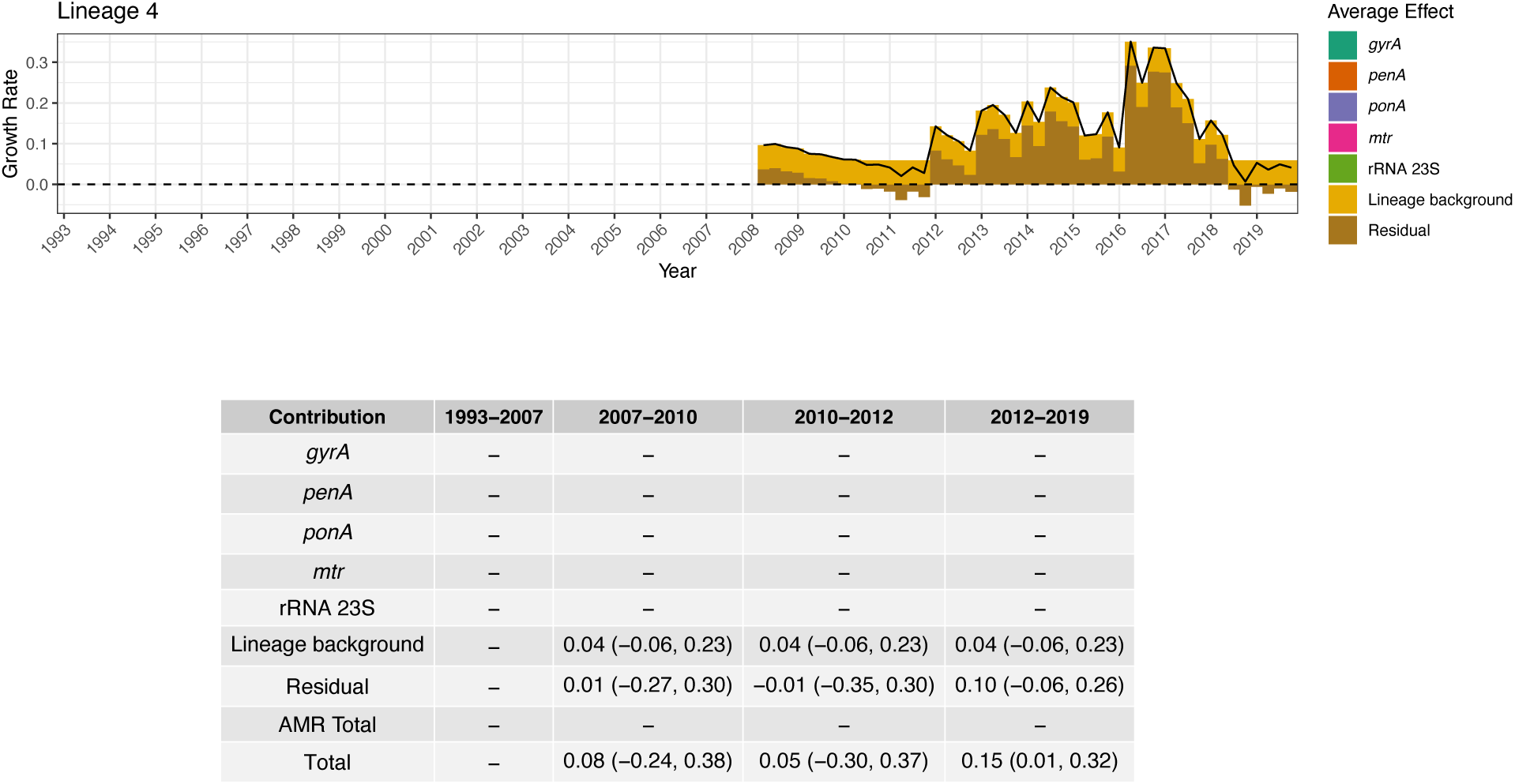
Growth rate effect summary for Lineage 4. The top panel shows the combined average growth rate effect of resistance determinants along with the lineage background term and the residual. The black solid line represents the total average effect. The dashed horizontal line indicates zero. The bottom panel depicts a table summarizing the median total growth rate effect across 4 treatment periods, as well as the 95% credible interval around the median in brackets. The period 1993-2007 corresponds to when fluoroquinolones were recommended as primary treatment; 2007-2010 to when multiple cephalosporins were recommended; 2010-2012 to when multiple cephalosporins were recommended along with azithromycin co-treatment; and 2012-2019 to when only ceftriaxone 250mg along with azithromycin co-treatment was recommended.

**Figure S18:**
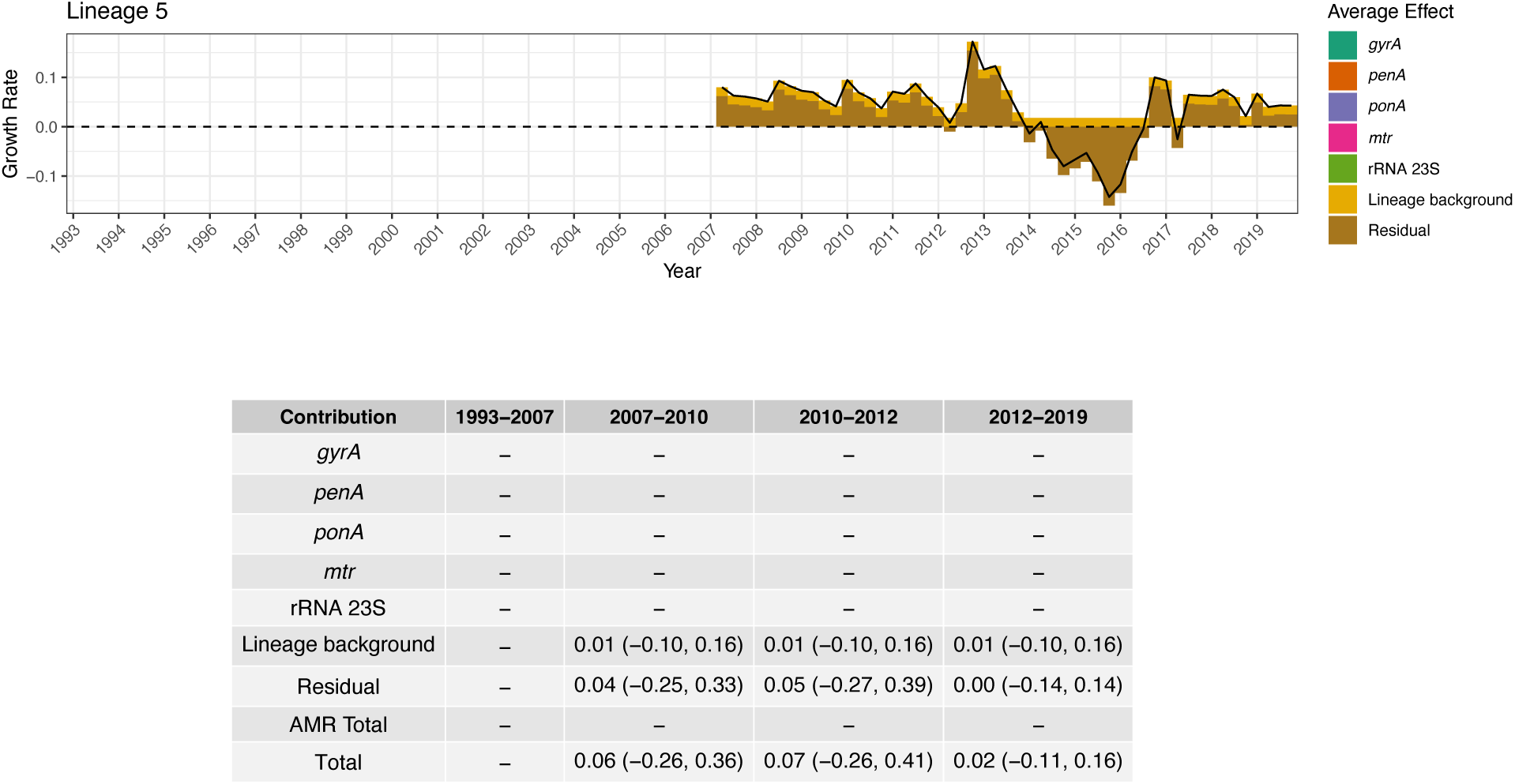
Growth rate effect summary for Lineage 5. The top panel shows the combined average growth rate effect of resistance determinants along with the lineage background term and the residual. The black solid line represents the total average effect. The dashed horizontal line indicates zero. The bottom panel depicts a table summarizing the median total growth rate effect across 4 treatment periods, as well as the 95% credible interval around the median in brackets. The period 1993-2007 corresponds to when fluoroquinolones were recommended as primary treatment; 2007-2010 to when multiple cephalosporins were recommended; 2010-2012 to when multiple cephalosporins were recommended along with azithromycin co-treatment; and 2012-2019 to when only ceftriaxone 250mg along with azithromycin co-treatment was recommended.

**Figure S19:**
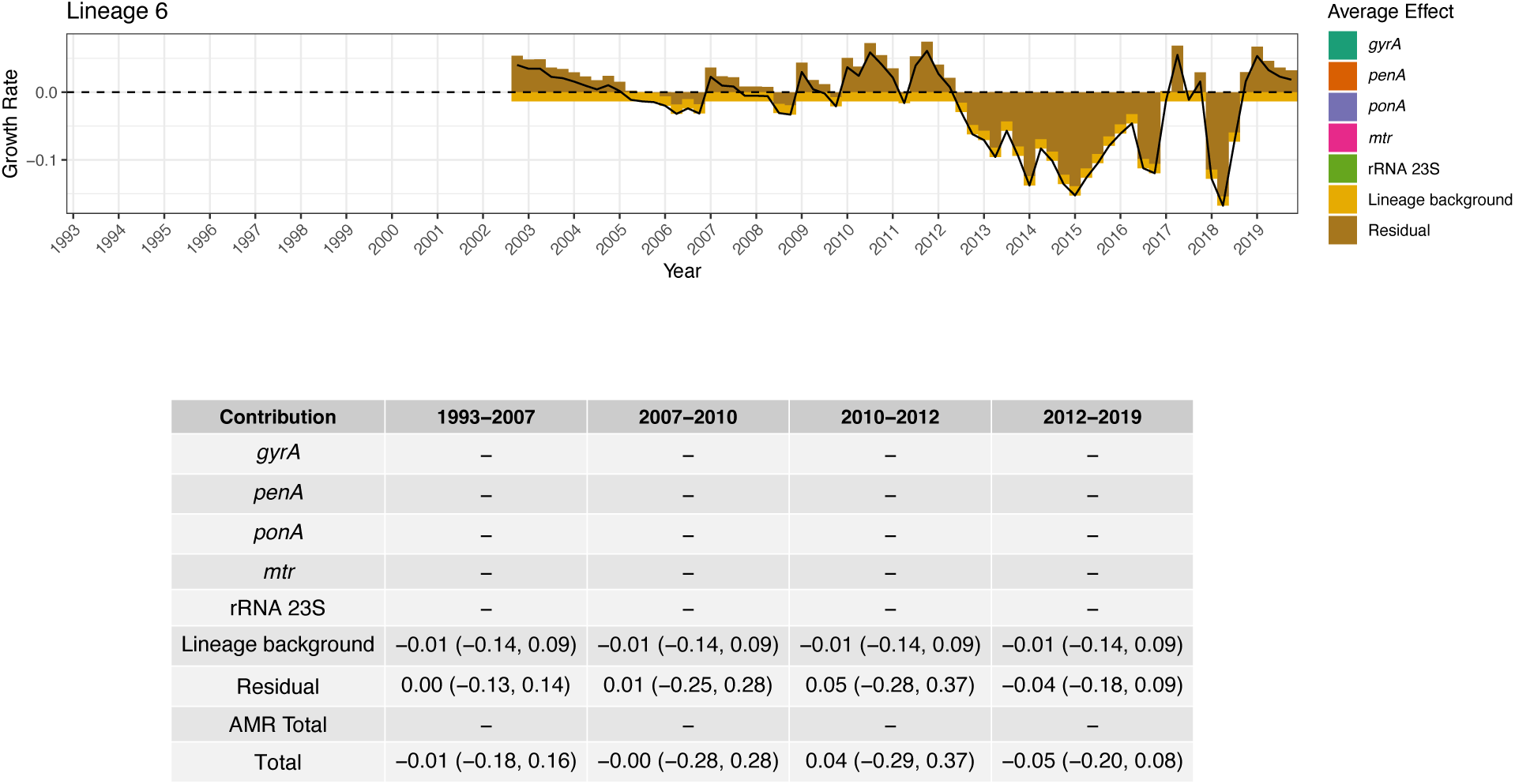
Growth rate effect summary for Lineage 6. The top panel shows the combined average growth rate effect of resistance determinants along with the lineage background term and the residual. The black solid line represents the total average effect. The dashed horizontal line indicates zero. The bottom panel depicts a table summarizing the median total growth rate effect across 4 treatment periods, as well as the 95% credible interval around the median in brackets. The period 1993-2007 corresponds to when fluoroquinolones were recommended as primary treatment; 2007-2010 to when multiple cephalosporins were recommended; 2010-2012 to when multiple cephalosporins were recommended along with azithromycin co-treatment; and 2012-2019 to when only ceftriaxone 250mg along with azithromycin co-treatment was recommended.

**Figure S20:**
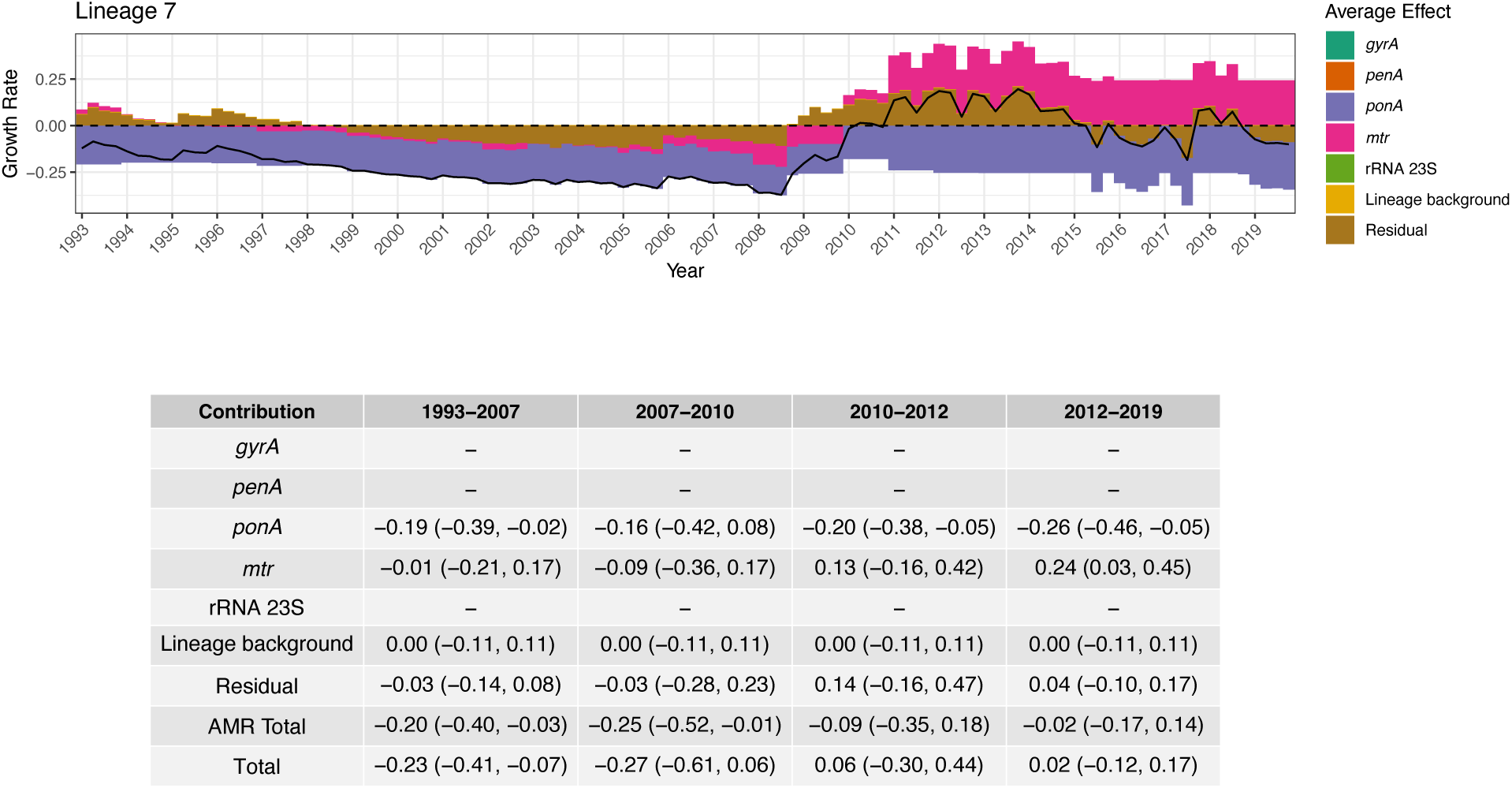
Growth rate effect summary for Lineage 7. The top panel shows the combined average growth rate effect of resistance determinants along with the lineage background term and the residual. The black solid line represents the total average effect. The dashed horizontal line indicates zero. The bottom panel depicts a table summarizing the median total growth rate effect across 4 treatment periods, as well as the 95% credible interval around the median in brackets. The period 1993-2007 corresponds to when fluoroquinolones were recommended as primary treatment; 2007-2010 to when multiple cephalosporins were recommended; 2010-2012 to when multiple cephalosporins were recommended along with azithromycin co-treatment; and 2012-2019 to when only ceftriaxone 250mg along with azithromycin co-treatment was recommended.

**Figure S21:**
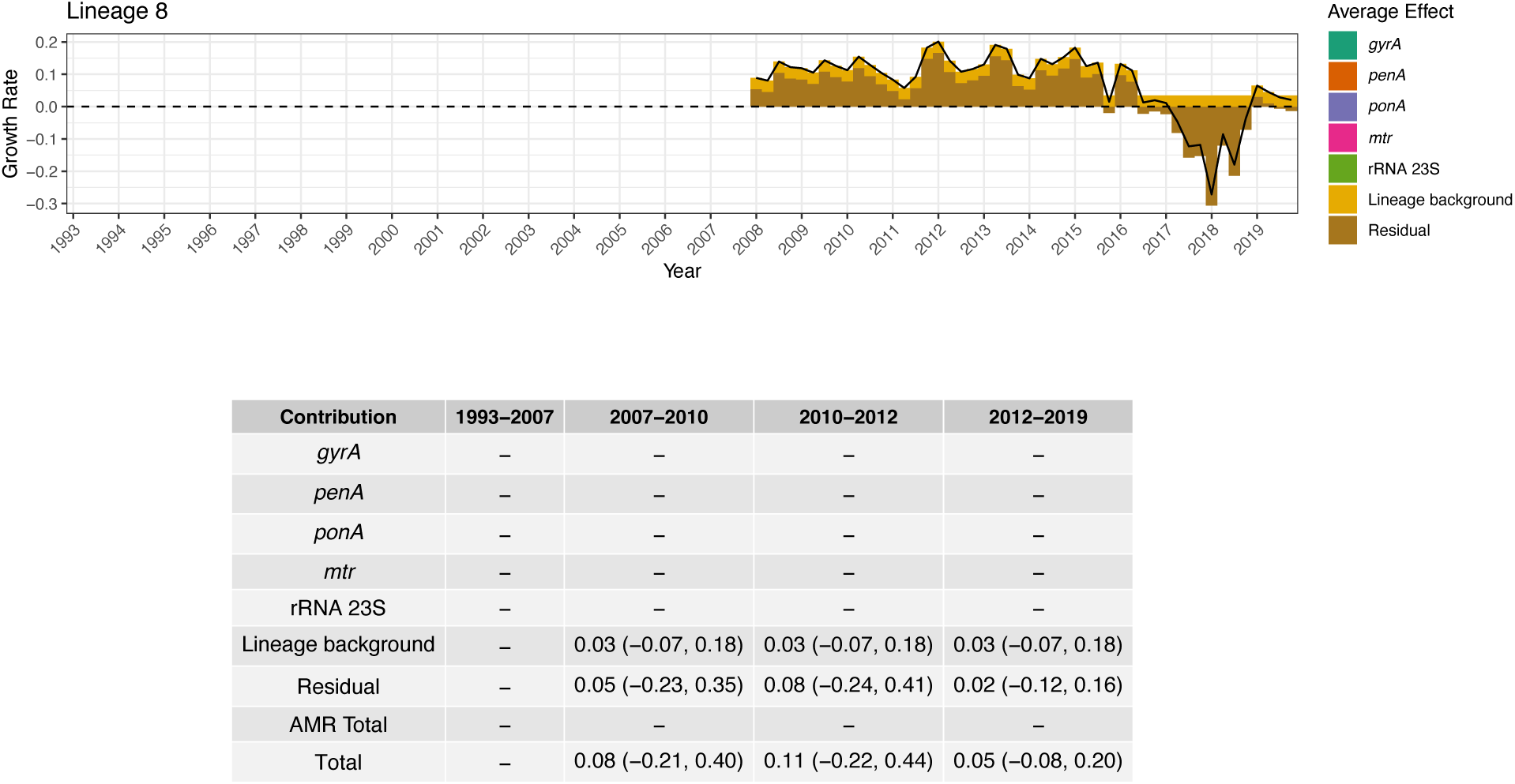
Growth rate effect summary for Lineage 8. The top panel shows the combined average growth rate effect of resistance determinants along with the lineage background term and the residual. The black solid line represents the total average effect. The dashed horizontal line indicates zero. The bottom panel depicts a table summarizing the median total growth rate effect across 4 treatment periods, as well as the 95% credible interval around the median in brackets. The period 1993-2007 corresponds to when fluoroquinolones were recommended as primary treatment; 2007-2010 to when multiple cephalosporins were recommended; 2010-2012 to when multiple cephalosporins were recommended along with azithromycin co-treatment; and 2012-2019 to when only ceftriaxone 250mg along with azithromycin co-treatment was recommended.

**Figure S22:**
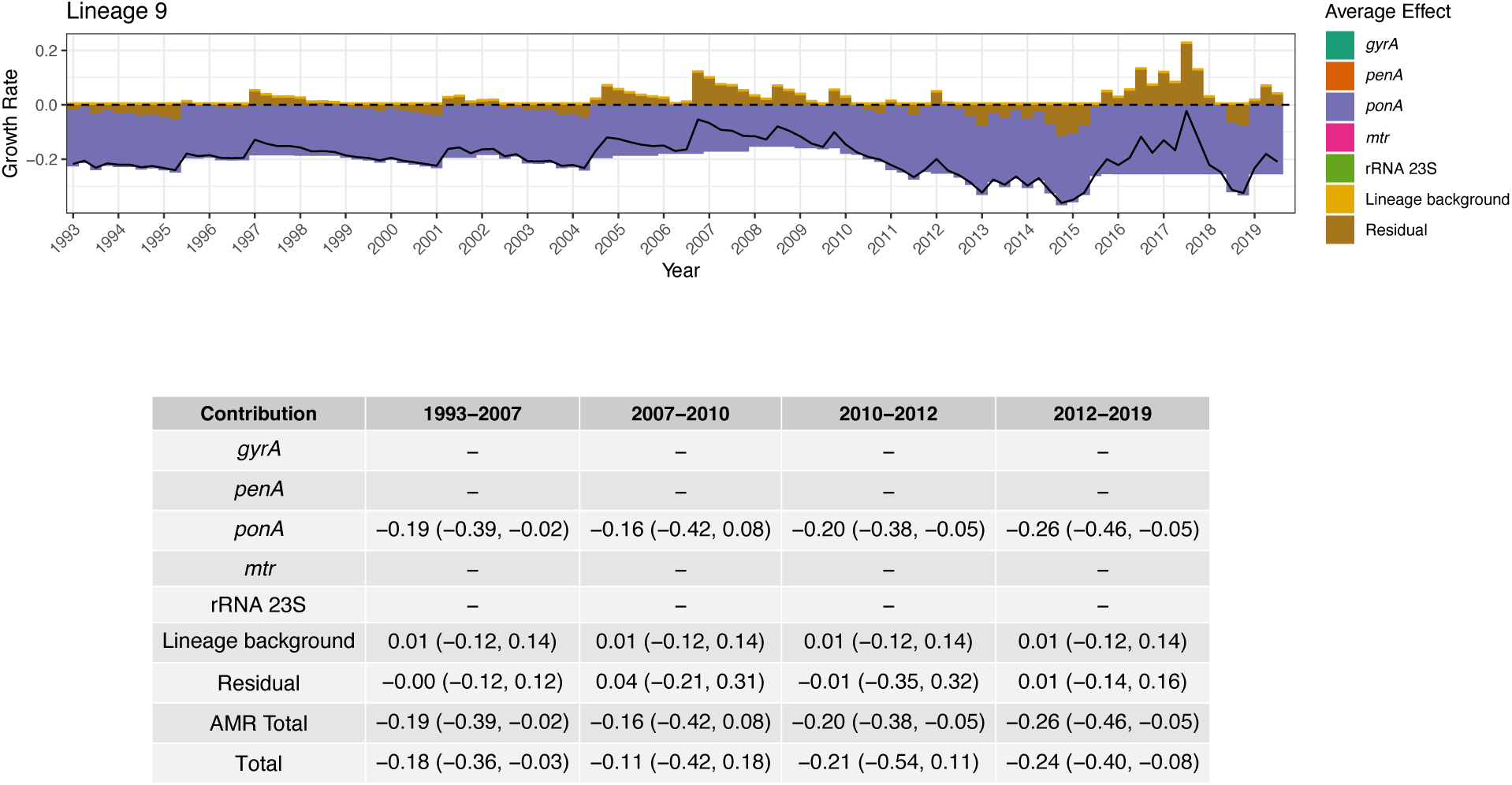
Growth rate effect summary for Lineage 9. The top panel shows the combined average growth rate effect of resistance determinants along with the lineage background term and the residual. The black solid line represents the total average effect. The dashed horizontal line indicates zero. The bottom panel depicts a table summarizing the median total growth rate effect across 4 treatment periods, as well as the 95% credible interval around the median in brackets. The period 1993-2007 corresponds to when fluoroquinolones were recommended as primary treatment; 2007-2010 to when multiple cephalosporins were recommended; 2010-2012 to when multiple cephalosporins were recommended along with azithromycin co-treatment; and 2012-2019 to when only ceftriaxone 250mg along with azithromycin co-treatment was recommended.

**Figure S23:**
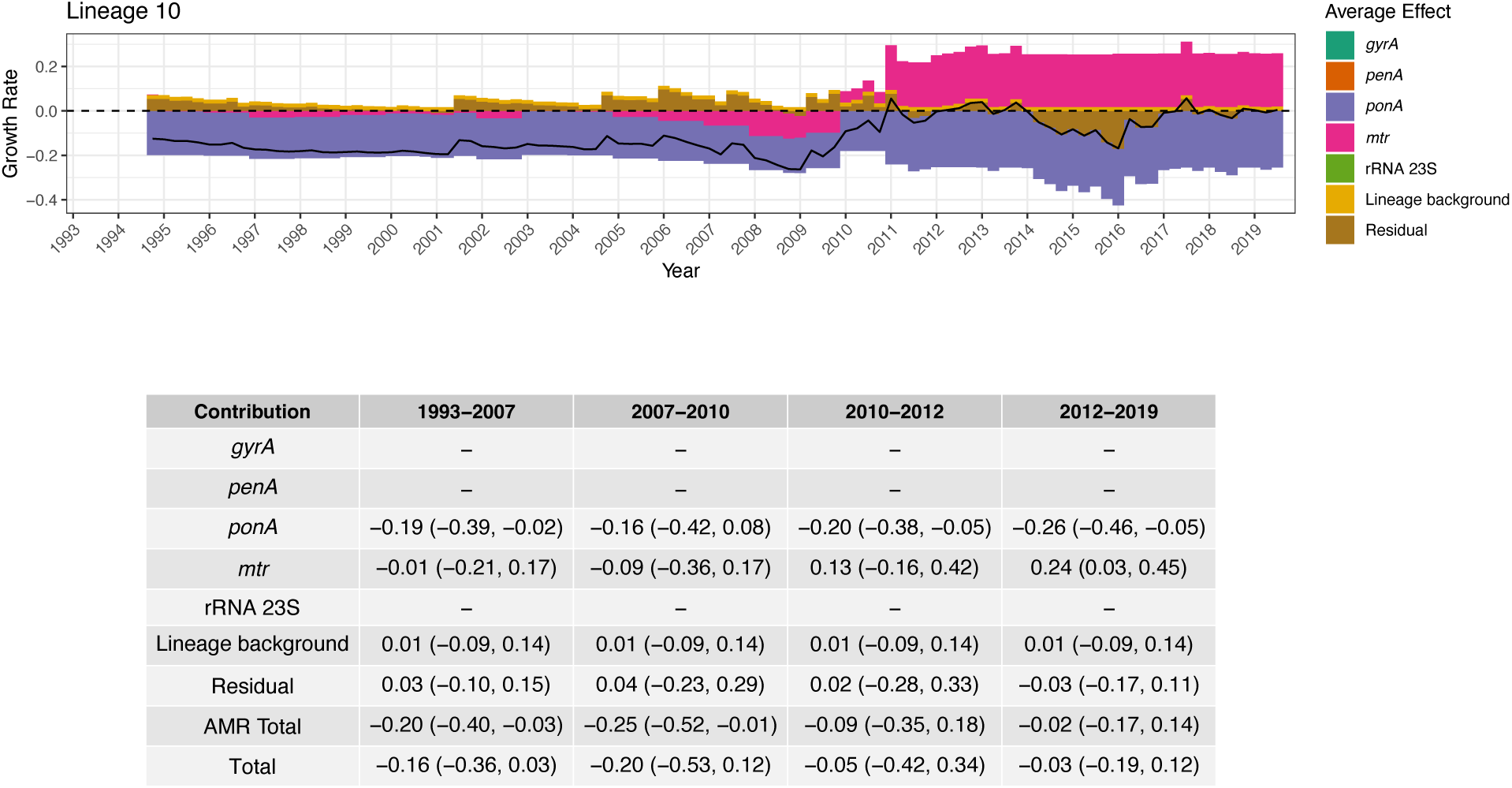
Growth rate effect summary for Lineage 10. The top panel shows the combined average growth rate effect of resistance determinants along with the lineage background term and the residual. The black solid line represents the total average effect. The dashed horizontal line indicates zero. The bottom panel depicts a table summarizing the median total growth rate effect across 4 treatment periods, as well as the 95% credible interval around the median in brackets. The period 1993-2007 corresponds to when fluoroquinolones were recommended as primary treatment; 2007-2010 to when multiple cephalosporins were recommended; 2010-2012 to when multiple cephalosporins were recommended along with azithromycin co-treatment; and 2012-2019 to when only ceftriaxone 250mg along with azithromycin co-treatment was recommended.

**Figure S24:**
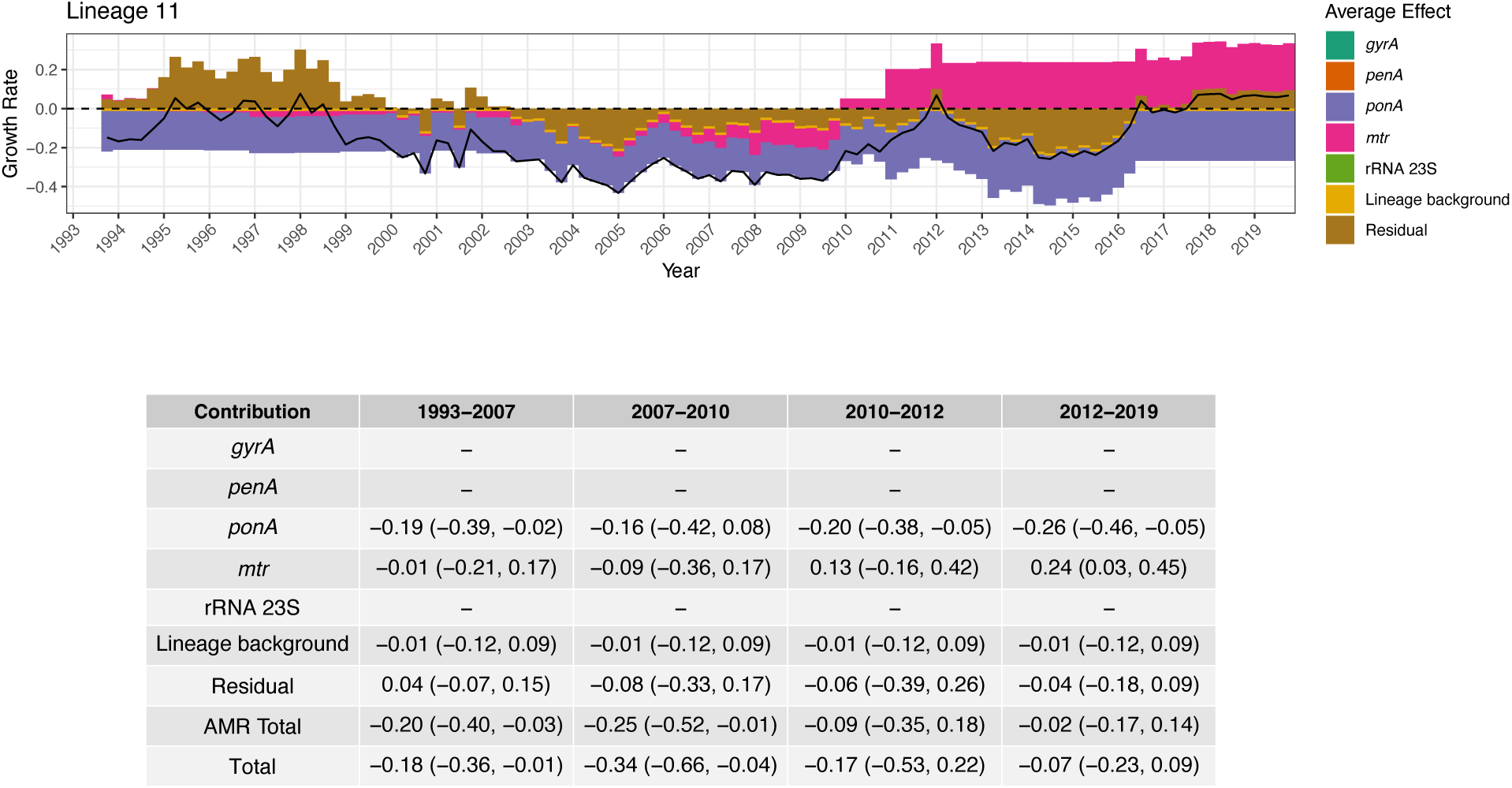
Growth rate effect summary for Lineage 11. The top panel shows the combined average growth rate effect of resistance determinants along with the lineage background term and the residual. The black solid line represents the total average effect. The dashed horizontal line indicates zero. The bottom panel depicts a table summarizing the median total growth rate effect across 4 treatment periods, as well as the 95% credible interval around the median in brackets. The period 1993-2007 corresponds to when fluoroquinolones were recommended as primary treatment; 2007-2010 to when multiple cephalosporins were recommended; 2010-2012 to when multiple cephalosporins were recommended along with azithromycin co-treatment; and 2012-2019 to when only ceftriaxone 250mg along with azithromycin co-treatment was recommended.

**Figure S25:**
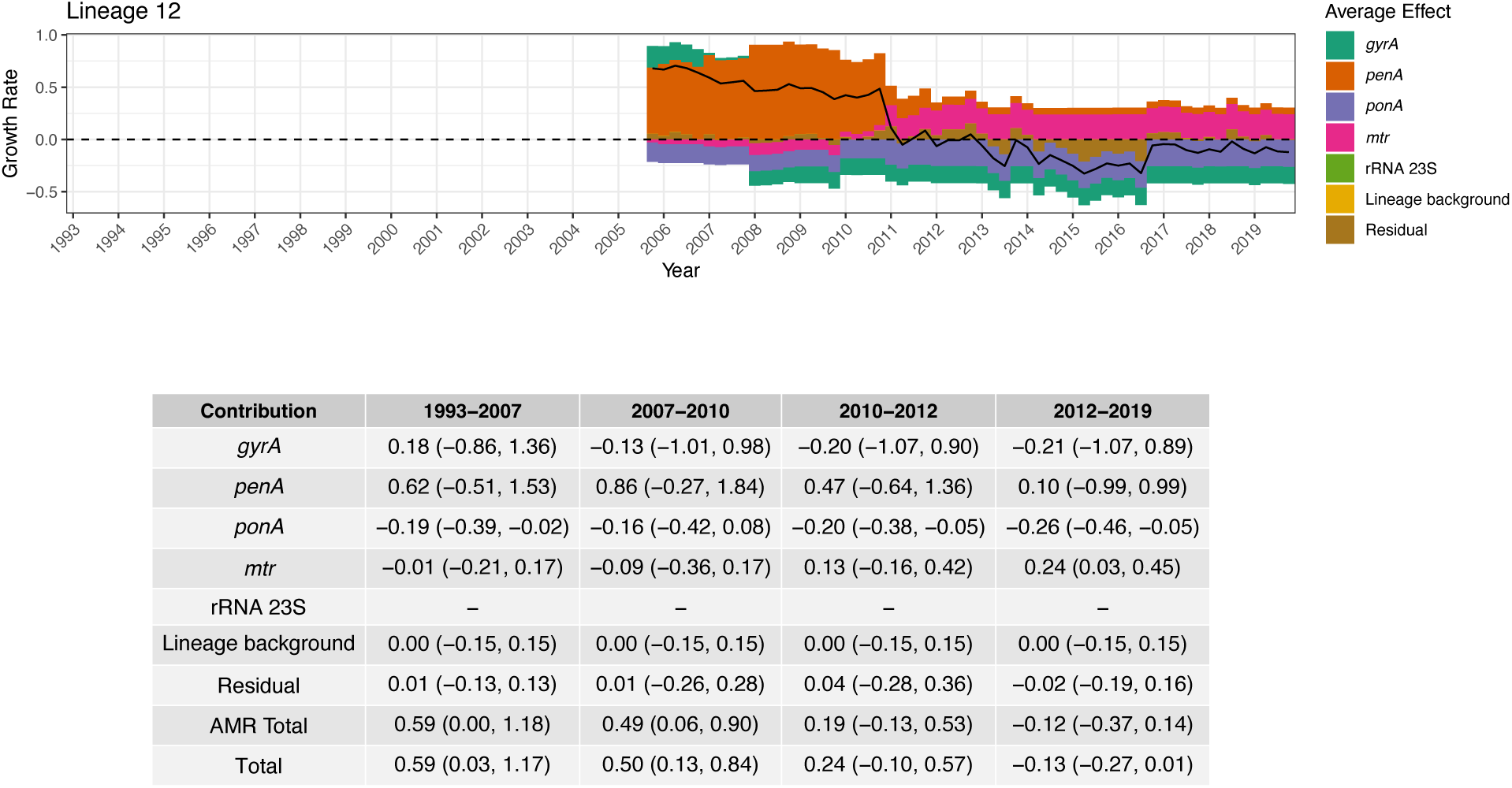
Growth rate effect summary for Lineage 12. The top panel shows the combined average growth rate effect of resistance determinants along with the lineage background term and the residual. The black solid line represents the total average effect. The dashed horizontal line indicates zero. The bottom panel depicts a table summarizing the median total growth rate effect across 4 treatment periods, as well as the 95% credible interval around the median in brackets. The period 1993-2007 corresponds to when fluoroquinolones were recommended as primary treatment; 2007-2010 to when multiple cephalosporins were recommended; 2010-2012 to when multiple cephalosporins were recommended along with azithromycin co-treatment; and 2012-2019 to when only ceftriaxone 250mg along with azithromycin co-treatment was recommended.

**Figure S26:**
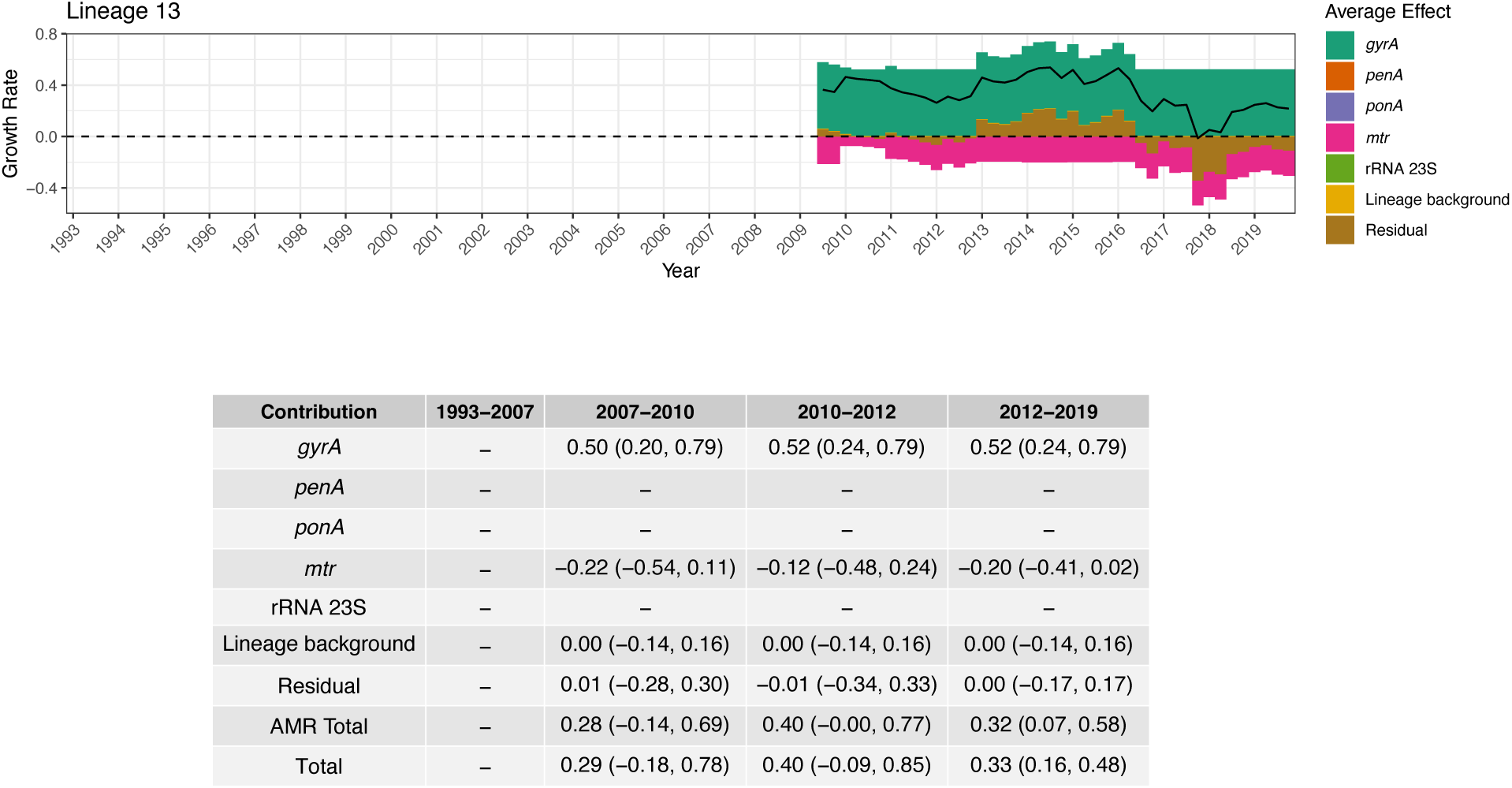
Growth rate effect summary for Lineage 13. The top panel shows the combined average growth rate effect of resistance determinants along with the lineage background term and the residual. The black solid line represents the total average effect. The dashed horizontal line indicates zero. The bottom panel depicts a table summarizing the median total growth rate effect across 4 treatment periods, as well as the 95% credible interval around the median in brackets. The period 1993-2007 corresponds to when fluoroquinolones were recommended as primary treatment; 2007-2010 to when multiple cephalosporins were recommended; 2010-2012 to when multiple cephalosporins were recommended along with azithromycin co-treatment; and 2012-2019 to when only ceftriaxone 250mg along with azithromycin co-treatment was recommended.

**Figure S27:**
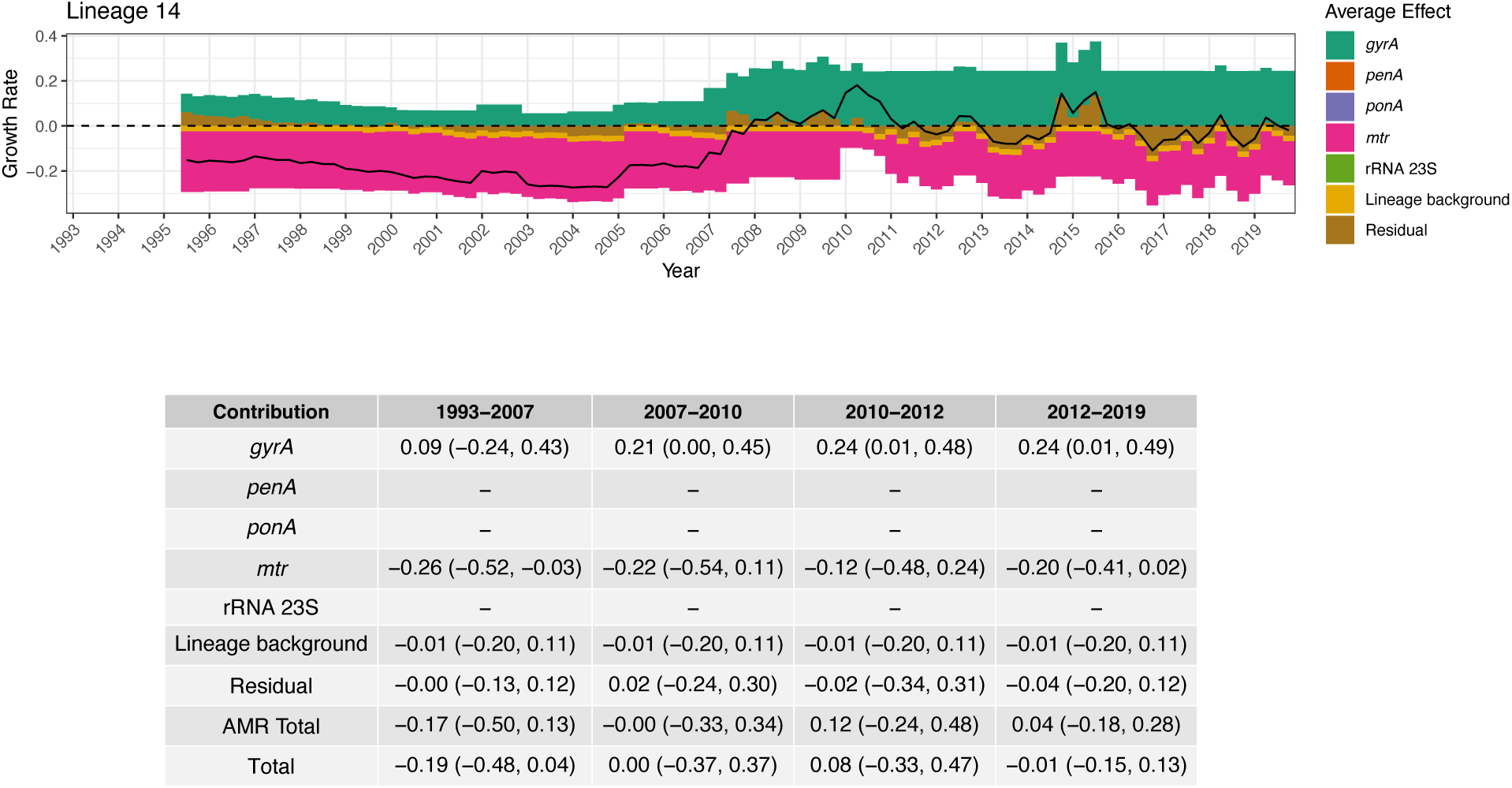
Growth rate effect summary for Lineage 14. The top panel shows the combined average growth rate effect of resistance determinants along with the lineage background term and the residual. The black solid line represents the total average effect. The dashed horizontal line indicates zero. The bottom panel depicts a table summarizing the median total growth rate effect across 4 treatment periods, as well as the 95% credible interval around the median in brackets. The period 1993-2007 corresponds to when fluoroquinolones were recommended as primary treatment; 2007-2010 to when multiple cephalosporins were recommended; 2010-2012 to when multiple cephalosporins were recommended along with azithromycin co-treatment; and 2012-2019 to when only ceftriaxone 250mg along with azithromycin co-treatment was recommended.

**Figure S28:**
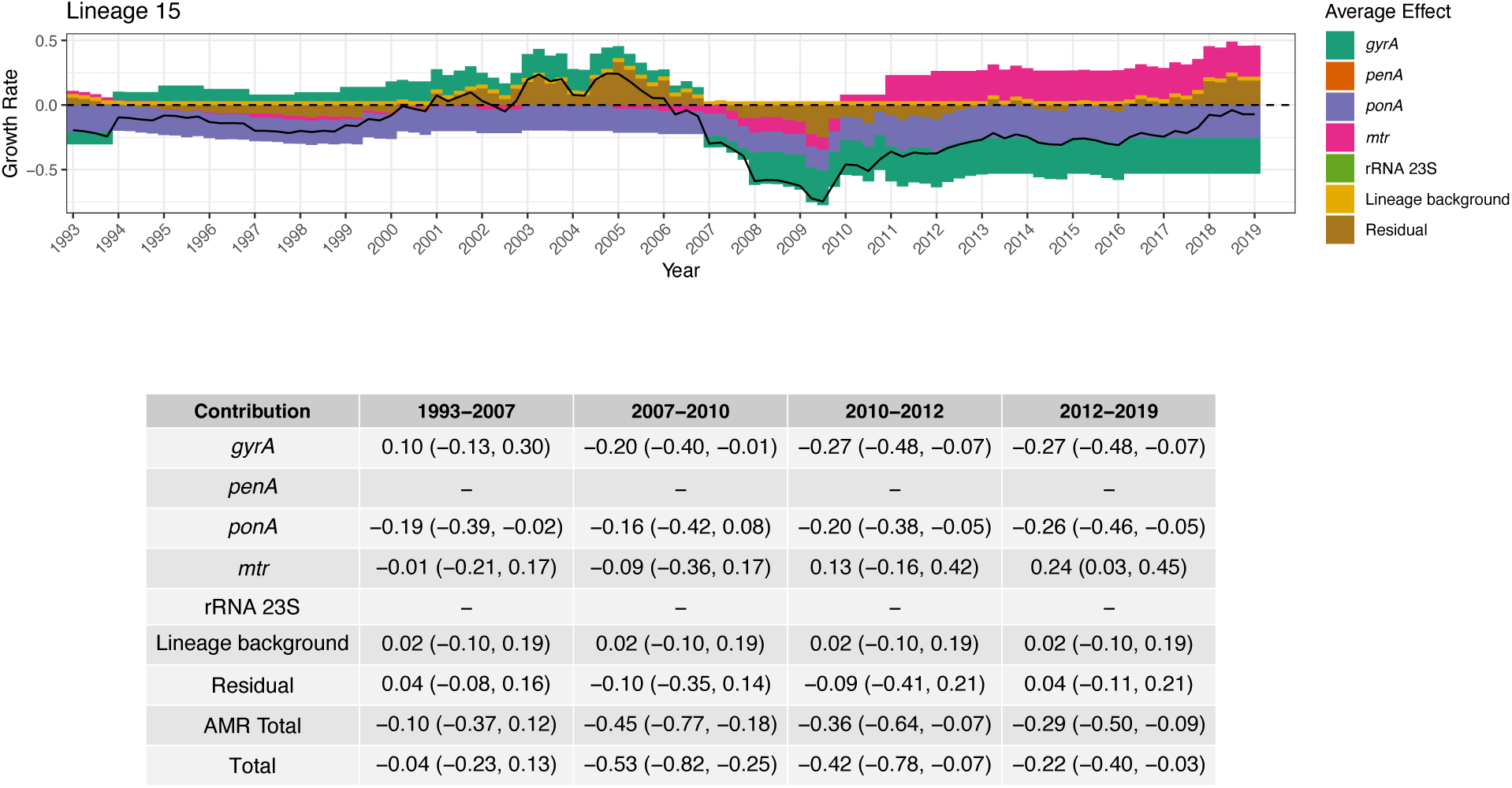
Growth rate effect summary for Lineage 15. The top panel shows the combined average growth rate effect of resistance determinants along with the lineage background term and the residual. The black solid line represents the total average effect. The dashed horizontal line indicates zero. The bottom panel depicts a table summarizing the median total growth rate effect across 4 treatment periods, as well as the 95% credible interval around the median in brackets. The period 1993-2007 corresponds to when fluoroquinolones were recommended as primary treatment; 2007-2010 to when multiple cephalosporins were recommended; 2010-2012 to when multiple cephalosporins were recommended along with azithromycin co-treatment; and 2012-2019 to when only ceftriaxone 250mg along with azithromycin co-treatment was recommended.

**Figure S29:**
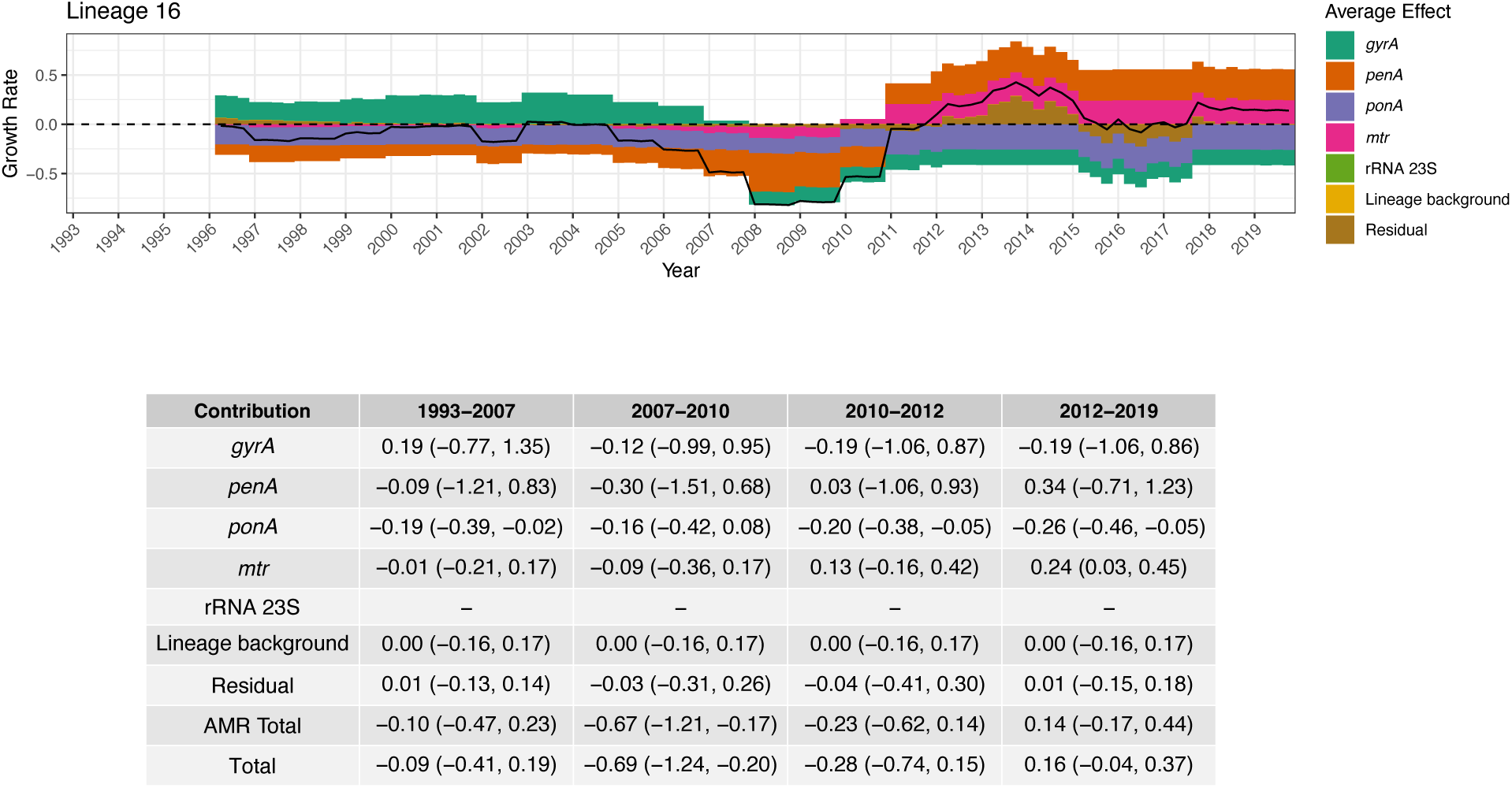
Growth rate effect summary for Lineage 1. The top panel shows the combined average growth rate effect of resistance determinants along with the lineage background term and the residual. The black solid line represents the total average effect. The dashed horizontal line indicates zero. The bottom panel depicts a table summarizing the median total growth rate effect across 4 treatment periods, as well as the 95% credible interval around the median in brackets. The period 1993-2007 corresponds to when fluoroquinolones were recommended as primary treatment; 2007-2010 to when multiple cephalosporins were recommended; 2010-2012 to when multiple cephalosporins were recommended along with azithromycin co-treatment; and 2012-2019 to when only ceftriaxone 250mg along with azithromycin co-treatment was recommended.

**Figure S30:**
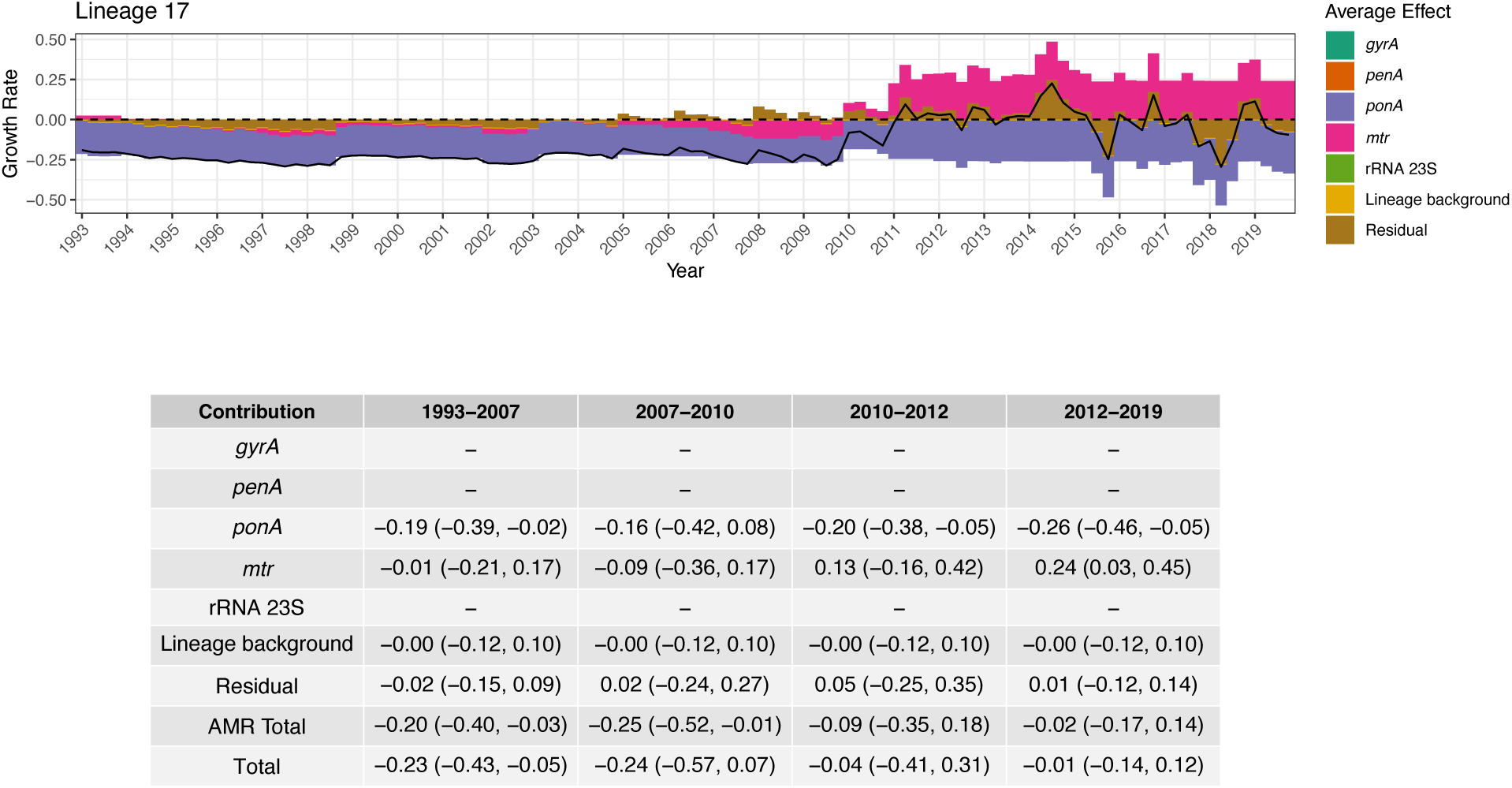
Growth rate effect summary for Lineage 17. The top panel shows the combined average growth rate effect of resistance determinants along with the lineage background term and the residual. The black solid line represents the total average effect. The dashed horizontal line indicates zero. The bottom panel depicts a table summarizing the median total growth rate effect across 4 treatment periods, as well as the 95% credible interval around the median in brackets. The period 1993-2007 corresponds to when fluoroquinolones were recommended as primary treatment; 2007-2010 to when multiple cephalosporins were recommended; 2010-2012 to when multiple cephalosporins were recommended along with azithromycin co-treatment; and 2012-2019 to when only ceftriaxone 250mg along with azithromycin co-treatment was recommended.

**Figure S31:**
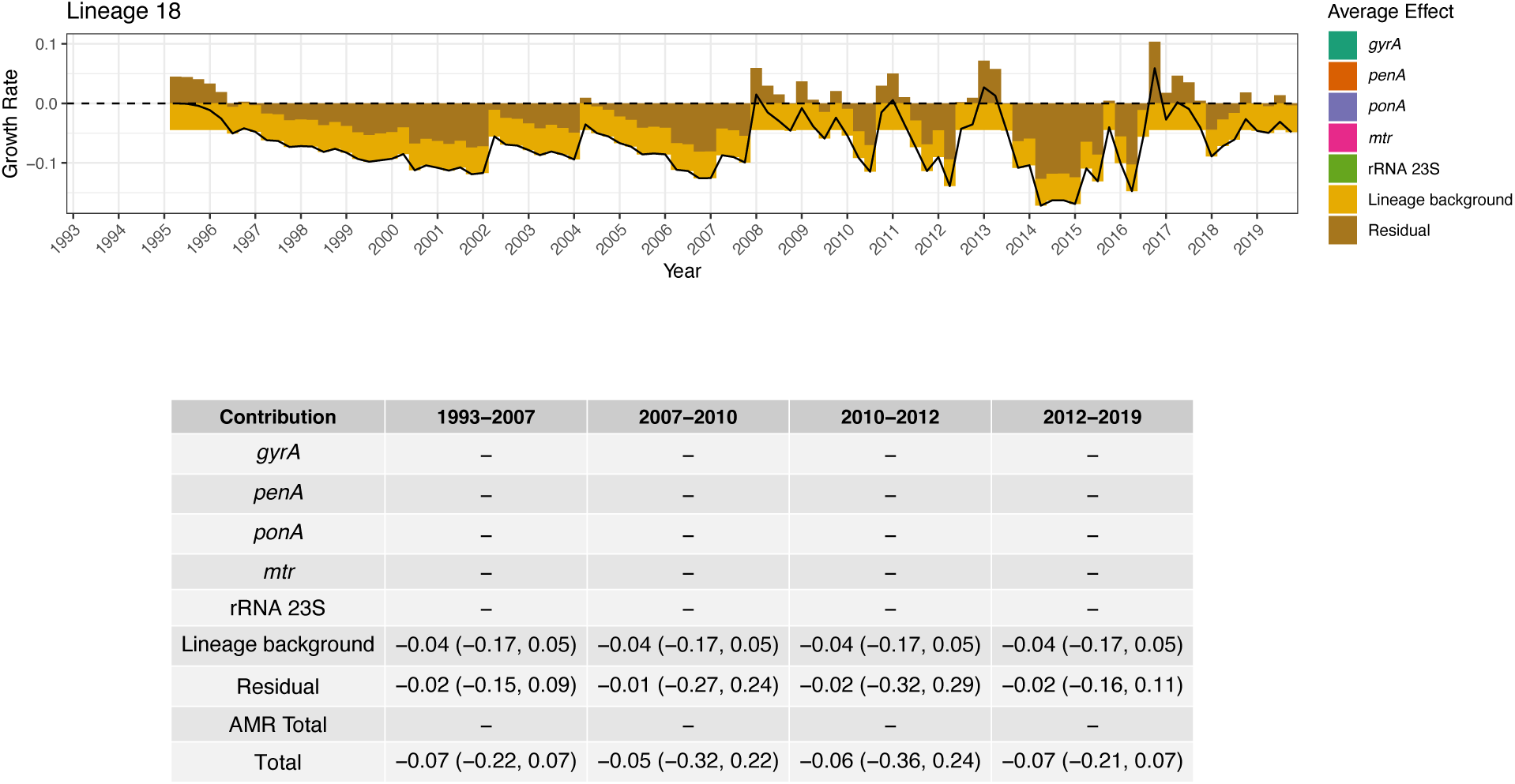
Growth rate effect summary for Lineage 18. The top panel shows the combined average growth rate effect of resistance determinants along with the lineage background term and the residual. The black solid line represents the total average effect. The dashed horizontal line indicates zero. The bottom panel depicts a table summarizing the median total growth rate effect across 4 treatment periods, as well as the 95% credible interval around the median in brackets. The period 1993-2007 corresponds to when fluoroquinolones were recommended as primary treatment; 2007-2010 to when multiple cephalosporins were recommended; 2010-2012 to when multiple cephalosporins were recommended along with azithromycin co-treatment; and 2012-2019 to when only ceftriaxone 250mg along with azithromycin co-treatment was recommended.

**Figure S32:**
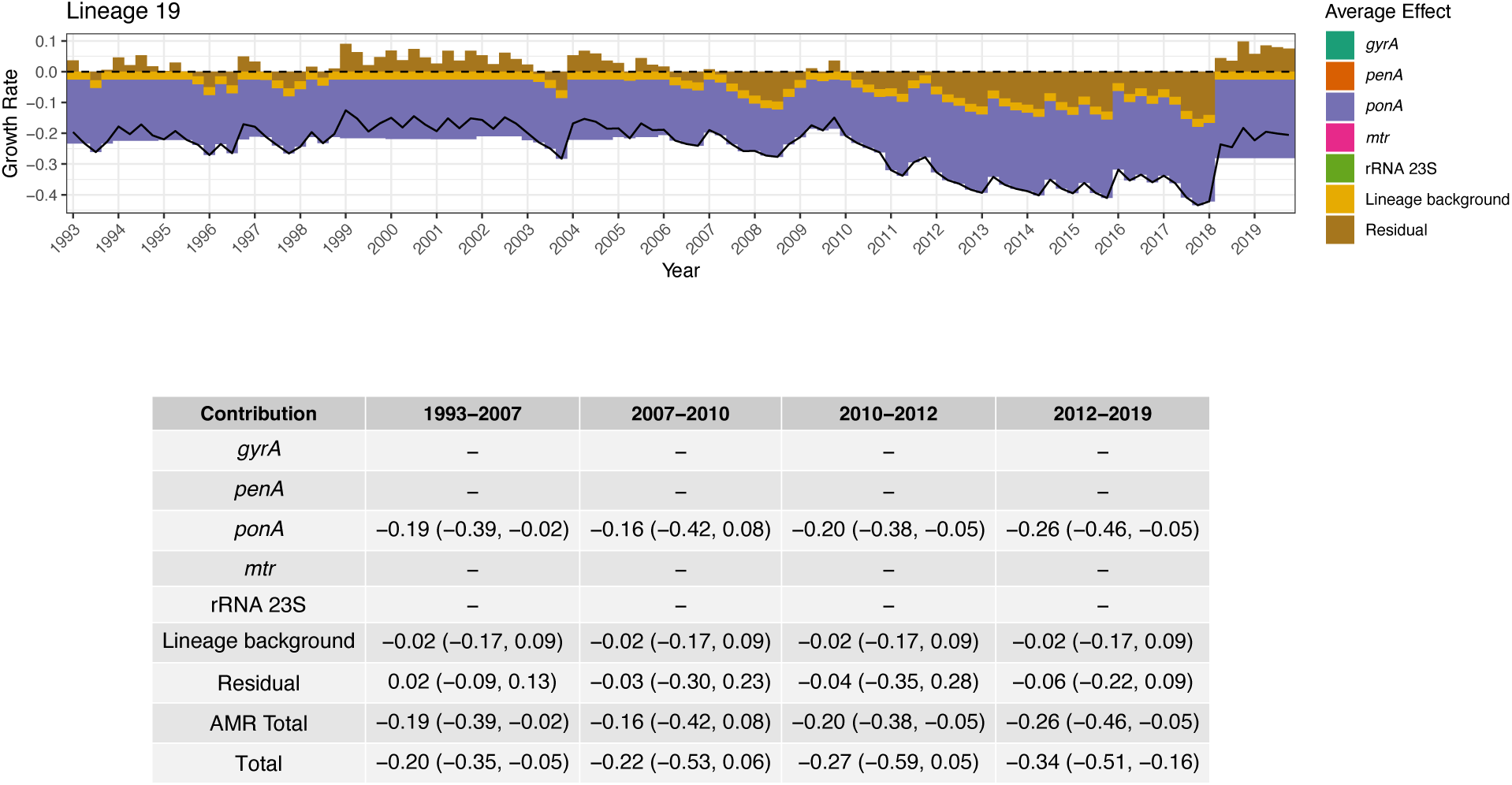
Growth rate effect summary for Lineage 19. The top panel shows the combined average growth rate effect of resistance determinants along with the lineage background term and the residual. The black solid line represents the total average effect. The dashed horizontal line indicates zero. The bottom panel depicts a table summarizing the median total growth rate effect across 4 treatment periods, as well as the 95% credible interval around the median in brackets. The period 1993-2007 corresponds to when fluoroquinolones were recommended as primary treatment; 2007-2010 to when multiple cephalosporins were recommended; 2010-2012 to when multiple cephalosporins were recommended along with azithromycin co-treatment; and 2012-2019 to when only ceftriaxone 250mg along with azithromycin co-treatment was recommended.

**Figure S33:**
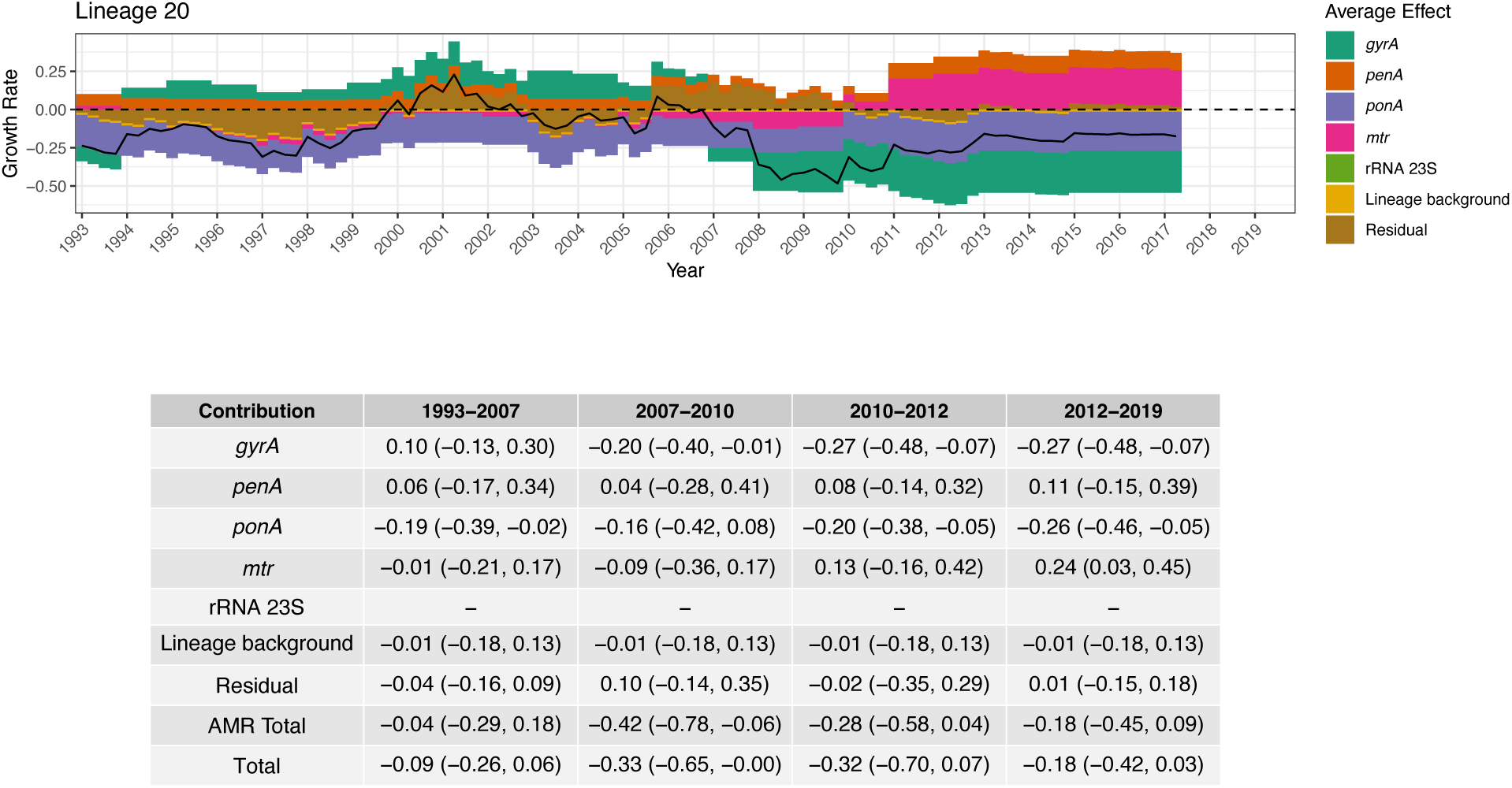
Growth rate effect summary for Lineage 20. The top panel shows the combined average growth rate effect of resistance determinants along with the lineage background term and the residual. The black solid line represents the total average effect. The dashed horizontal line indicates zero. The bottom panel depicts a table summarizing the median total growth rate effect across 4 treatment periods, as well as the 95% credible interval around the median in brackets. The period 1993-2007 corresponds to when fluoroquinolones were recommended as primary treatment; 2007-2010 to when multiple cephalosporins were recommended; 2010-2012 to when multiple cephalosporins were recommended along with azithromycin co-treatment; and 2012-2019 to when only ceftriaxone 250mg along with azithromycin co-treatment was recommended.

**Figure S34:**
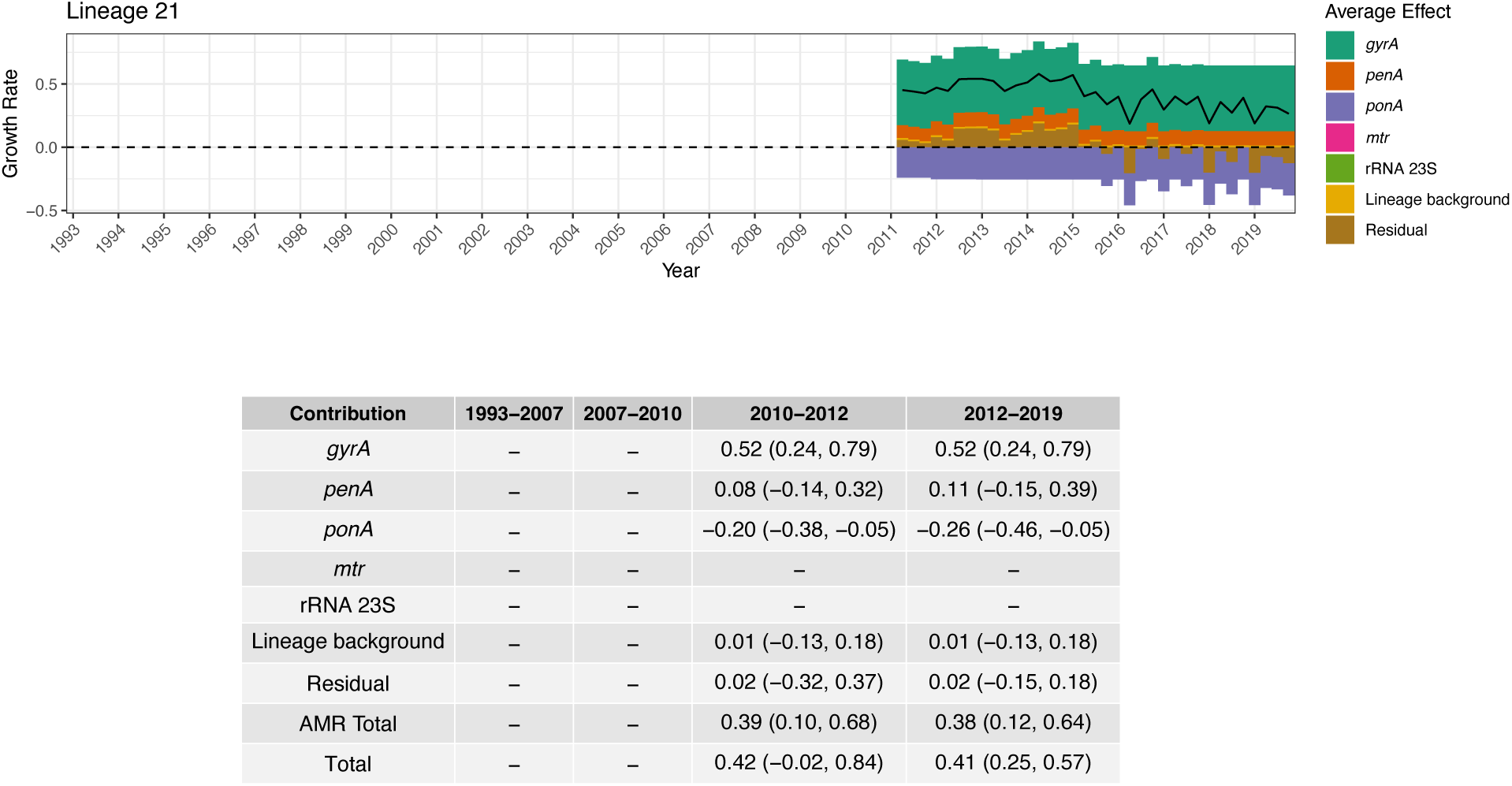
Growth rate effect summary for Lineage 21. The top panel shows the combined average growth rate effect of resistance determinants along with the lineage background term and the residual. The black solid line represents the total average effect. The dashed horizontal line indicates zero. The bottom panel depicts a table summarizing the median total growth rate effect across 4 treatment periods, as well as the 95% credible interval around the median in brackets. The period 1993-2007 corresponds to when fluoroquinolones were recommended as primary treatment; 2007-2010 to when multiple cephalosporins were recommended; 2010-2012 to when multiple cephalosporins were recommended along with azithromycin co-treatment; and 2012-2019 to when only ceftriaxone 250mg along with azithromycin co-treatment was recommended.

**Figure S35:**
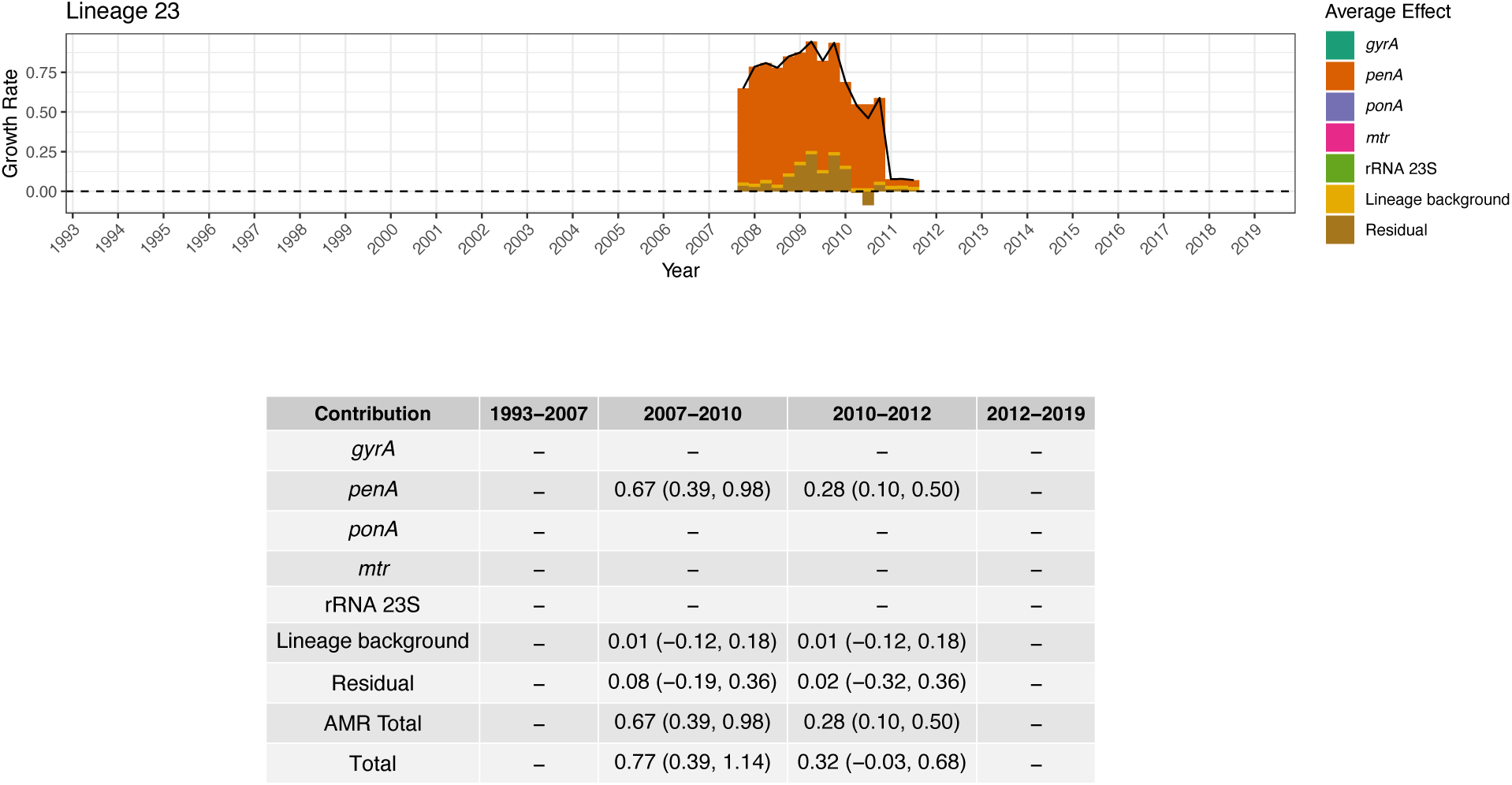
Growth rate effect summary for Lineage 23. The top panel shows the combined average growth rate effect of resistance determinants along with the lineage background term and the residual. The black solid line represents the total average effect. The dashed horizontal line indicates zero. The bottom panel depicts a table summarizing the median total growth rate effect across 4 treatment periods, as well as the 95% credible interval around the median in brackets. The period 1993-2007 corresponds to when fluoroquinolones were recommended as primary treatment; 2007-2010 to when multiple cephalosporins were recommended; 2010-2012 to when multiple cephalosporins were recommended along with azithromycin co-treatment; and 2012-2019 to when only ceftriaxone 250mg along with azithromycin co-treatment was recommended.

**Figure S36:**
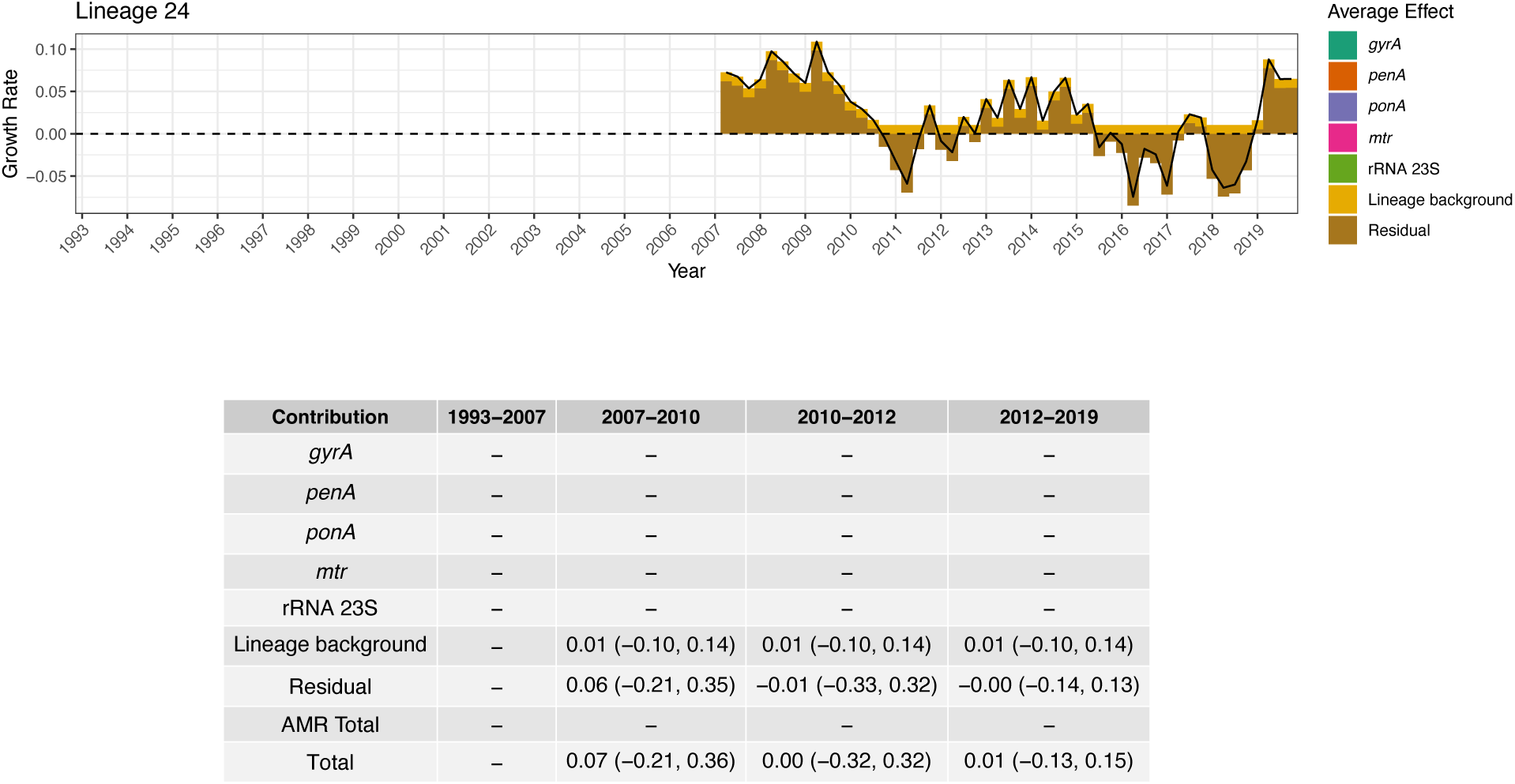
Growth rate effect summary for Lineage 24. The top panel shows the combined average growth rate effect of resistance determinants along with the lineage background term and the residual. The black solid line represents the total average effect. The dashed horizontal line indicates zero. The bottom panel depicts a table summarizing the median total growth rate effect across 4 treatment periods, as well as the 95% credible interval around the median in brackets. The period 1993-2007 corresponds to when fluoroquinolones were recommended as primary treatment; 2007-2010 to when multiple cephalosporins were recommended; 2010-2012 to when multiple cephalosporins were recommended along with azithromycin co-treatment; and 2012-2019 to when only ceftriaxone 250mg along with azithromycin co-treatment was recommended.

**Figure S37:**
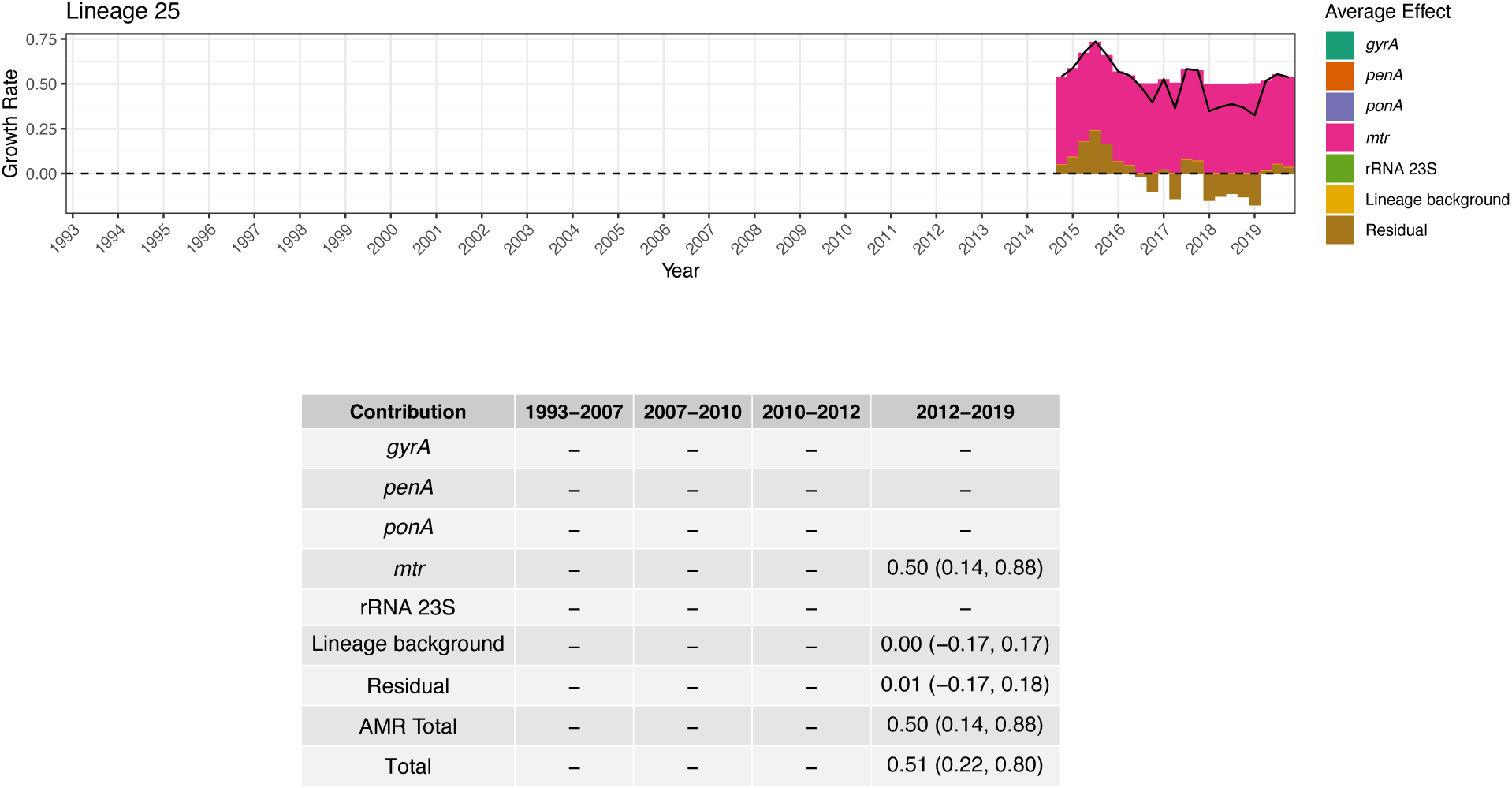
Growth rate effect summary for Lineage 25. The top panel shows the combined average growth rate effect of resistance determinants along with the lineage background term and the residual. The black solid line represents the total average effect. The dashed horizontal line indicates zero. The bottom panel depicts a table summarizing the median total growth rate effect across 4 treatment periods, as well as the 95% credible interval around the median in brackets. The period 1993-2007 corresponds to when fluoroquinolones were recommended as primary treatment; 2007-2010 to when multiple cephalosporins were recommended; 2010-2012 to when multiple cephalosporins were recommended along with azithromycin co-treatment; and 2012-2019 to when only ceftriaxone 250mg along with azithromycin co-treatment was recommended.

**Figure S38:**
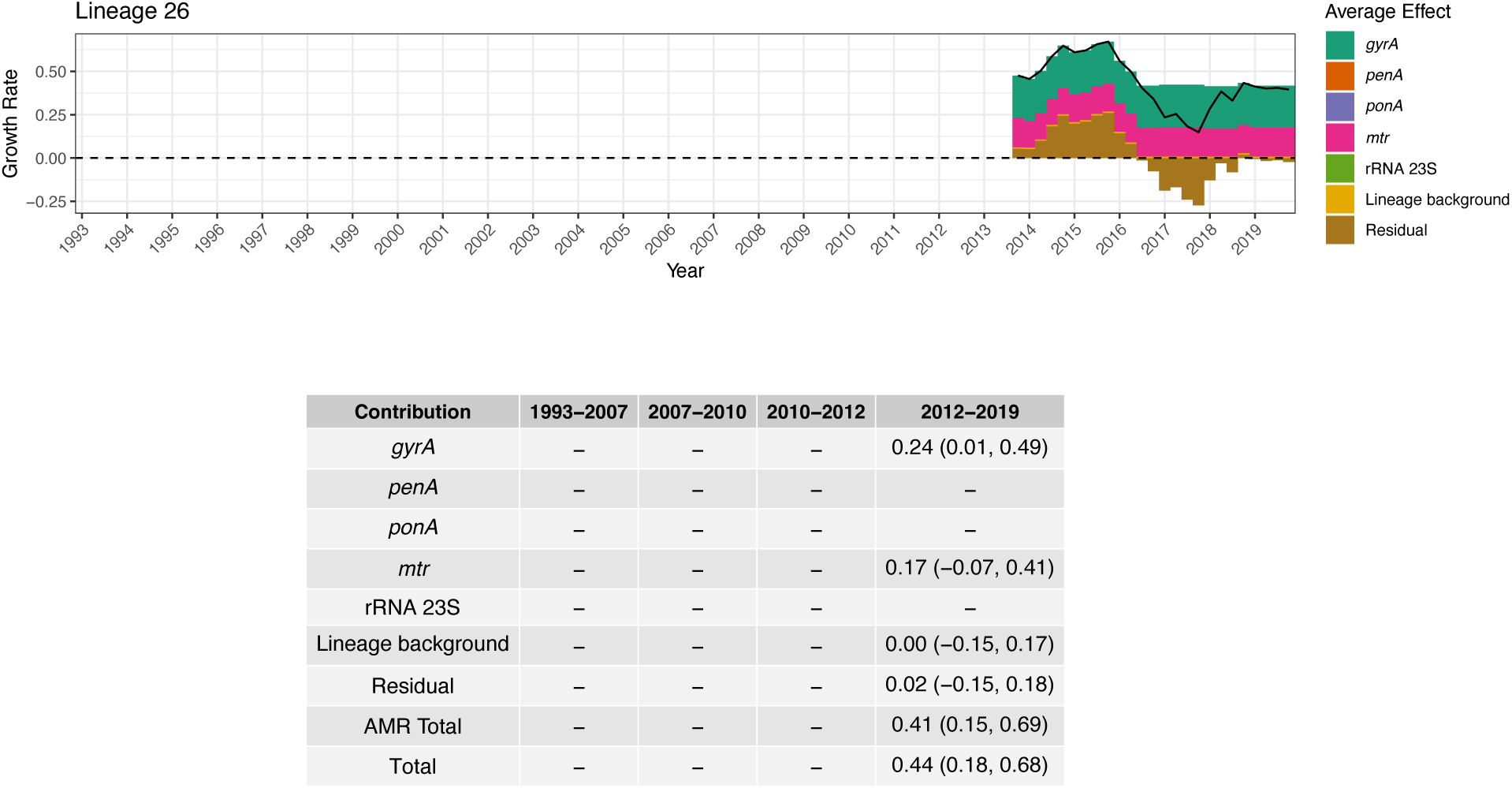
Growth rate effect summary for Lineage 26. The top panel shows the combined average growth rate effect of resistance determinants along with the lineage background term and the residual. The black solid line represents the total average effect. The dashed horizontal line indicates zero. The bottom panel depicts a table summarizing the median total growth rate effect across 4 treatment periods, as well as the 95% credible interval around the median in brackets. The period 1993-2007 corresponds to when fluoroquinolones were recommended as primary treatment; 2007-2010 to when multiple cephalosporins were recommended; 2010-2012 to when multiple cephalosporins were recommended along with azithromycin co-treatment; and 2012-2019 to when only ceftriaxone 250mg along with azithromycin co-treatment was recommended.

**Figure S39:**
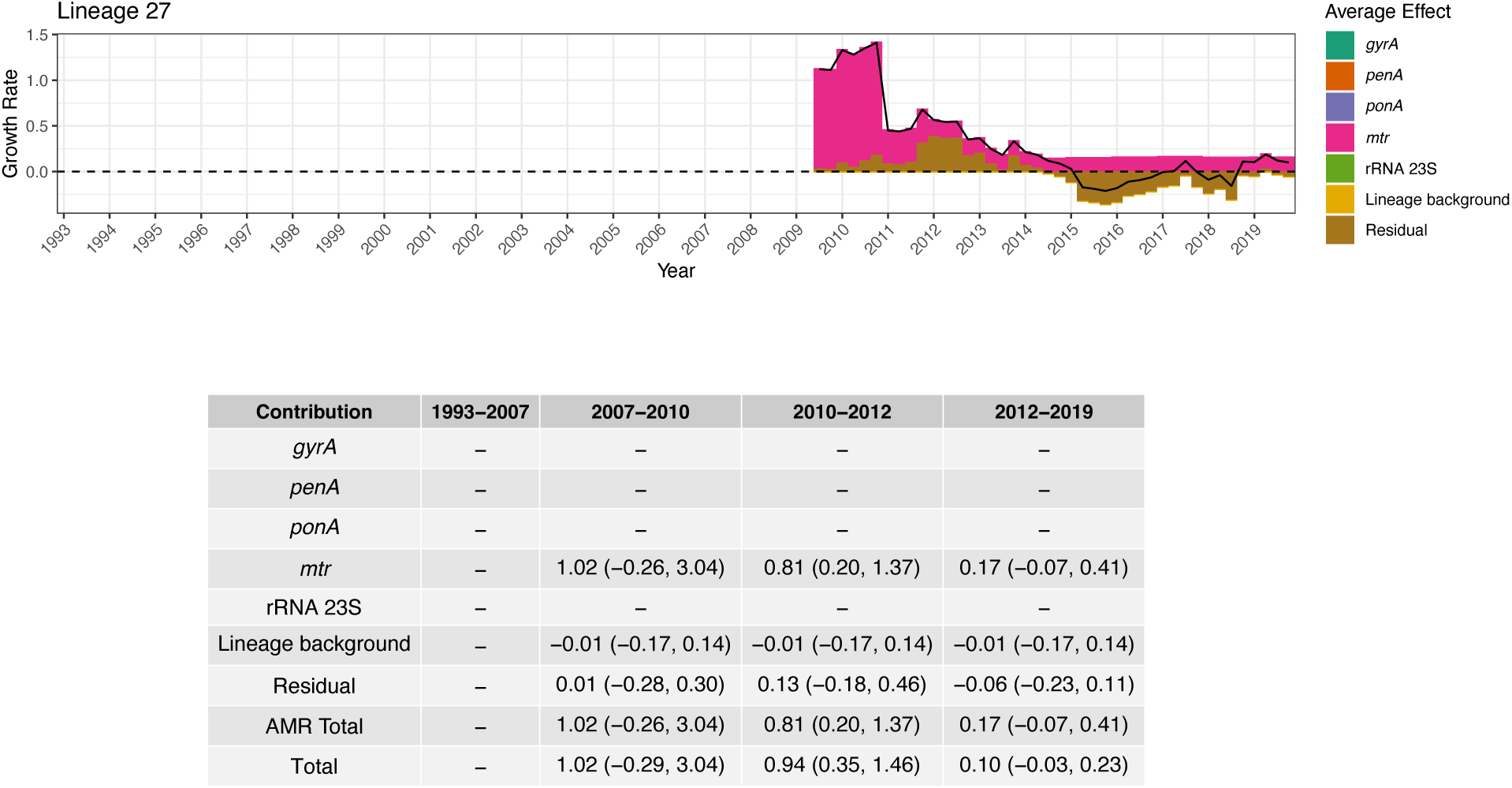
Growth rate effect summary for Lineage 27. The top panel shows the combined average growth rate effect of resistance determinants along with the lineage background term and the residual. The black solid line represents the total average effect. The dashed horizontal line indicates zero. The bottom panel depicts a table summarizing the median total growth rate effect across 4 treatment periods, as well as the 95% credible interval around the median in brackets. The period 1993-2007 corresponds to when fluoroquinolones were recommended as primary treatment; 2007-2010 to when multiple cephalosporins were recommended; 2010-2012 to when multiple cephalosporins were recommended along with azithromycin co-treatment; and 2012-2019 to when only ceftriaxone 250mg along with azithromycin co-treatment was recommended.

**Figure S40:**
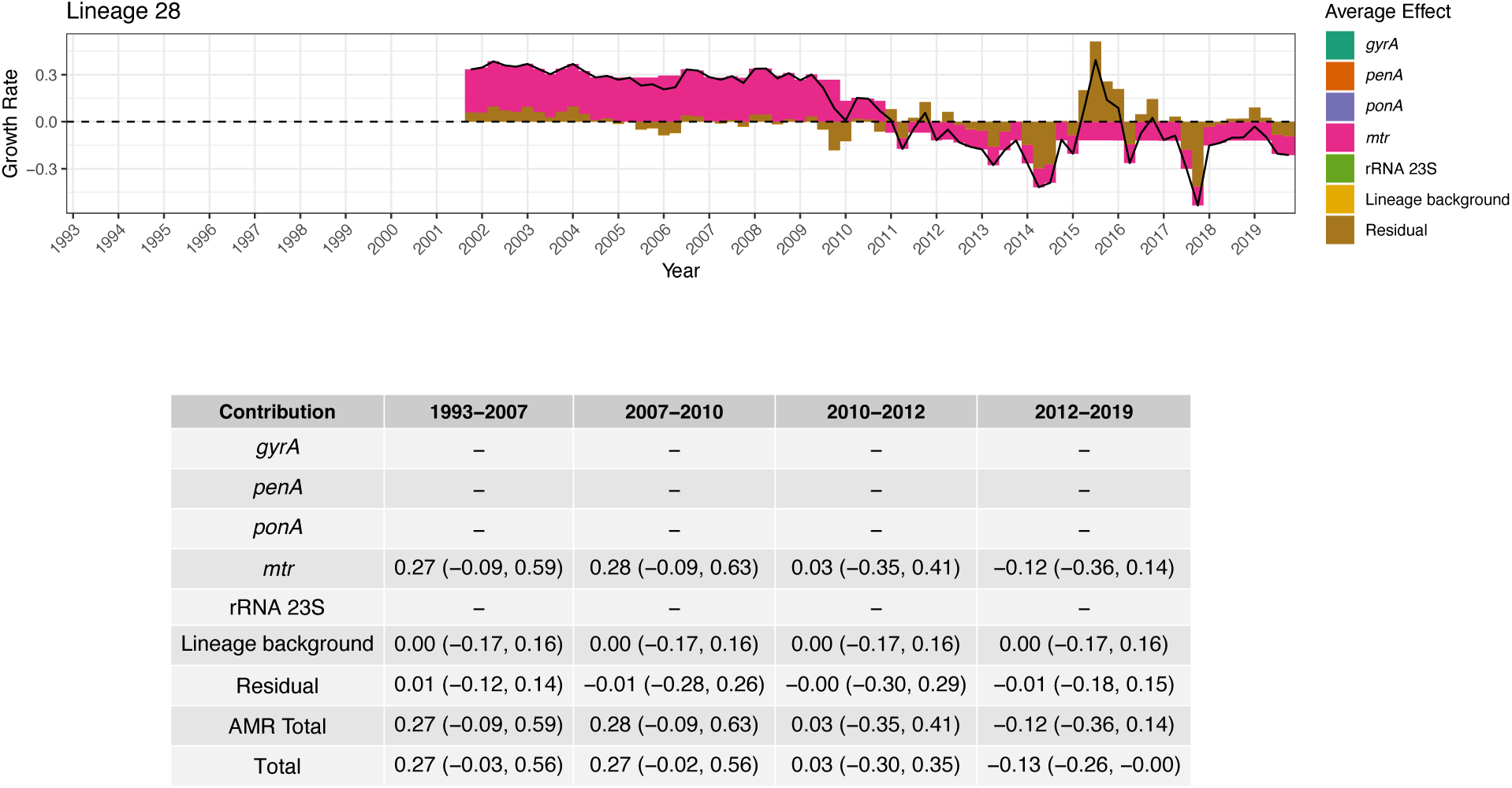
Growth rate effect summary for Lineage 28. The top panel shows the combined average growth rate effect of resistance determinants along with the lineage background term and the residual. The black solid line represents the total average effect. The dashed horizontal line indicates zero. The bottom panel depicts a table summarizing the median total growth rate effect across 4 treatment periods, as well as the 95% credible interval around the median in brackets. The period 1993-2007 corresponds to when fluoroquinolones were recommended as primary treatment; 2007-2010 to when multiple cephalosporins were recommended; 2010-2012 to when multiple cephalosporins were recommended along with azithromycin co-treatment; and 2012-2019 to when only ceftriaxone 250mg along with azithromycin co-treatment was recommended.

**Figure S41:**
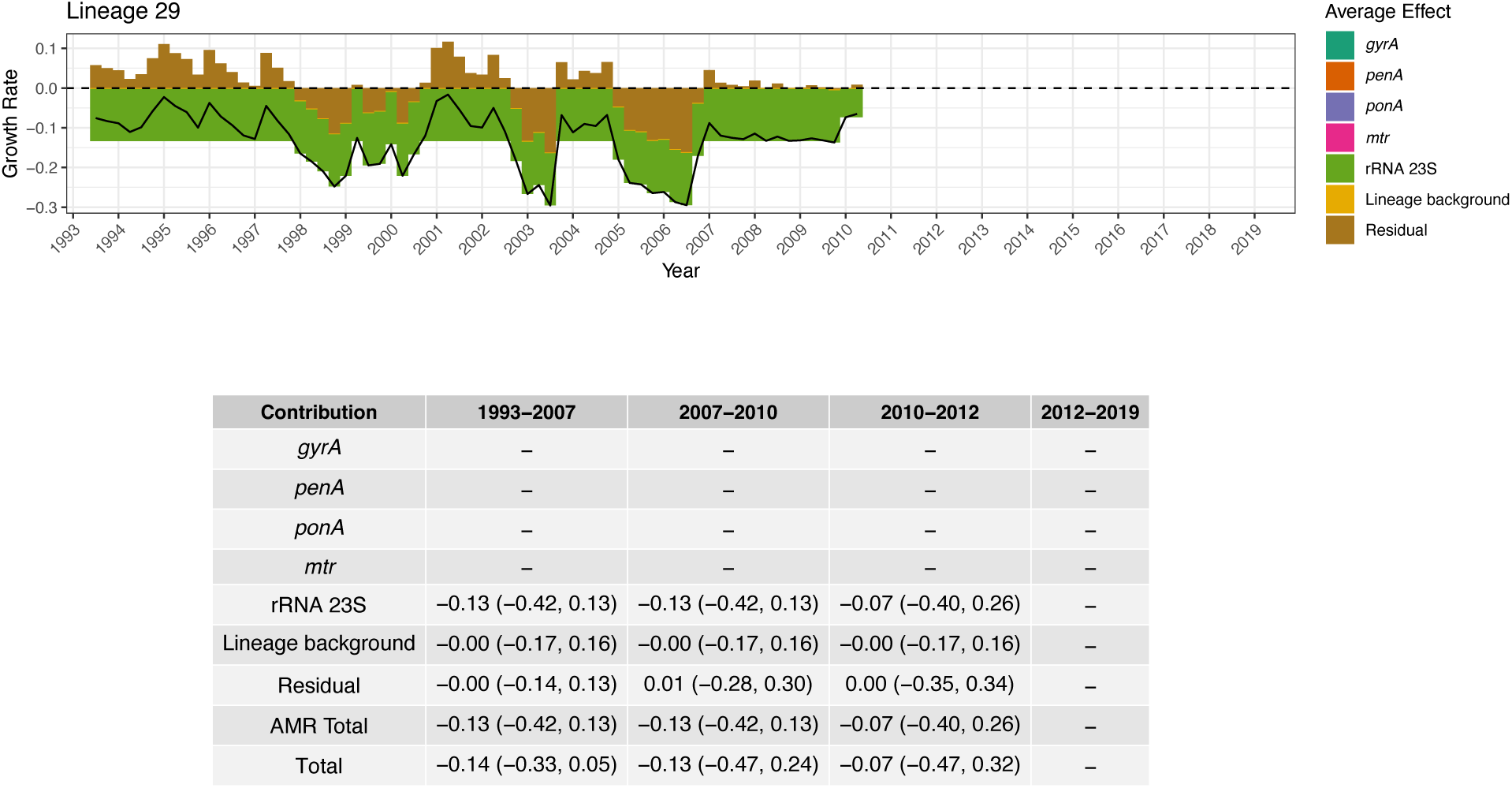
Growth rate effect summary for Lineage 29. The top panel shows the combined average growth rate effect of resistance determinants along with the lineage background term and the residual. The black solid line represents the total average effect. The dashed horizontal line indicates zero. The bottom panel depicts a table summarizing the median total growth rate effect across 4 treatment periods, as well as the 95% credible interval around the median in brackets. The period 1993-2007 corresponds to when fluoroquinolones were recommended as primary treatment; 2007-2010 to when multiple cephalosporins were recommended; 2010-2012 to when multiple cephalosporins were recommended along with azithromycin co-treatment; and 2012-2019 to when only ceftriaxone 250mg along with azithromycin co-treatment was recommended.

**Figure S42:**
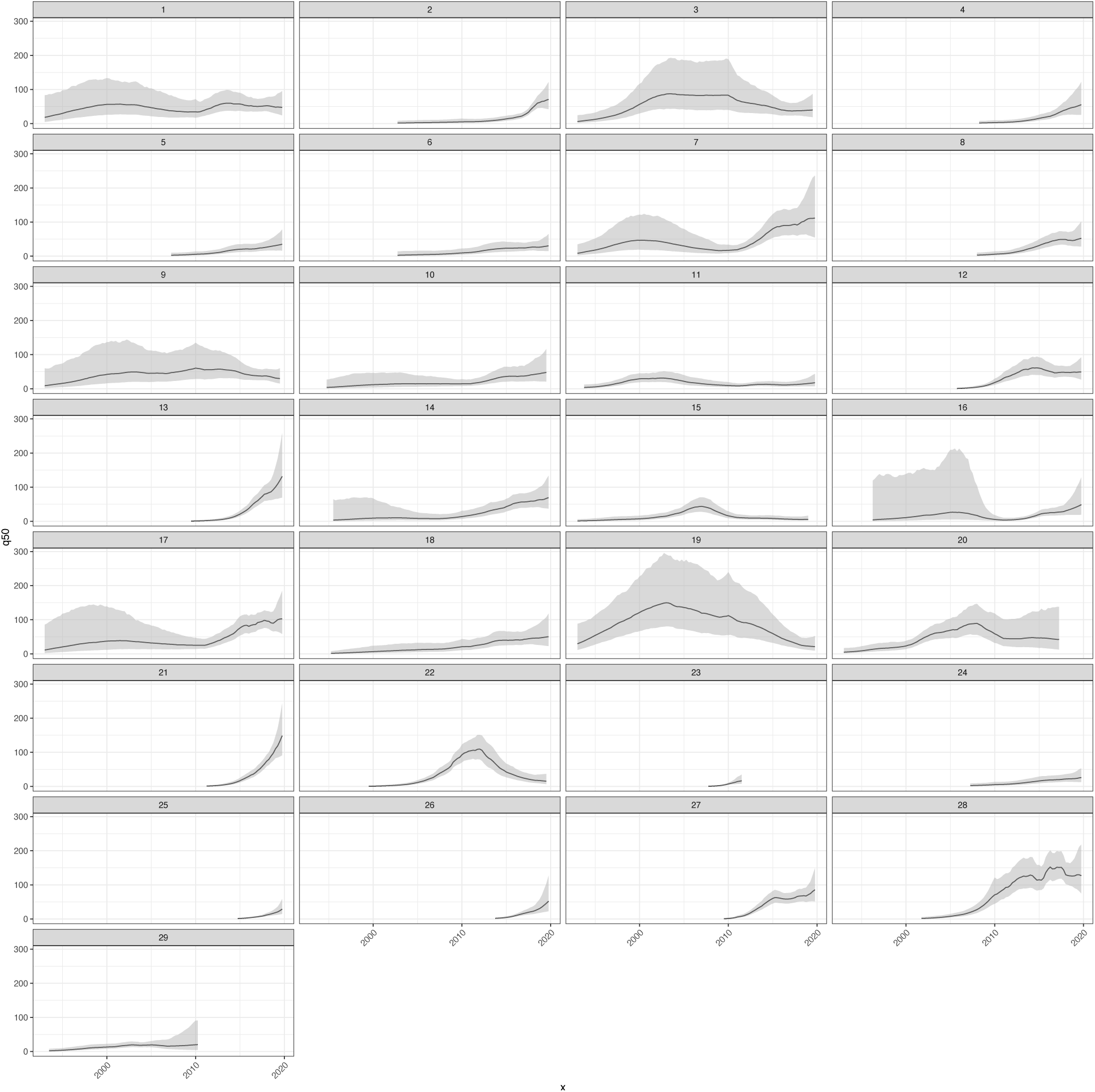
Estimated Ne(t) trajectories for each of the 29 lineages included in the model. The trajectories include the residual term. The trajectories are truncated past the respective lineage MRCA.

**Figure S43:**
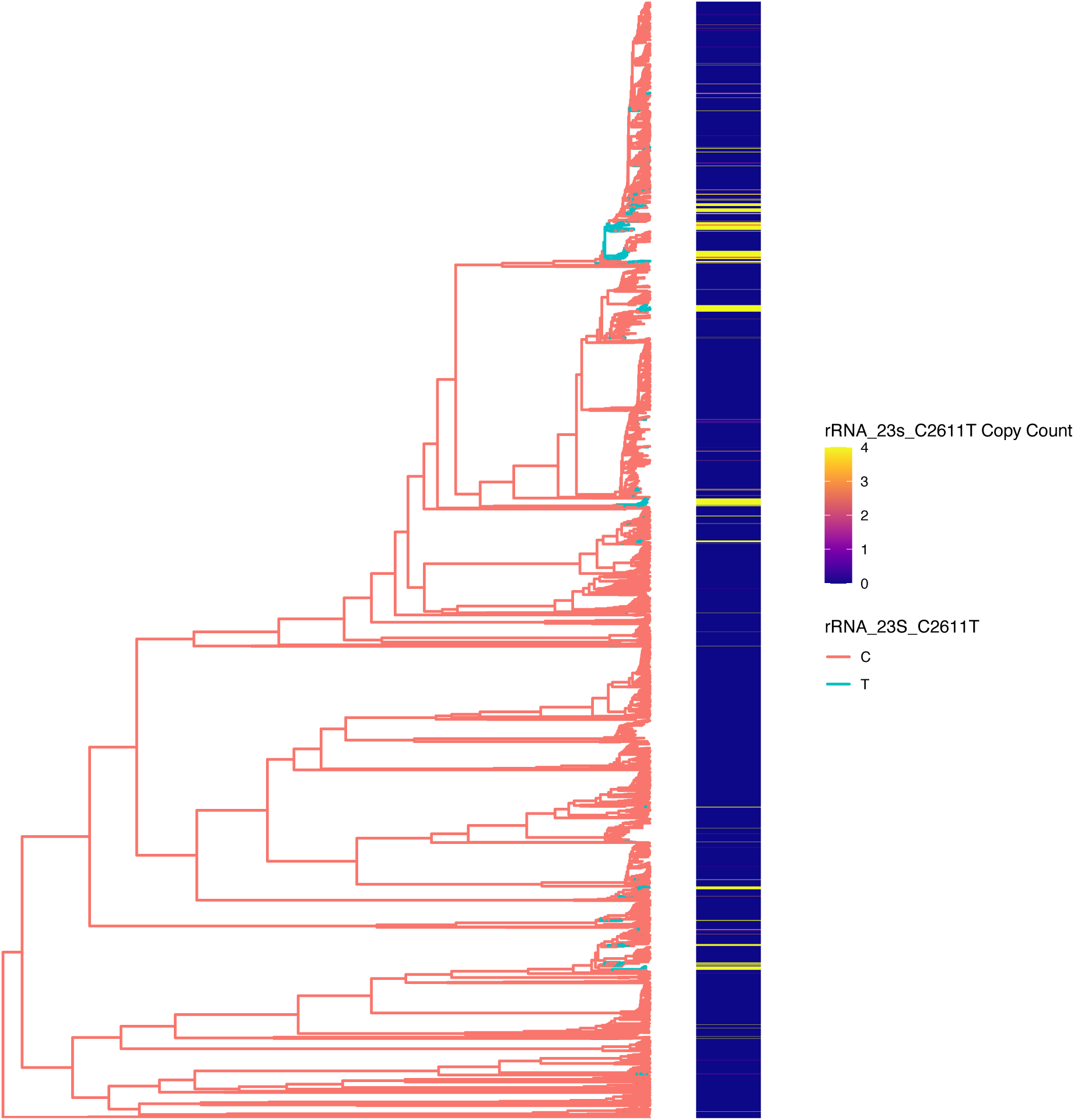
Distribution of 23S rRNA C2611T across the phylogenetic tree. The tree coloring corresponds to a DELTRAN based ancestral state reconstruction.

**Figure S44:**
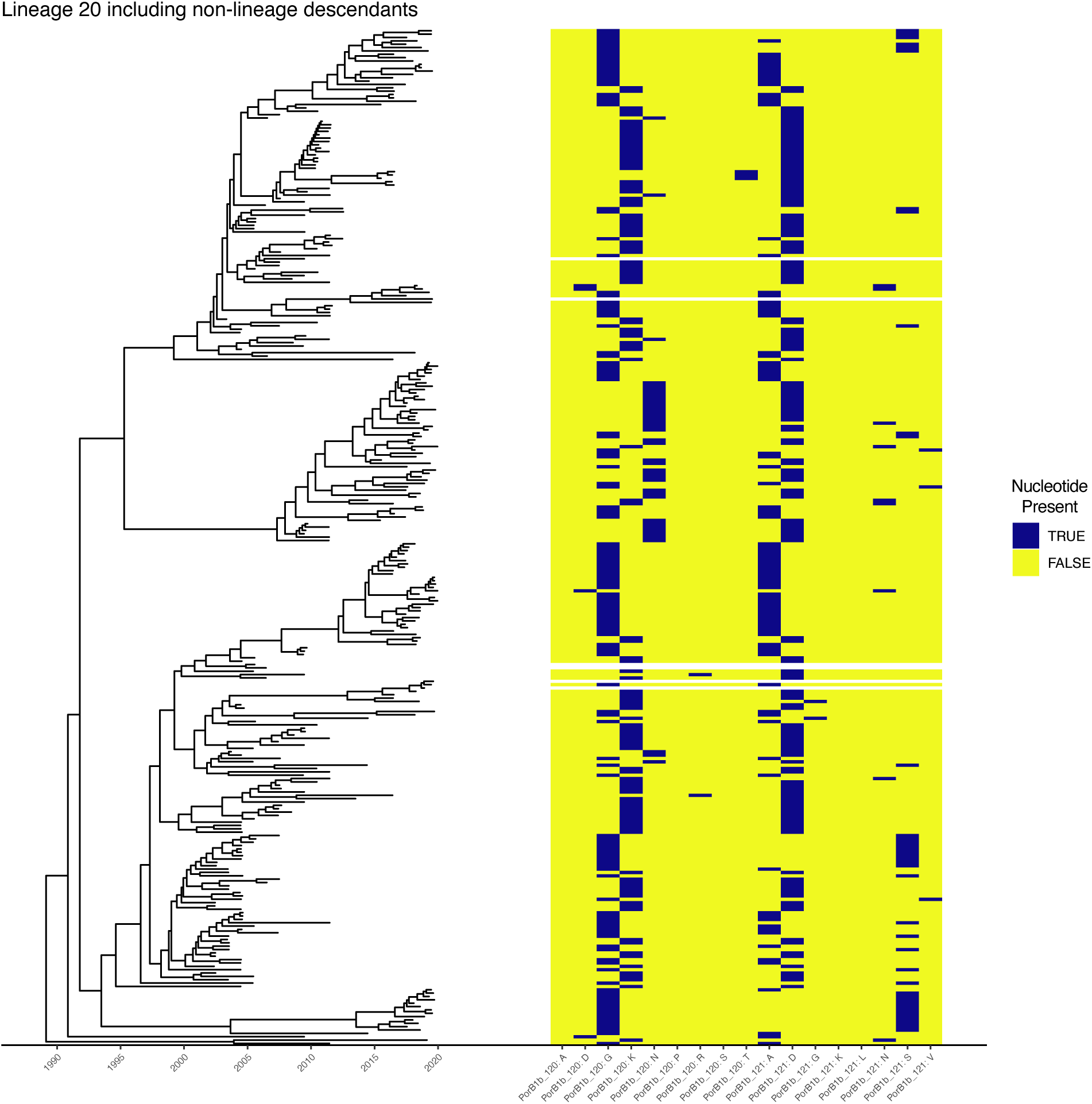
The distribution of PorB G120 and A121 polymorphisms in lineage 20. Note the high rate of loss and gain of polymorphisms and lack of clonal inheritance.

**Table S1:** Interactions between individual loci and antibiotics classes as coded in the regression model used.

| Locus | Penicillin | Fluoroquinolones | Cephalosporins other than Ceftriaxone 250mg | Azithromycin |
| --- | --- | --- | --- | --- |
| *gyrA* | No | Yes | No | No |
| *ponA* | Yes | No | Yes | No |
| *penA* | Yes | No | Yes | No |
| *mtr* | Yes | Yes | Yes | Yes |
| 23S rRNA | No | No | No | Yes |

**Table S2:**
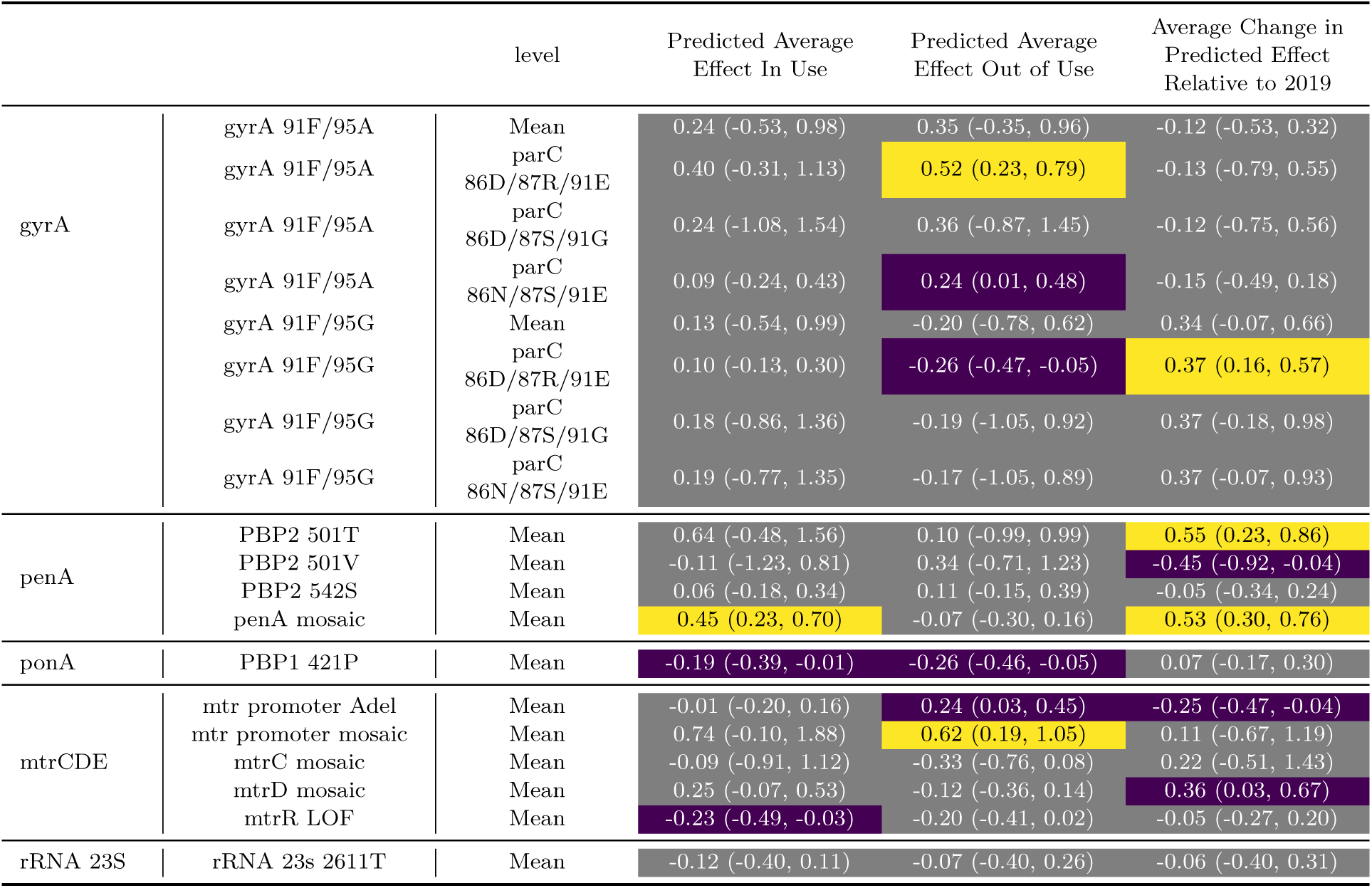
Predicted effect summaries (Posterior median and in brackets 95% posterior credible interval) for all the determinants included in the model. The predicted effect summary is computed as the given effect averaged over either the in-use time period or the out-of-use time period. The average change in relative effect is computed as the change in effect relative to 2019 averaged over the in-use period. For *gyrA* the in-use period covers 1993-2006, and the out-of-use period covers 2007 onward. For *penA*, *ponA*, *mtrCDE*, and rRNA 23S the in-use period covers 1993-2011, and the out-of-use period covers 2012 onwards. Yellow color corresponds to estimates where the 95% posterior credible interval excludes the region of [–0.1,0.1], and blue color corresponds to estimates where the 95% posterior credible interval excludes 0.

**Table S3:** Ciprofloxacin Minimum Inhibitory Concentrations (MICs) of *N. gonorrhoeae* isogenic strains.

| Strains | Ciprofloxacin MIC ($\mu\text{g/mL}$) |
| --- | --- |
| GCGS0481 GyrA 91S/95G (Parent) | $\geq 32$ |
| GCGS0481 GyrA 91S/95D, CMR | 0.25 |
| GCGS0481 GyrA 91F/95G, CMR | $\geq 32$ |
| GCGS0481 GyrA 91F/95A, CMR | $\geq 32$ |

**Table S4:**
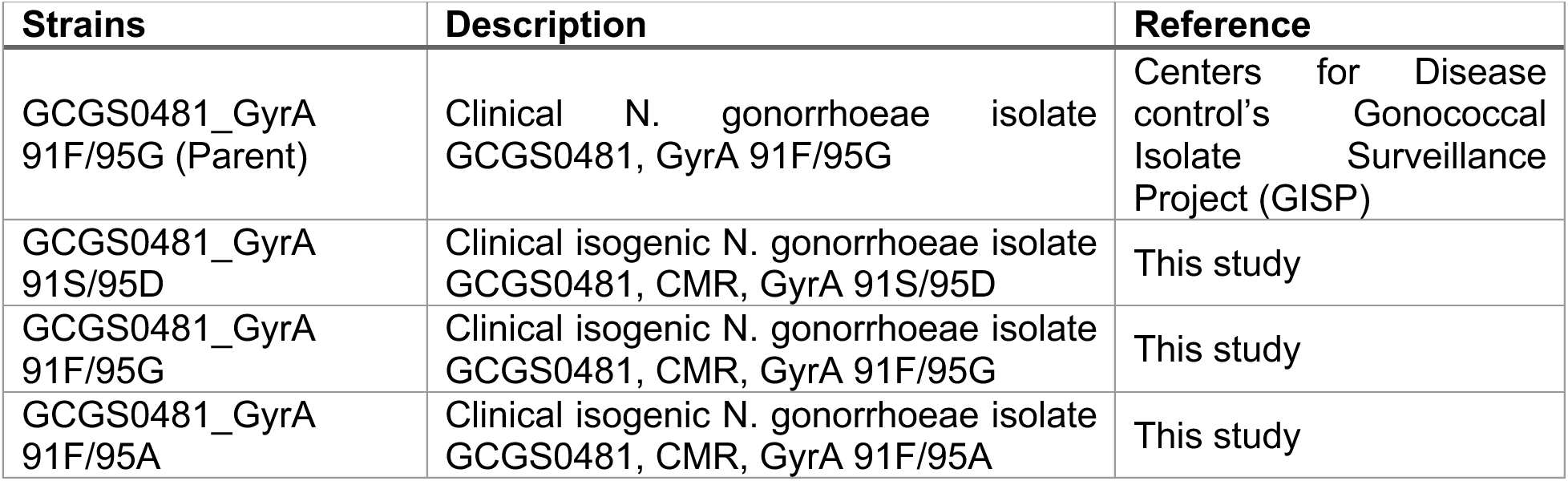
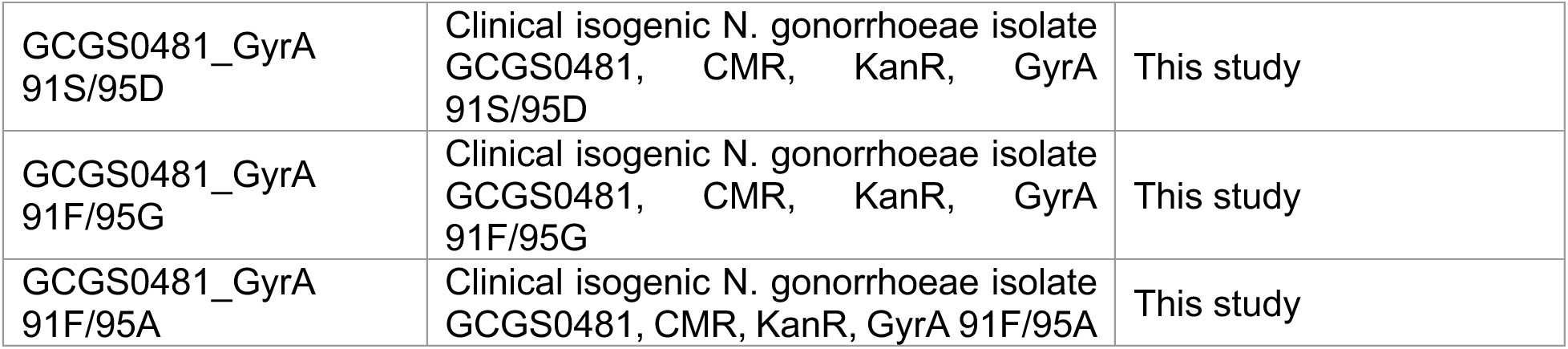
*N. gonorrhoeae* strains used in this study for GyrA mutant competition experiments.

| Strains | Description | Reference |
| --- | --- | --- |
| GCGS0481_GyrA 91F/95G (Parent) | Clinical *N. gonorrhoeae* isolate GCGS0481, GyrA 91F/95G | Centers for Disease control's Gonococcal Isolate Surveillance Project (GISP) |
| GCGS0481_GyrA 91S/95D | Clinical isogenic *N. gonorrhoeae* isolate GCGS0481, CMR, GyrA 91S/95D | This study |
| GCGS0481_GyrA 91F/95G | Clinical isogenic *N. gonorrhoeae* isolate GCGS0481, CMR, GyrA 91F/95G | This study |
| GCGS0481_GyrA 91F/95A | Clinical isogenic *N. gonorrhoeae* isolate GCGS0481, CMR, GyrA 91F/95A | This study |
| GCGS0481_GyrA 91S/95D | Clinical isogenic *N. gonorrhoeae* isolate GCGS0481, CMR, KanR, GyrA 91S/95D | This study |
| GCGS0481_GyrA 91F/95G | Clinical isogenic *N. gonorrhoeae* isolate GCGS0481, CMR, KanR, GyrA 91F/95G | This study |
| GCGS0481_GyrA 91F/95A | Clinical isogenic *N. gonorrhoeae* isolate GCGS0481, CMR, KanR, GyrA 91F/95A | This study |

**Table S5:** Plasmids used in this study for GyrA mutant competition experiments.

| Plasmids | Description | Reference |
| --- | --- | --- |
| pDRE77 | pUC19 with AphA3 Kan<sup>R</sup> cassette + homology to clone GyrA 91S/95D | (77) |
| pKH37 | CM<sup>R</sup>, lac-inducible, inserts into lctP/aspC site of *N. gonorrhoeae* | (78) |
| pAM_3 | pUC19 with CM<sup>R</sup> cassette + homology to clone GyrA 91S/95D | This study |
| pDR1 | KanR derivative of pKH37 | (79) |
| pDR53 | A derivative of pDR1 without Lac promoter and the regulatory elements + added multi cloning site (MCS) | This study |

**Table S6:**
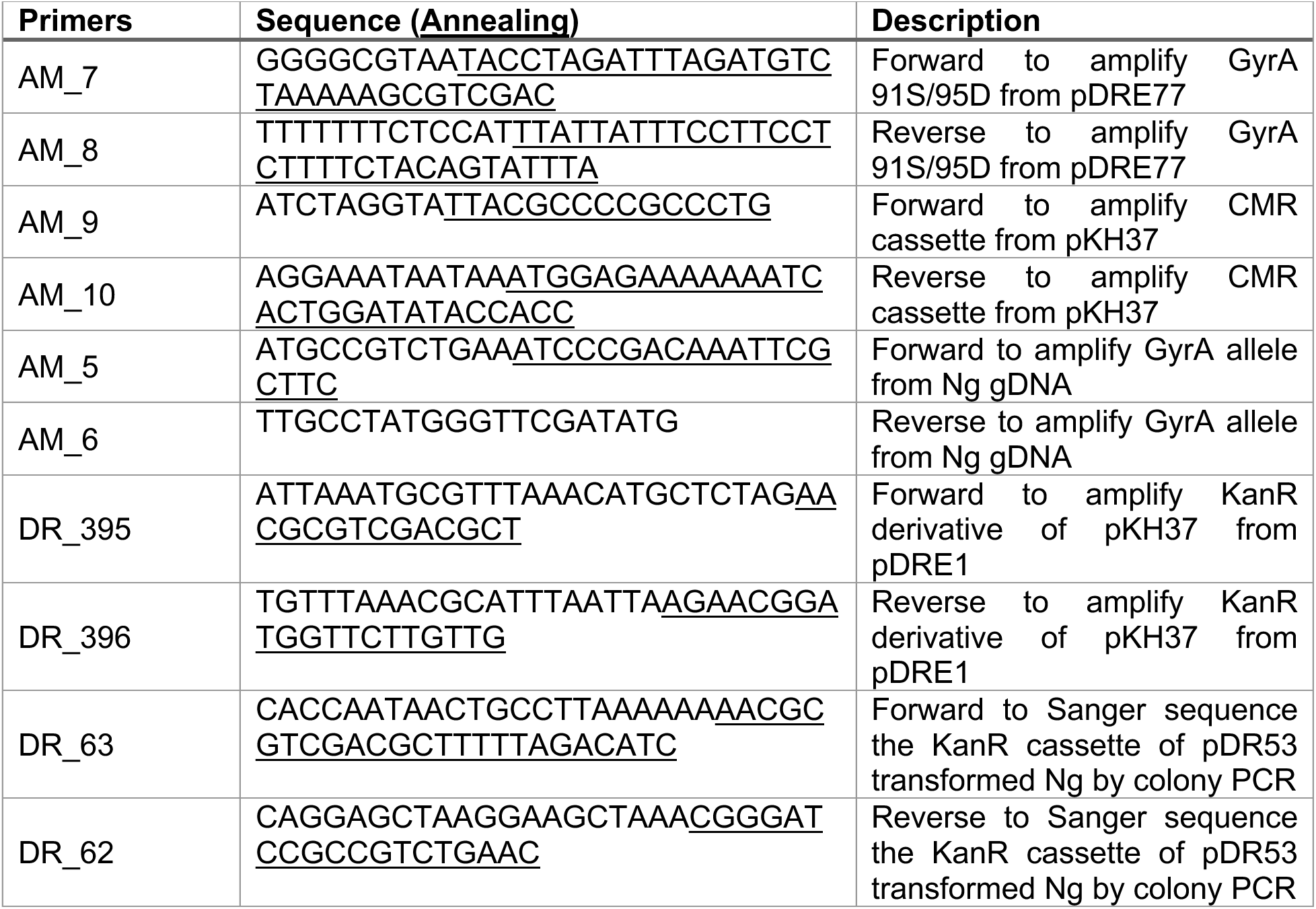
Primers used in this study for GyrA mutants competition experiments.

**Table S7:** *N. gonorrhoeae* strains used in this study for *ponA* mutant competition experiments. Transformants are shown as recipient strain x donor DNA, produced as described in Materials and Methods. PCR signifies a PCR product was used to transform, and pUC18 denotes all cloning plasmids used.

| Strains | Description | Reference |
| --- | --- | --- |
| FA19 | Wild type, antibiotic-susceptible lab strain | (82) |
| FA6140 | Penicillin-resistant clinical isolate | (34) |
| FA19 *rpsL* PBP1<sup>421L</sup> | FA19 x *rpsL*1 PCR; naturally expresses PBP1 421L | (84) |
| FA6140 *rpsL* PBP1<sup>421P</sup> | FA6140 x *rpsL*1 PCR; naturally expresses PBP1 421P | This study |
| FA19 *rpsL* PBP1<sup>421P</sup> | FA19 *rpsL* x pUC18us-PBP1<sup>421P</sup>-Ω; replaces PBP1 421L with PBP1 421P | This study |
| FA6140 *rpsL* PBP1<sup>421L</sup> | FA6140 *rpsL* x pUC18us-PBP1<sup>421L</sup>-Ω; replaces PBP1 421P with PBP1 421L | This study |

**Table S8:** Plasmids used in this study for *ponA* mutant competition experiments.

| Plasmids | Description | Reference |
| --- | --- | --- |
| pUC18us-PBP1<sup>421P</sup>-Ω | pUC18-us containing bp 831-2400 of *ponA*1, 54 bp downstream of *ponA*, the *aad1* resistance cassette ($\Omega$), and 531 bp of additional downstream sequence | This study |
| pUC18us-PBP1<sup>421L</sup>-Ω | pUC18-us containing bp 831-2400 of *ponA*, 54 bp downstream of *ponA*, the *aad1* resistance cassette ($\Omega$), and 531 bp of additional downstream sequence | This study |

**Table S9:**
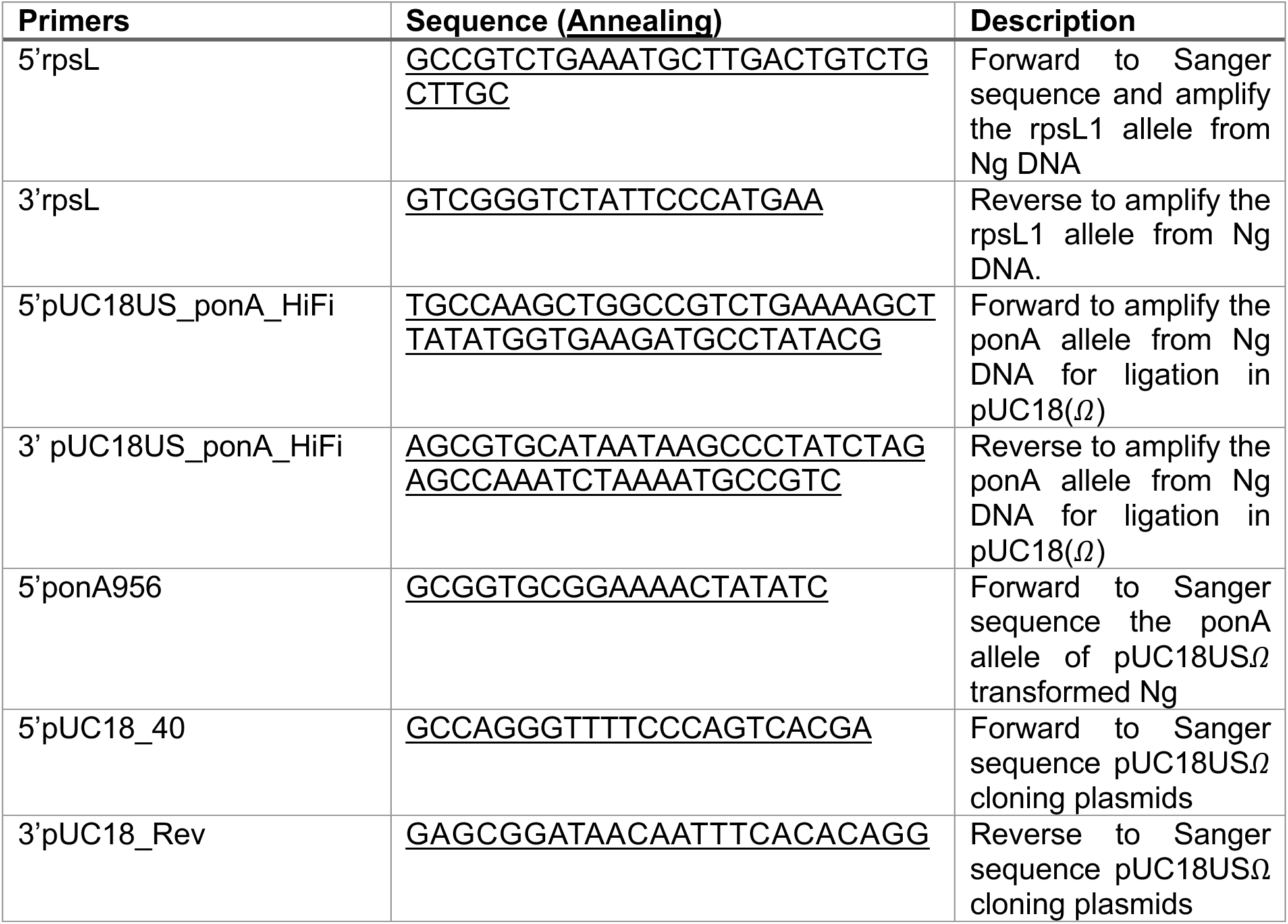
Primers used in this study for *ponA* mutant competition experiments.

## Notes

### Competing Interest Statement

The authors have declared no competing interest.

### Summary of Updates

Fix pdf title to match the biorxiv title, fix broken hyperlinks within the PDF document.

